# The genetic architecture of fruit colour in strawberry (*Fragaria × ananassa*) uncovers the predominant contribution of the *F. vesca* subgenome to anthocyanins and reveals underlying genetic variations

**DOI:** 10.1101/2020.06.25.171496

**Authors:** Marc Labadie, Guillaume Vallin, Aurélie Petit, Ludwig Ring, Thomas Hoffmann, Amèlia Gaston, Aline Potier, Juan Munoz-Blanco, José L. Caballero, Wilfried Schwab, Christophe Rothan, Béatrice Denoyes

## Abstract

Fruit colour, which is central to the sensorial and nutritional quality of the fruit, is a major breeding target in cultivated strawberry (*Fragaria × ananassa*). Red fruit colour is caused by the accumulation of anthocyanins, which are water-soluble flavonoids. Here, using pseudo F_1_ progeny derived from the cv. ‘Capitola’ and the advanced line ‘CF1116’ and taking advantage of the available high density SNP array, we delineated fruit flavonoids mQTLs (13 compounds: anthocyanin, flavonols and flavan-3-ols) and colour-related QTLs (total anthocyanins and fruit colour) to narrow genomic regions corresponding to specific subgenomes of the cultivated strawberry. Findings showed that the overwhelming majority of the anthocyanin mQTLs and other colour-related QTLs are specific to *F. vesca* subgenome but that other subgenomes also contribute to colour variations. We then focused on two major homoeo-mQTLs for pelargonidin-3-glucoside (PgGs) localized on both male and female maps on linkage group LG3a (*F. vesca* subgenome). Combined high-resolution mapping of PgGs mQTLs and colour QTLs, transcriptome analysis of selected progeny individuals and whole genome sequencing of the parents led to the identification of several INDELS in the *cis*-regulatory region of a *MYB102-like ODORANT* gene and of the deletion of a putative MADS box binding motif in the 5’UTR upstream region of an anthocyanidin reductase (*ANR)* gene, which likely underlie significant colour variations in strawberry fruit. The implications of these findings are important for the functional analysis and genetic engineering of colour-related genes and for the breeding by Marker-Assisted-Selection of new strawberry varieties with improved colour and health-benefits.

## Introduction

Cultivated strawberry (*Fragaria × ananassa*) is the most consumed small fruit worldwide. Since its creation in the 18^th^ century in botanical gardens in Europe by fortuitous hybridization between the two New World strawberry species *F. chiloensis* and *F. virginiana* (Edger et al. 2019), cultivated strawberry has been continuously improved to fit the needs of both producers and consumers. Until recently, strawberry improvement was primarily focused on features of concern to producers including plant and fruit yield, resistance to strawberry diseases and adaptation to cultural practices. In the last decades, Quantitative Trait Loci (QTLs) controlling several related traits, among which early or late fruit harvesting, extension of the production period, modulation of the runnering/flowering trade-off (Perrotte et al. 2016; Gaston et al. 2020), and resistance to anthracnose and other diseases (Cockerton et al. 2018; Salinas et al. 2019) have been identified, thereby enabling the design of genetic markers for strawberry improvement by Marker-Assisted Selection (MAS) (Denoyes et al. 2017).

In the recent years, fruit sensorial quality has also become a major target for strawberry breeding (Mezzetti et al. 2018). Among the sensorial traits to improve is the colour of the fruit. Fruit colour plays an important role in the attractiveness of the strawberry and is due to the accumulation of the red pigments anthocyanins, which are water-soluble flavonoids. The flavonoids detected in strawberry fruit (anthocyanins, flavonols, flavan-3-ols) (Ring et al., 2013; Urrutia et al. 2016; Labadie et al. 2020) are derived from the phenylpropanoid pathway (Figure 1A). Flavonols and flavan-3-ols are mainly glycosides of quercetin and kaempferol as well as derivatives of catechin and epicatechin. Anthocyanins are mainly glycosides of pelargonidin and cyanidin (Hanhineva et al. 2011), the composition of which gives to the fruit its distinctive colour hue, from bright red (pelargonidin derivatives) to dark red (cyanidin derivatives). In addition, the anthocyanins are antioxidant molecules with proven dietary health-benefits (Tulipani et al. 2008, 2009; Winter et al. 2018). They therefore contribute to the nutritional quality of strawberries (Battino et al. 2009; Giampieri et al. 2014, 2017; Gasparrini et al. 2018; Miller et al. 2019), which consumers are increasingly taking into account. Studies in mice fed with engineered purple tomato accumulating high anthocyanin content or with anthocyanin-enriched extract from strawberry established unambiguously the health protecting effect of flavonoids (Butelli et al. 2008; Winter et al. 2018). Furthermore, different classes of polyphenols including anthocyanins, flavonols and stilbene were recently shown to act synergistically in protecting against inflammatory gut disease in mice (Scarano et al. 2018). All this information points to the need for the discovery of genetic variations underlying the fruit colour traits in strawberry, so that MAS can be applied to improve the sensorial and nutritional value of strawberry.

**Figure 1.**
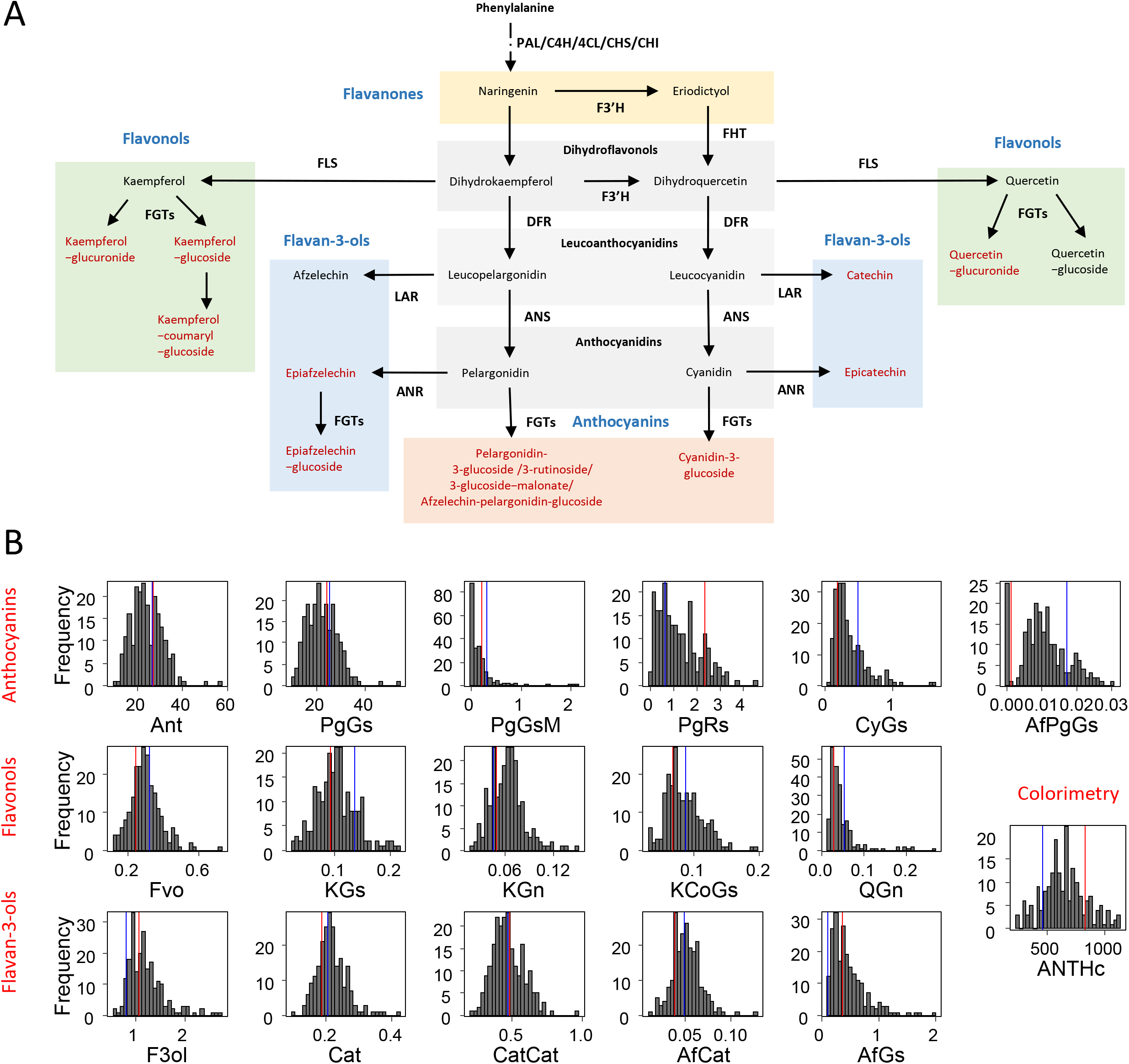
Flavonoids of strawberry (*Fragaria × ananassa*) fruit and their distribution in the progeny. **A)** Simplified flavonoid biosynthetic pathway. Compounds assessed in this study are in red. Chemical classes are in blue. ANR, anthocyanidin reductase; ANS, anthocyanidin synthase; CHI, chalcone isomerase; C4H, cinnamic acid-4-hydroxylase; 4CL, 4-coumarate:CoA ligase; CHS, chalcone synthase; DFR, dihydroflavonol-4-reductase; FGT, UDPglucose: flavonoid-3-O-glucosyltransferase; FHT/F3H, flavanone-3-hydroxylase; FLS, flavonol synthase; LAR, leucoanthocyanidin reductase; PAL, phenylalanine ammonia-lyase. **B**) Distribution of the progeny mean in 2010. Mean phenotypic values from parents are represented in red for ‘Capitola’ and in blue for ‘CF1116’. Ant, Fvo, F3ol values were obtained by summation of total anthocyanins, total flavonols and total flavan-3-ols, respectively; PgGs, Pelargonidin-3-glucoside; PgGsM, Pelargonidin-3-glucoside-malonate; PgRs, Pelargonidin-3-rutinoside; CyGs, Cyanidin-3-glucoside; AfPgGs, (epi)Afzelechin-pelargonidin-3-glucoside; KGs, Kaempferol-glucoside; KGn, Kaempferol-glucuronide; KCoGs, Kaempferol-coumaryl-glucoside; QGn, Quercetin-glucuronide; Cat, Catechin; CatCat, (epi)Catechin dimers; AfCat, (epi)Afzelechin-(epi)catechin dimers; AfGs, (epi)Afzelechin-glucoside; ANTHc, anthocyanins (colourimetry). The flavonoid metabolites values are expressed as mg-equ/100 g FW assuming a response factor of 1. ANTHc results are expressed as mg pelargonidin-3-glucoside equivalents/100 g FW.

In a previous study aimed at uncovering the genetic architecture of fruit sensorial traits in cultivated strawberry, we studied a pseudo full-sibling F_1_ progeny and identified several fruit colour QTLs by measuring total anthocyanins with colourimetric assay and by assessing fruit colour through physical parameters (Lerceteau-Köhler et al. 2012). To get more insights into the genetic control of fruit flavonoids and colour, we then used LC-ESI-MS (Ring et al. 2013; Haugeneder et al. 2018) to identify individual flavonoids in this progeny and mapped the corresponding metabolic QTLs (mQTLs) onto strawberry genetic map using SSR markers (Labadie et al. 2020). This study, which considerably advanced our insights into the genetic architecture of flavonoids in strawberry, hitherto little-known (Lerceteau-Köhler et al. 2012; Urrutia et al. 2016), also enabled us to identify genetic markers for breeding strawberry varieties with improved fruit colour, sweetness and acidity by MAS (Lerceteau-Köhler et al. 2012; Labadie et al. 2020). However, identifying the genetic variants underlying fruit colour variations requires much higher map resolution than that provided by SSR, SSCP and AFLP markers as previously done (Labadie et al. 2020). The octoploid status of cultivated strawberry (2n = 8x = 56) renders this task even more difficult; at a single locus, up to eight homoeoalleles located on four linkage groups (LG) corresponding to the four subgenomes of *F. x ananassa* may control the content in a given anthocyanin compound. Advances in strawberry genomics recently provided the tools necessary to achieve this task. The unprecedented resolution offered by the high-density strawberry SNP genotyping array (Bassil et al., 2015) now permits to narrow down a given mQTL to a small chromosomal region on a specific LG. In addition, the mapping of genomic sequences of selected genotypes onto the recently released high quality reference genome from cultivated strawberry (Edger et al. 2019) should now allow the identification of homoeoallelic variants harbored by specific subgenomes. Combination of both will likely accelerate the discovery of genetic variants underlying trait variations in strawberry.

In this study, using a cross between cv. ‘Capitola’ and the advanced line ‘CF1116’ of cultivated strawberry (Lerceteau-Khöler et al. 2012), we first mapped with SNP markers the flavonoid mQTLs and the colour-related QTLs previously identified over a two-year study (Labadie et al. 2020). Findings showed that the overwhelming majority of the anthocyanin mQTLs and the fruit colour QTLs are harboured by the *F. vesca* subgenome. We then focused on two major homoeo-mQTLs for pelargonidin-3-glucoside (PgGs) that were localized on linkage group LG3a (*F. vesca* subgenome) on both male and female maps in 2010 and 2011. Combination of PgGs mQTL mapping, transcriptome analysis of progeny individuals and whole genome sequencing of the parents led to the identification of homoeoallelic specific variations in the upstream *cis*-regulatory region of anthocyanidin reductase (*ANR)* gene and in the coding region of a *MYB* gene, both of which may contribute to the increased accumulation of PgGs and therefore to the enhanced colour of strawberry fruit. These discoveries are important to further our knowledge of the genes controlling anthocyanin biosynthesis in the fruit and to define the best strategy for breeding new strawberry varieties with improved colour and health benefits.

## Results

### Mapping of fruit colour-related traits in strawberry

In order to delineate genomic regions responsible for flavonoid and colour variations in the pseudo full-sibling F1 progeny derived from a cross between ‘Capitola’ X ‘CF1116’ (72 individuals in 2010 and 131 individuals in 2011), we first analyzed by LC-ESI-MS thirteen fruit flavonoid compounds belonging to three chemical classes (Fig. 1A). As described in Labadie et al. (2020), the flavonoids detected in the fruit were the flavonols (4 compounds), the flavan-3-ols (4 compounds) and the anthocyanins (5 compounds: pelargonidin-3-glucoside (PgGs), pelargonidin-3-glucoside-malonate (PgGsM), pelargonidin-3-rutinoside (PgRs), cyanidin-3-glucoside (CyGs), and (epi)afzelechin-pelargonidin-glucoside (EpPgGs)). We also measured the total anthocyanin content (ANTHc) by colourimetry. We additionally scored the fruit colour in 2011 by comparison with a strawberry colour chart (COLOUR trait). For all the traits, except COLOUR, the trait quantitative values and range in parents and progeny, the broad sense heritability and the transgression values in the ‘Capitola’ X ‘CF1116’ population were reported in Labadie et al. (2020). The COLOUR values are reported in Supplemental Table 1 and Supplemental Fig. 1. Distributions of flavonoids and colour-related traits are represented in Figure 1B for 2010 and in Supplemental Fig. 2 for 2011. Continuous variations of the phenotypic values were observed in the progeny for all the traits assessed in 2010 and in 2011. In agreement with the distribution of most flavonoid metabolites and colour-related traits in the progeny, which were in the 4 to 10-fold range, the pelargonidin-3-glucoside, which contributes to the total anthocyanin content for as much as ∼90%, displayed a 5-fold variation among the most contrasted individuals. As could be expected, the ANTHc values measured by colourimetry varied in the same range as pelargonidin-3-glucoside. In contrast, considerable variations were observed for the minor anthocyanin forms cyanidin-3-glucoside and pelargonidin-3-rutinoside, which displayed 23-fold and 114-fold variations in the progeny, respectively; and for the COLOUR value, which ranged from 0.5 to 6 in the most extreme progeny individuals. The pelargonidin-3-glucoside-malonate and cyanidin-3-glucoside additionally showed a much skewed distribution in the progeny, with a high number of individuals in which these anthocyanin compounds were not detected (Figure 1B and Supplemental Fig. 2).

For linkage map construction with SNP, SSR, SSCP and AFLP markers, only single-dose markers (SD) in backcross configuration that segregated in the expected 1:1 ratio (P < 0.01) were considered, as previously described (Lerceteau-Köhler et al. 2012; Perrotte et al. 2016). A total of 9,455 SNP markers were selected from the Axiom® IStraw90® SNP array (4,974 female and 4,481 male markers) (Bassil et al. 2015) and added to the previous linkage maps (Rousseau-Gueutin et al. 2008). In the end, the construction of the linkage maps was done with a total of 5,216 and 4,879 markers for the female and the male linkage maps, respectively. The final number of markers covered the expected 28 linkage groups for the female map and 32 linkage groups for the male map (3 additional small linkage groups were included). The length of the female and male linkage maps were 4,135.1 cM and 3,929.2 cM, respectively, with an average distance between markers of 0.8 cM.

Linkage groups were assigned to one of the seven homoeologous groups. Letters naming a given homoeologous linkage group (e.g. LG1a, b, c or d) are as previously described (Rousseau-Gueutin et al. 2008; Labadie et al. 2020). Thanks to the assignation of Affymetrix markers to the reference octoploid genome (Edger et al. 2019) (S. Verma and V. Whitaker, unpublished results), we were able to assign a given LG to one of the four subgenomes from cultivated strawberry. Detailed information on linkage map construction and assignation of specific LGs to one of the four subgenomes (referred to as *F. iinumae, F. nipponica, F. viridis* and *F. vesca* subgenomes in (Edger et al. 2019, 2020)) are provided in Supplemental Table 2.

### The Fragaria vesca subgenome is predominant in the genetic architecture of the fruit colour in cultivated strawberry

QTL were detected for all the quantitative traits analyzed using composite interval mapping (CIM) for either ‘Capitola’ or ‘CF1116’. Information on markers for male and female maps are respectively provided in Supplemental Table 3 and Supplemental Table 4. The list of significant QTLs detected for each trait including associated markers, position on male and female linkage maps, LOD score and effect is provided in Supplemental Table 5. The values of QTL thresholds at 5 and 10% that were used to select significant male and female QTLs for each trait and year are displayed in Supplemental Table 6. Distributions per homoeology groups (HGs) and by linkage groups (LGs) of significant colour-related mQTLs (PgGs, PgGsM, PgRs, CyGs, EpPgGs and Ant (summation of anthocyanins compounds)) and QTLs (ANTHc and COLOUR) are synthesized in Table 2 and Table 3 for male and female maps respectively. Graphical representation of location on male and female linkage map and Bayesian credible interval for the significant QTLs detected for the two years of observation (2010 and 2011) is shown in Supplemental Fig. 3.

Among the 65 QTLs detected for all the flavonoid and colour-related traits analyzed over the two years of study (Supplemental Table 5), a total of 37 significant colour-related QTLs (29 anthocyanins mQTLs, 4 ANTHc and 4 COLOUR QTLs) was detected (Table 1 and 2). Most LGS harbored only one region controlling variations in colour-related traits, the notable exception being the LG6a carrying 2 male and 3 female regions with colour-related QTLs (Table 1). Quantitative variations of few anthocyanin compounds were controlled by mQTLs located on different LGs (i.e. on different subgenomes) within the same HG (Table 2). The only examples are those of PgGs QTLs located on female LGs F3a and F3b in 2011 and of PgRs QTLs located on male LGs M5a and M5b and M7a and M7d in 2011. Moreover, as with the results obtained by considering all the other flavonoid mQTLs, half of the colour-related QTLs detected were located on only three LGs on male map (M1a, M3a, M6a) (11 QTLs out of 22). The overwhelming majority of the colour-related QTLs detected on female map (12 QTLs out of 15) were located on only three LGs (F2a, F3a, F6a). Remarkably, the above-mentioned LGs correspond to the *F. vesca* subgenome (Supplemental Table 2) thus highlighting the predominant role of *F. vesca* in the determination of fruit colour in cultivated strawberry. While the *F. vesca* subgenome is responsible for major variations in PgGs content (F2a QTL: R^2^= 15; M3a QTL: R^2^= 20.6; F3a QTL: R^2^= 8.8; F6a QTL: R^2^= 19.7), other subgenomes derived from *F. iinumae* (M1a QTL; R^2^= 11.65) and from *F. viridis* (M3b and M4d QTLs; R^2^>7.5) also contribute to PgGs variation’s content in the progeny (Supplemental Table 5). Likewise, while variations in PgRs are mainly controlled by the *F. vesca* subgenome, the *F. iinumae, F. viridis* and *F. nipponica-*derived subgenomes are also involved in the control of PgRs. Other subgenomes than *F. vesca* also contribute to the value of breeding traits such as COLOUR (*F. viridis* F4d; R^2^=9). Because recent studies demonstrated the crucial role played by the *MYB10-2* homoeoallele located on the LG1 *F. iinumae*–derived subgenome in natural variations in fruit colour (Castillejo et al. 2020), we investigated whether *MYB10-2* could underlie the M1a Ant, PgGs, PgRs, AfPgGs, ANTHc and COLOUR QTLs (Supplemental Table 5). We found a SNP marker (AX-89846847) very close to the *MYB10-2* homoeoallele at position 15517937 bp on the subgenome Fvb1-2 of the Camarosa reference genome (Edger et al. 2019). This marker is linked to the flavan-3-ols AfCat_2011 mQTL (Supplemental Fig. 3), but is far from the other M1a colour-related QTLs detected in our study, thereby indicating that genetic variations in *MYB10-2* are not causal.

**Table 1.** Distribution and number of significant QTLs detected for colour-related traits according to male and female linkage groups (LGs) and year.

| LGs | Nb QTLs<br>in 2010 | Nb QTLs<br>in 2011 | Total Nb<br>of QTLs | Nb regions<br>with QTLs | QTLs located on a same linkage group<sup>a</sup> |
| --- | --- | --- | --- | --- | --- |
| M1a |  | 6 | 6 | 3 | M_2011_COLOUR/2011_AfPgGs;<br>M_2011_Ant/2011_PgGs/2011_PgRs;<br>M_2011_ANTHc |
| M1c |  | 1 | 1 | 1 | M_2011_PgGsM |
| M2a/F2a | 2 | 1 | 3 | 2 | M_2011_AfPgGs; F_2010_Ant/2010_PgGs |
| **M3a/F3a** | **2** | **3** | **5** | **1** | **M_2011_COLOUR/2010_Ant/2010_PgGs/F_2011_Ant/2011_PgGs** |
| F3b |  | 1 | 1 | 1 | F_2011_PgGs |
| M4a | 1 |  | 1 | 1 | M_2010_ANTHc |
| M4d/F4d |  | 2 | 2 | 2 | M_2011_PgGs; F_2011_COLOUR |
| M5a |  | 1 | 1 | 1 | M_2011_PgRs |
| M5b |  | 1 | 1 | 1 | M_2011_PgRs |
| M6a/F6a | 2 | 8 | 10 | 5 | M_2011_PgRs; M_2011_ANTHc;<br>F_2010_Ant/2010_PgGs/2011_COLOUR/2011_CyGs;<br>F_2011_Ant/2011_ANTHc/2011_PgGs; F_2011_PgGsM |
| M6b/F6b |  | 2 | 2 | 2 | M_2011_PgRs; F_2011_AfPgGs |
| M6d |  | 1 | 1 | 1 | M_2011_PgRs |
| M7a |  | 1 | 1 | 1 | M_2011_PgRs |
| M7d |  | 1 | 1 | 1 | M_2011_PgRs |
| M41 |  | 1 | 1 | 1 | M_2011_PgRs |
| **Total** | **7** | **30** | **37** | **24** |  |

Colour-related traits considered: anthocyanins measured by LC-ESI-MS (Ant, PgGs, PgGsM, PgRs, GyGs, AfPgGs); total anthocyanins measured by colourimetry (ANTHc); and colour assessed visually (COLOUR). QTLs were identified using CIM analysis with LOD > LOD threshold at 10%. In bold, LG3a QTLs further investigated.
<sup>a</sup>QTLs overlapping within a given LG are separated by a slash (/). QTLs found on different regions within a given LG are separated by a semicolon (;). mQTLs are underlined.

**Table 2.**
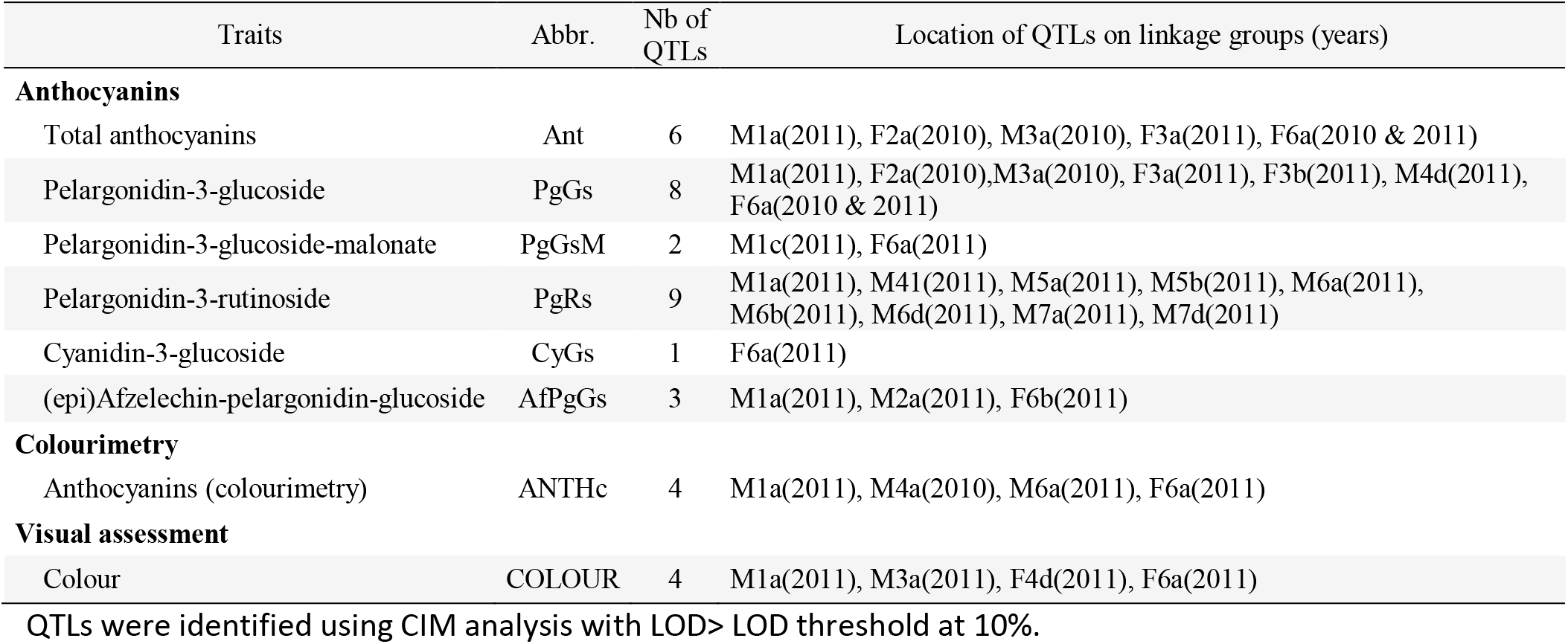
Distribution of significant anthocyanin mQTLs and visually assessed COLOUR QTL in male (M) and female (F) linkage maps.

Among the main colour-related QTLs, the mQTls located on M3a and F6a (Table 1), display opposite effects on anthocyanin content (Supplemental Table 5). To further investigate if the Ant_2010, PgGs_2010 and COLOUR_2011 QTLs had epistatic relationships, which would suggest common underlying control, we further studied the interaction of the colour- and anthocyanin-related homoeoalleles in LG3a and LG6a (Fig. 2). Variance analysis confirmed that male M3a QTLs had indeed a positive effect on trait values and female F6a QTLs a negative effect. However, because trait values were similar in individuals harbouring both M3a and F6a QTLs, we can conclude that there is likely no significant interaction between the M3a and F6a colour-related QTLs.

**Figure 2.**
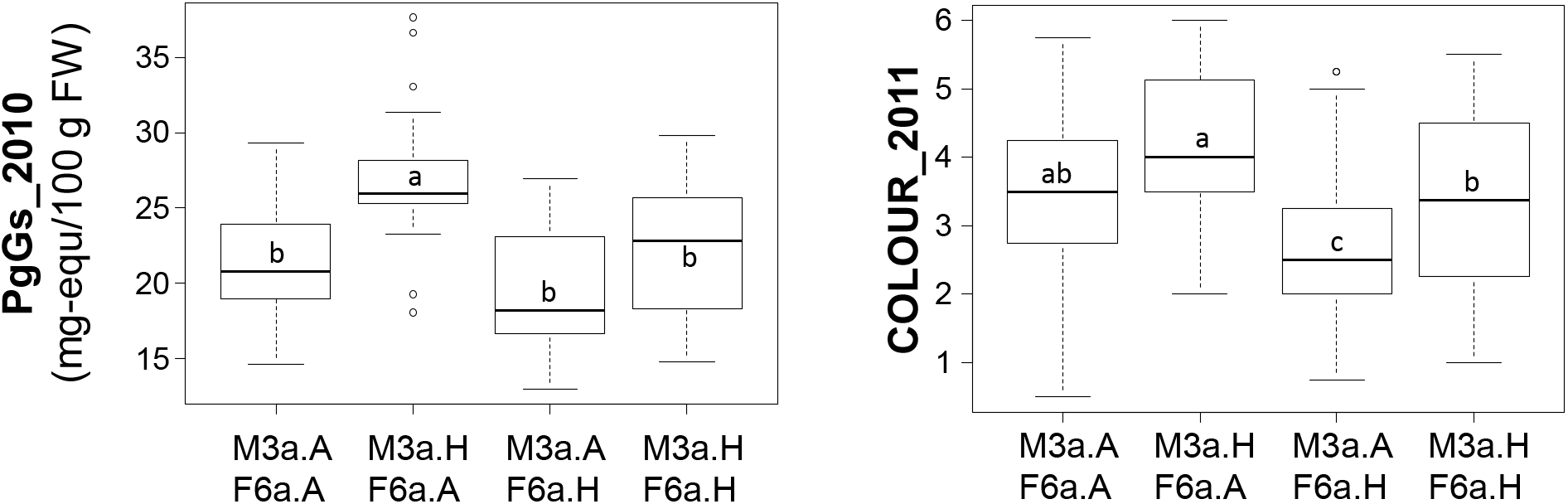
Contribution of the M3a and F6a QTLs to colour-related trait values. Effect of M3a and F6a localized alleles on PgGs content and COLOUR values. The allelic status (presence, H; absence, A) of the two markers, AX.89826440.M3a (M3a) and AX.89842368.F6a (F6a) is indicated on the abscissa. These markers were chosen because they were localized at the peak of colour-related QTLs (PgGs_2010 and Colour_2011) on M3a and F6a. Boxes represent the trait variation of individuals with the reported combination of alleles. Boxplots with the same letter are not significantly different (Kruskal-Wallis test, P < 0.05).

Although a given LG may carry different regions controlling various traits (e.g. LG6a on male or female linkage maps; Table 1 and 2), several QTLs were likely controlled by the same or by close homoeoalleles because the corresponding QTLs overlapped the Bayesian credible intervals in the parental map (Table 1; Supplemental Fig. 3). Among them, the QTLs controlling the anthocyanin PgGs and the COLOUR trait overlapped in a narrow chromosomal interval on M3a and F3a (Fig. 3A and Supplemental Table 5). This suggests the presence of potential allelic forms of the same gene or of closely linked genes responsible for the variation of quantitative characters in this region.

**Figure 3.**
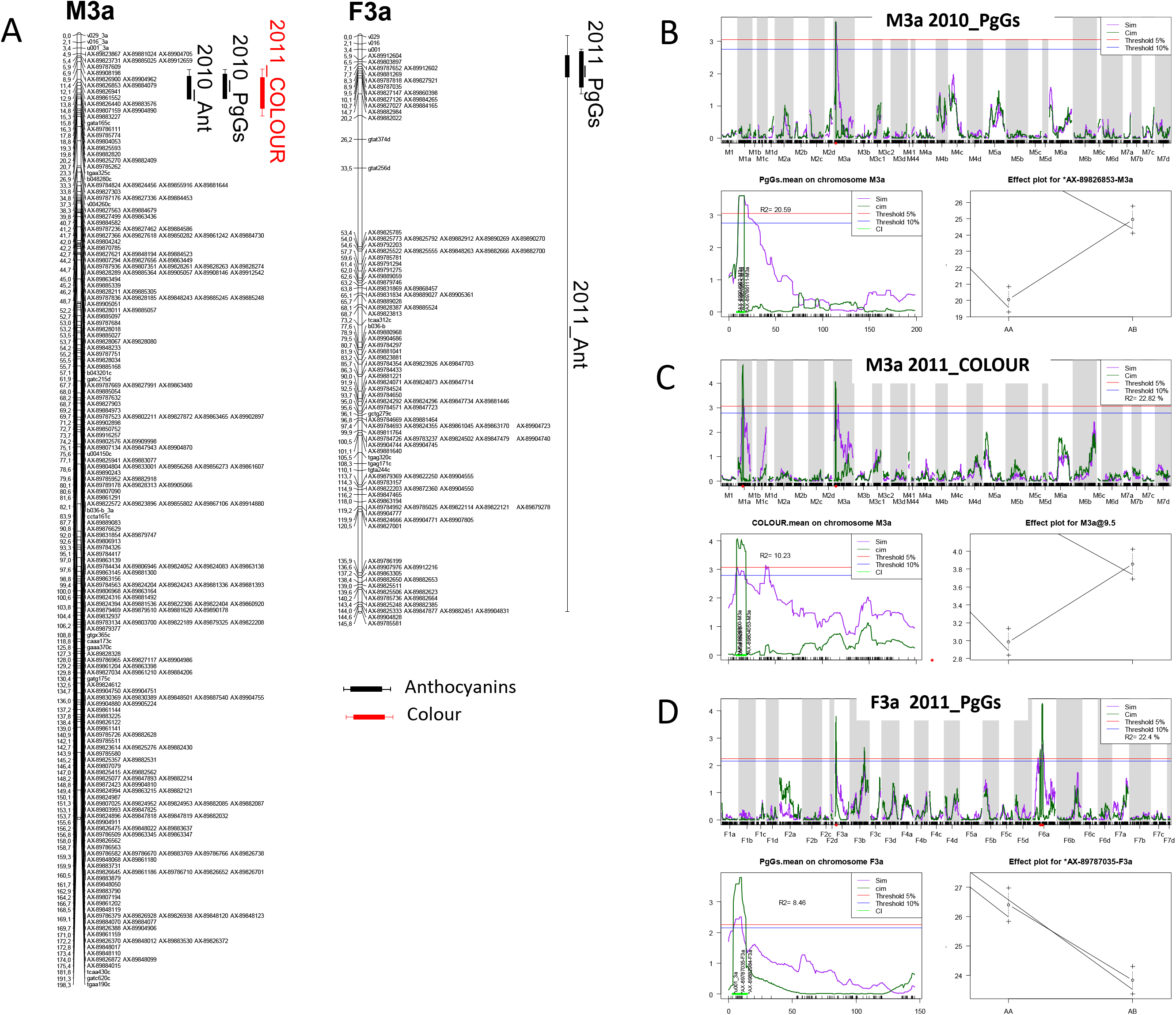
Localization of colour-related QTLs on the linkage groups M3a and F3a. **A**) Mapchart of linkage group LG3A. Linkage groups are represented in MapChart 2.3 (R. E. Voorrips, 2002) with a space of 3mm per cM. Each boxplot corresponds to a QTL identified with a threshold of 10%. Bayesian credible interval of QTL is indicated at 5%. Ant, total anthocyanins; PgGs, Pelargonidin-3-glucoside; COLOUR, visual evaluation of fruit colour. Each QTL name is preceded by the year (2010_ or 2011_). **B-D)** mQTL scans and effect of markers linked to Pelargonidin-3-glucoside (PgGs) and COLOUR (colourimetry) QTLs localized on linkage groups M3a (male) and F3a (female). For each of the three QTLs, genome scan (top), scan on specific linkage groups M3a or F3a (bottom left) and plot effect of the QTL marker (bottom right) are shown. Linkage groups M3a: (A) 2010_PgGs; (B) 2011_COLOUR and F3a: (C) 2011_PgGs. For QTL genome scan and scan of specific linkage group, LOD values are shown on the y-axis and genetic positions in centiMorgans are on the x-axis. Simple and composite Interval mapping analysis (SIM and CIM) are represented by the purple solid and the dark green solid lines respectively. Threshold of QTL detection at 5% (red) and 10% (blue) for each trait is represented. In scan on specific linkage group, Bayesian credible interval of QTL is indicated at 5% in light green. For each QTL flanking markers, markers of QTL and variance (R^2^) are indicated. For each trait, effect of QTL is indicated at the marker corresponding to the maximal LOD value of QTL.

### Anthocyanidin reductase (ANR) and MYB transcription factor underlie major QTLs controlling variations in fruit colour and pelargonidin-3-glucoside content

The ~5-fold variations in anthocyanin content in the progeny were largely due to variations in pelargonidin-3-glucoside (Labadie et al. 2020; Fig. 1B and Supplemental Fig. 2). We could detect mQTLs for total anthocyanin (Ant) and PgGs on male map (M3a) in 2010 and on female map (F3a) in 2011, and COLOUR QTL on M3a in 2011 (Fig. 3A). CIM analysis further showed that the fruit COLOUR QTL and the Ant and PgGs mQTls, which displayed high LOD scores values > 3.0, were co-located in a narrow chromosomal interval (Fig. 3B, C and D). Furthermore, the male allele from ‘CF1116’ (on M3a) had a positive effect on the levels of anthocyanins and COLOUR (Fig. 3B and C) while the female allele from ‘Capitola’ (on F3a) had the opposite effect (Fig. 3D). This points to the likely presence in this region of two different allelic variants that control these traits, one from the male parent and one from the female parent. For Ant and PgGs, the percentages of phenotypic variance explained by the M3a mQTLs were twice as high as the F3a mQTLs (Supplemental Table 5). Because allelic variants underlining these QTLs are in heterozygous state in the segregating pseudo full-sibling F1 population analyzed (Lerceteau-Köhler et al. 2012), these results indicate that a single LG3a homoeoallelic variant out of the 8 homoeoalleles present in the *F. x ananassa* genome is sufficient to affect either positively (male homoeoallelic variant) or negatively (female homoeoallelic variant) the same colour and flavonoid-related traits.

To detect the possible underlying candidate genes, we next identified for the QTLs co-localized on M3a and F3a an interval framed by SNP markers with physical positions on the diploid FvH4 *F. vesca* genome (Edger et al. 2018) that overlapped the Bayesian credible intervals. On M3a, this region is flanked by AX-89904962 and AX-89785774 Affymetrix markers and spans an interval on chromosome 3 (Fvb3) from 1.213.489 b to 2.673.762 b (Edger et al. 2018), while on F3a this region is larger (826.085 b to 2.539.615 b) and almost overlapped with the M3a interval. Based on the latest annotation of the *F. vesca* genome (Li et al. 2019), we identified a total of 305 genes in the M3a interval (Supplemental Table 7) and searched them for candidate genes possibly involved in the regulation and/or synthesis of flavonoids in strawberry fruit. Among them, we found two genes encoding MYB-related transcription factors annotated as *MYB58* (*FvH4_3g03680*) and *MYB102-like ODORANT* (*FvH4_3g03780*). A third potential candidate gene, annotated as NAD(P)-binding Rossmann-fold superfamily protein (*FvH4_3g02980*), encodes an *anthocyanidin reductase* (*ANR*) enzyme, which catalyzes the conversion of pelargonidin to epiafzelechin and of cyanidin to epicatechin (Fig. 1A).

To further investigate these genes and/or to identify additional candidate genes underlying the strawberry fruit colour and anthocyanins mQTLs co-localized on M3a and F3a, we mined transcriptome data obtained by microarray analysis of 21 individuals from the segregating population which displayed contrasted flavonoid-related phenotypes. To this end, we first constituted two groups of seven F_1_ individuals each displaying either high or low PgGs content at red ripe stage, with a PgGs content ratio between the two pools of 1.7 and 1.4 in 2010 and 2011, respectively (Fig. 4; Supplemental Table 8). We next analyzed these individuals for differentially-expressed genes (DEGs) using a custom-made oligonucleotide-based (60-mer length) platform representing a total of 18,152 strawberry unigenes (Ring et al. 2013). To identify significant DEGs present in the region of interest, we applied a Student t-test on the two phenotypic groups for all genes located in the Bayesian credible interval. Out of the 305 genes found in the M3a interval that harbors the QTL and mQTLs of interest, 232 genes including *MYB102-like ODORANT* (*FvH4_3g03780*), *MYB58* (*FvH4_3g03680*) and *ANR* (*FvH4_3g02980*) were present on the microarray (Supplemental Table 9). Among them, 49 DEGs (P value < 0.05) were found, which included *ANR* and *MYB102-like ODORANT* but not *MYB58*. The expression of the *MYB102-like ODORANT* and *ANR* genes were respectively 1.6 and 1.9 fold higher in the individuals with low PgGs content than in the individuals with high PgGs content (Supplemental Table 9 and Fig. 4). Analysis of the remaining DEGs did not highlight any additional candidate gene for the control of anthocyanin accumulation in the fruit (Supplemental Table 9).

**Fig. 4.**
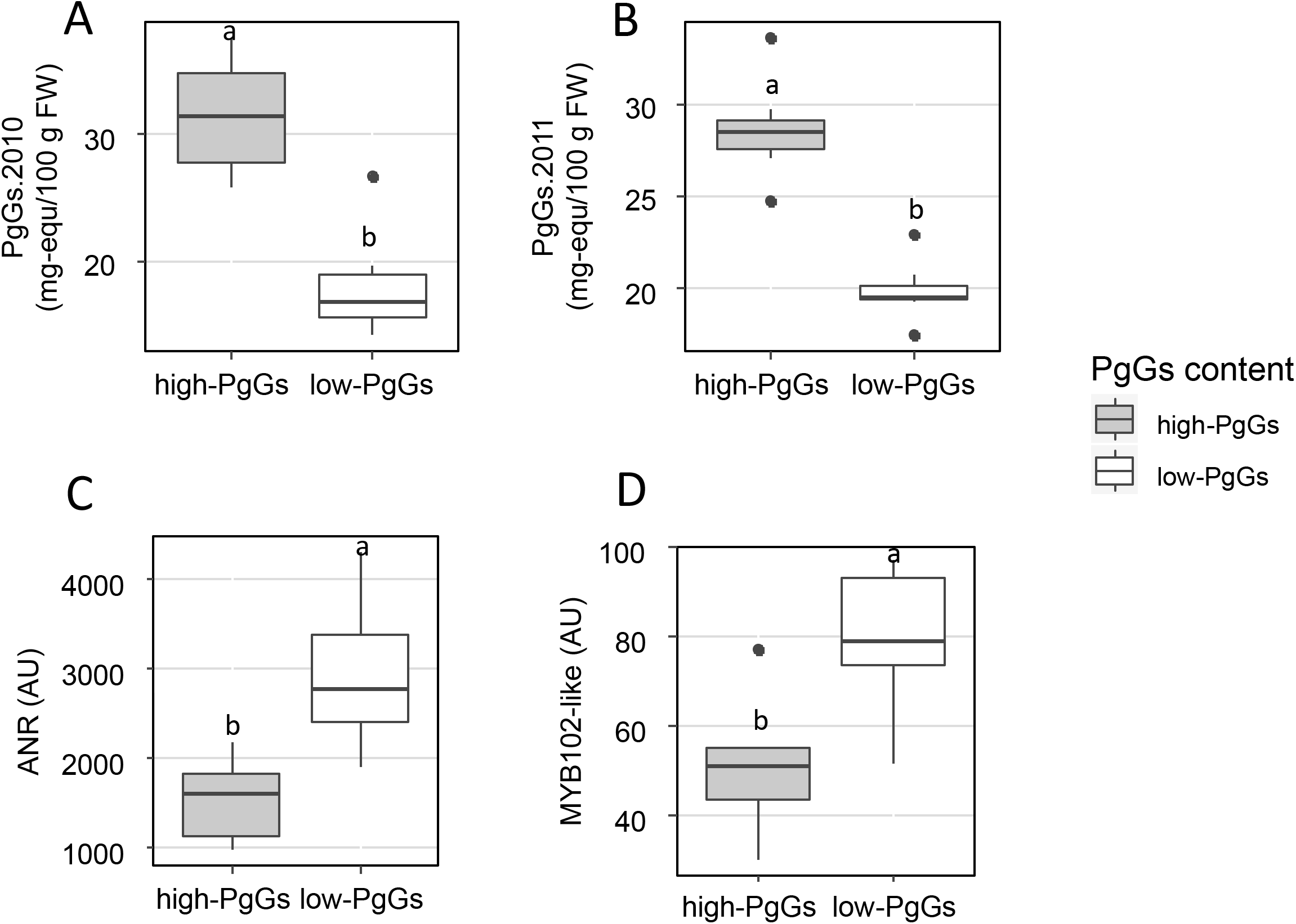
Expression of *ANR* and *MYB102-like ODORANT* in two extreme pools of individuals with contrasted pelargonidin-3-glucoside (PgGs) content. **A-B**) Two pools of seven individuals from the ‘Capitola’ and ‘CF1116’ progeny were constituted according to their PgGs content in 2010 (A) and 2011 (B). **C-D**) Microarray signal values (arbitrary units) are reported for ANR *(FvH4_3g02980)* (C) and for MYB102-like ODORANT *(FvH4_3g03780*) (D). Boxplots with different letters are significantly different (Kruskall-Wallis test, P < 0.05).

Next, we performed Illumina Hiseq 3000 whole genome sequencing of ‘Capitola’ and ‘CF1116’ parents that provided 137 and 142 million 150 pb paired-end reads, representing 50.7X and 52.7X coverage of octoploid genome (Edger et al. 2019), respectively. Alignment of paired reads to the FvH4 *F. vesca* genome (Edger et al. 2018) gave us access to the sequence polymorphisms of the MYB102-like ODORANT and ANR genes in both parents. Because these two genes are not only positional but also expressional candidate genes, we first analyzed sequence polymorphisms in regions that may affect gene regulation of *MYB102-like ODORANT* and *ANR*. In both *MYB102-like ODORANT* and *ANR* genes, several SNPs and insertions were found in the 1 kb region upstream of the 5’untranslated region (UTR) by comparison of the ‘Capitola’ and ‘CF1116’ sequences. Only polymorphisms with segregation ratio fitting the 1:7 ratio expected for single male or female homoeoalleles (Chi2 test; P > 0.05) (Fig. 5) are discussed hereafter. Interestingly, we found a 18 bp deletion in the 5’UTR of *ANR* located from – 109 to – 127 upstream of the start codon in the ‘CF1116’ (male parent) (Fig. 5). Analysis of the corresponding CTTCTTCCTCTTCTTCTT sequence with the plant regulation analysis platform PlantRegMap (Jin et al. 2017) predicted the deletion to be in the 21 nucleotide binding site of the *Arabidopsis* MYKC-MADS Agamous-like transcription factor AT2G45660. Blast analysis of *F. vesca* v1.0 ab hybrid reference genomic sequence at GDR (Jung et al. 2019) using AT2G45660 protein sequence as a query and visualization of fruit gene expression of top ten hits at *F. vesca* eFP browser (Hawkins et al. 2017) allowed the identification of four fruit MYKS-MADS box genes expressed in strawberry fruit cortex (*gene24852*, *gene04229*, *gene26119* and *gene06301*) that may possibly interact with the sequence deleted in ‘CF1116’ ANR 5’UTR. Other small insertion/deletions (INDELS) or SNPs found in 5’UTR or promoter regions from *MYB102-like ODORANT* and *ANR* genes did not match known motifs. In a second step, we analyzed the protein coding regions of MYB102-like ODORANT and ANR (Fig. 5). While SNP polymorphisms found in ANR are synonymous and do not affect the function of the protein, two non-synonymous missense variations leading to H146Q and I206N amino acid substitutions were found at the expected 1:7 segregating ratio in exon 3 of the MYB102-like ODORANT from ‘Capitola’ (female parent).

**Fig. 5.**
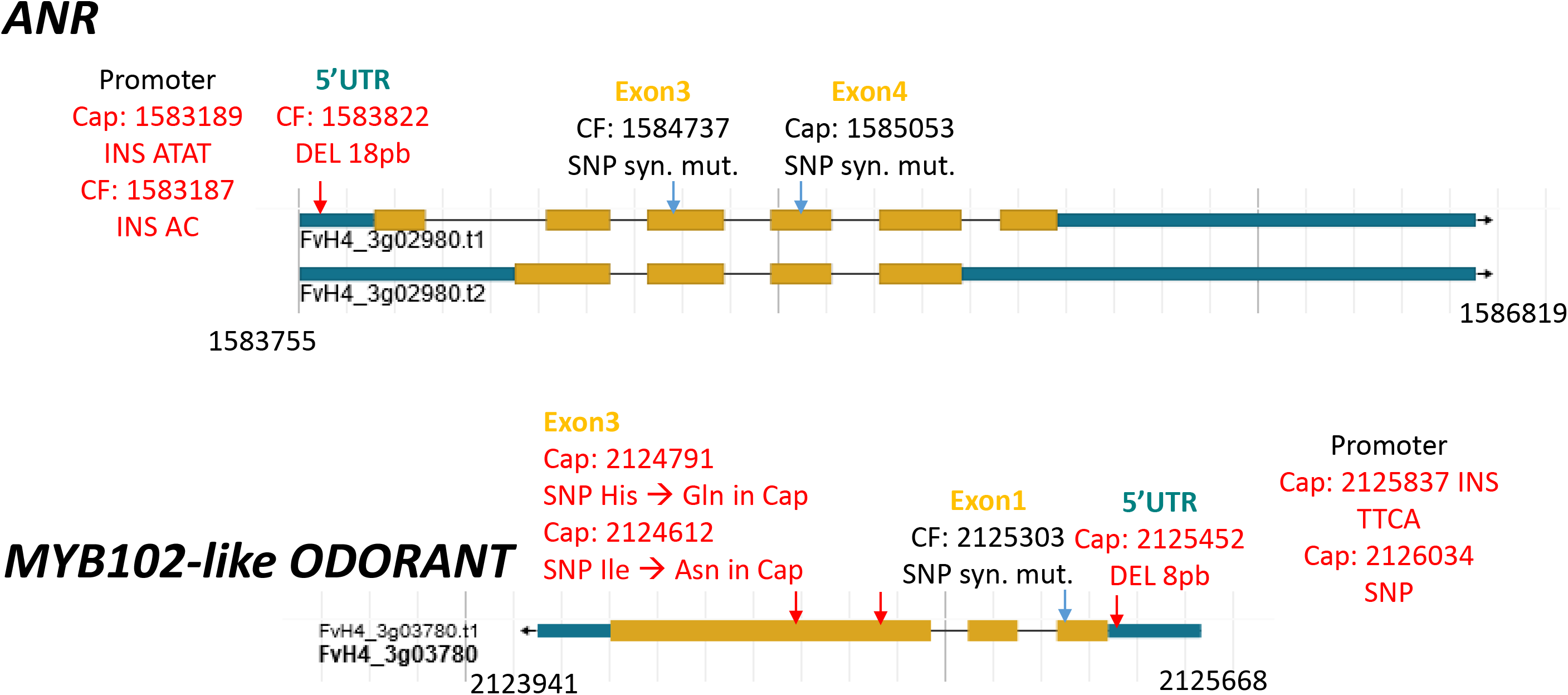
Polymorphisms identified in the *MYB102-like ODORANT* and *ANR* genes underlying the LG3a colour-related QTLs by whole genome sequencing of the progeny parents. Schematic representation of the *MYB102-like ODORANT (FvH4_3g03780)* and *ANR (FvH4_3g02980)* genes was extracted from GDR (https://www.rosaceae.org/). Polymorphism was searched in *cis*-regulatory regions including promoter (~1000pb upstream of the 5’UTR), 5’UTR region and coding region (exons). Only polymorphisms between ‘Capitola’ (Cap) and ‘CF1116’ (CF) that respond to a Chi2 test for the presence of 1 allele out of the 8 alleles (1:7 ratio) are reported. In red, polymorphisms with potential effect on expression or function. Type of polymorphisms are described: DEL, deletion; INS, insertion; SNP, single nucleotide polymorphism. syn. mut. for synonymous mutation. Three letter code is used for amino acid description. Numbers refer to positions on the Fvb3 release of the reference genome (Fragaria vesca Whole Genome v4.0.a1 Assembly; Edger et al. 2018).

To further explore whether SNP markers linked to either the LG3a *MYB102-like ODORANT* or the LG3a *ANR* gene would be useful for strawberry breeding, we analyzed the association between the SNP markers linked to the F3a and M3a PgGs mQTLs (Fig. 3) and fruit PgGs content. To this end, we used the 14 selected individuals displaying high or low PgGs content and ANR and MYB expression in the ripe fruit (Fig. 4). While no significant results were found for the F3a PgGs mQTL (R^2^= 8.8), all the individuals with high-PgGs content showed the presence of the SNP marker AX-89826853 (position: 1981447 bp on FvH4), which is strongly linked to the major M3a PgGs mQTL (R^2^=20.6) and is close to the ANR gene (position: 1583755 bp on FvH4). Conversely, the low-PgGs individuals did not display the presence of this marker. This result indicates that a MAS strategy based on the AX-89826853 marker can be envisaged for breeding strawberry varieties with improved fruit colour.

## Discussion

Flavonoid biosynthesis and, in particular, anthocyanin biosynthesis, have been thoroughly investigated in strawberry because of their considerable importance for the sensorial and nutritional quality of the fruit, which is a major breeding target (Mezzetti et al. 2018). So far, most studies were focused on fruit composition analysis in various strawberry species and varieties (Muñoz et al. 2011; Roy et al. 2018) and on candidate genes encoding known regulators or structural enzymes of the flavonoid pathway. As evidenced by reverse genetic studies on wild diploid and cultivated strawberry, the phenylpropanoid-regulating MYB, bHLH and WD-repeat proteins transcription factors play prominent roles in anthocyanin regulation (Jaakola 2013; Härtl et al. 2017; Zhang et al. 2020). Additional candidates are the structural enzymes involved in flavonoid and anthocyanin pathways (Almeida et al. 2007; Griesser et al. 2008; Fischer et al. 2014) and in connected pathways leading to phenylpropanoid-derived compounds (Ring et al. 2013). To date, insights into the genetic architecture of anthocyanin biosynthesis in strawberry were limited to few studies on the wild diploid *F. vesca* (Urrutia et al. 2016) and on the cultivated strawberry (Lerceteau-Köhler et al. 2012; Labadie et al. 2020). Identification of the genetic variations underlying fruit colour were even scarcer until the recent studies of the contribution of MYB10 to the white fruit phenotype in the wild diploid strawberry *F. nilgerrensis* (Zhang et al. 2020) and to the variation of fruit skin and flesh colour in woodland diploid strawberry *F. vesca* and in cultivated octoploid strawberry *F. × ananassa* (Castillejo et al. 2020).

In the last decade, novel tools including high density SNP arrays (Bassil et al. 2015) for high resolution genotyping of cultivated strawberry *F. x ananassa* and MS-based methods for the exhaustive analysis of specialized metabolites in strawberry fruit (Ring et al. 2013; Haugeneder et al. 2018) have been established. In parallel, considerable advances have been made in our understanding of strawberry genomes, including the release of high quality genome sequences of diploid woodland strawberry *F. vesca* (Shulaev et al. 2011; Edger et al. 2018) and octoploid *F. x ananassa* (Edger et al. 2019; Liston et al. 2020). Thanks to these advances, we could thoroughly investigate the genetic architecture of the colour-related QTLs previously identified (Lerceteau-Khöler et al. 2012) by (*i*) high resolution mapping to single linkage groups (LGs) of colour traits broken down into their individual components, (*ii*) the identification of the donor subgenomes carrying the homoeoalleles responsible for colour variation (Edger et al. 2019), and (*iii*) the discovery of causal homoeoallelic variations underlying several colour QTLs. This opens the possibility to use MAS to breed strawberry varieties with the desired colour characteristics.

### Flavonoids and anthocyanins metabolism are largely controlled by the dominant woodland strawberry *F. vesca* subgenome in cultivated strawberry fruit

Thanks to the precise assignation of a QTL to a narrow genomic region, the genetic architecture of flavonoid-dependent traits can now be explored to an unprecedented level in cultivated strawberry. In polyploid plant species, each trait is likely controlled by homoeologous gene series or homoeoalleles. Homoeoalleles are alleles located at orthologous positions on the subgenomes that compose the polyploid species (Gaston et al. 2020). The recent sequencing of the octoploid cultivated strawberry genome (Edger et al. 2019; Edger et al. 2020) showed that it is composed of four diploid subgenomes including the wild diploid woodland strawberry *F. vesca*, a diploid ancestor of *F. iinumae*, and two species relative to *F. viridis* and *F. nipponica*, the origin of which is under study (Liston et al. 2019). QTL localization on linkage groups together with the positions of the SNP markers (Bassil et al. 2015) on the various subgenomes from *F. x ananassa* (Supplemental Table 2) allowed us to identify the subgenome(s) that are likely responsible for flavonoid trait variations. Remarkably, our results showed that most variations in anthocyanins, and more broadly in flavonoids, were controlled by mQTLs located on a single linkage group (i.e. on a single subgenome) within a given homoeology group. As an example, the *F. vesca* subgenome was likely responsible for the major pelargonidin-3-glucoside mQTLs located on M3a, F3a and F6a linkage groups. More generally, the main linkage groups accounting for half of male colour-related QTLs (3 LGs) and for the majority of female QTLs (3 LGs) can be attributed to the *F. vesca* subgenome, thus highlighting the predominant role of *F. vesca* in fruit flavonoid metabolism and therefore in *F. x ananassa* fruit quality. These findings support the conclusions from Edger et al. (2019) who showed that *F. vesca* is the dominant subgenome in the octoploid strawberry and suggested that *F. vesca* homoeologs are responsible for almost 89% of the biosynthesis of anthocyanins in octoploid strawberry.

Though the precise assignation to a given subgenome is still delicate and debated (Liston et al. 2020), our results also point out that, in addition to *F. vesca*, other subgenomes may play key roles in flavonoid metabolism, for example the *F. iinumae* and *F. viridis*-derived subgenomes for pelargonidin-3-glucoside content; the *F. iinumae, F. viridis* and *F. nipponica* subgenomes for pelargonidin-3-rutinoside content; and the *F. viridis* for COLOUR. A striking example is the crucial role played by *MYB10*-2 homoeoallele (*F. iinumae*–derived subgenome) in the regulation of fruit colour in cultivated strawberry (Castillejo et al. 2020). Careful comparison of the location of the *MYB10*-2 homoeoallele and of the colour-related QTLs detected in our study showed that *MYB10*-2 has likely no incidence on fruit colour variations in the progeny studied here. These findings further indicates that additional, *MYB10*-2-independent, genetic variations carried by the *F. iinumae*–derived subgenome are responsible for the colour-related QTLs found on M1a including the pelargonidin-3-glucoside, pelargonidin-3-rutinoside, (epi)afzelechin-pelargonidin-glucoside and COLOUR QTLs. From a broader perspective, it cannot be excluded that subgenome-dependent control of anthocyanin content and composition may become prevalent in different genetic contexts or in adverse environmental conditions, as hypothesized in Lerceteau-Köhler et al. (2012). By deciphering the contribution of the four different subgenomes of *F. x ananassa* to the genetic architecture of the flavonoid and anthocyanin composition, our results further open the possibility to identify the subgenome-specific homoeoalleles responsible for major variations in colour-related traits in cultivated strawberry.

### *ANR* and *MYB* genes from the *F. vesca* subgenome are likely candidates for controlling variations in strawberry fruit colour

The major QTLs detected here on LG3a for the total anthocyanins, for pelargonidin-3-glucoside and for the visually scored fruit colour are all co-located at the beginning of the linkage group (Fig. 3A) and overlap the colour-related QTLs previously detected over three years of study for the physical parameters L and b (colour space values) (Lerceteau-Köhler et al. 2012), indicating that these colour QTLs are robust. We could further decipher the genetic architecture of the colour trait and show that: (*i*) pelargonidin-3-glucoside mQTLs displaying high LOD scores values > 3.0 are co-located on both male (M3a) and female (F3a) maps in a narrow chromosomal interval encompassing 305 genes, (*ii*) the male homoeoallele from ‘CF1116’ has a positive effect on pelargonidin-3-glucoside content (Fig. 3B and C) while the female homoeoallele from ‘Capitola’ has the opposite effect (Fig. 3D), (*iii*) homoeoallelic variants of two candidate genes encoding anthocyanidin reductase (ANR) and MYB102-like ODORANT, both of which are differentially expressed in pools of individuals with contrasted pelargonidin-3-glucoside contents, are found in the male (‘CF1116’) and female (‘Capitola’) parents, respectively.

MYB transcription factors (TFs) belong to a family of TFs playing predominant roles in the control of phenylpropanoid pathway (Ma and Constabel 2019). To date, various MYB transcription factors and the MYB/bHLH/WD-repeat protein complex are known to regulate anthocyanin biosynthesis in strawberry (Salvatierra et al. 2013; Schaart et al., 2013; Medina-Puche et al. 2014; Wang et al. 2020). The best studied strawberry MYB is MYB10, whose mutation induces a loss of red fruit colour in the cultivated strawberry and in the wild diploid species *F. vesca* and *F. nilgirensis* (Medina-Puche et al. 2014; Hawkins et al. 2016; Wang et al. 2020; Castillejo et al. 2020; Zhang et al. 2020). However, the most likely MYB candidate gene underlying the female F3a pelargonidin-3-glucoside mQTL is not homologous to a well characterized strawberry MYB. It encodes a MYB102-like ODORANT protein homologous to the petunia R2R3-MYB ODORANT1 (ODO1) described as regulator of the floral-scent related genes of the phenylpropanoid biosynthesis pathway (Spitzer-Rimon et al. 2010). Though ODO1 has no effect on anthocyanin biosynthesis in petunia (Spitzer-Rimon et al. 2010), its ectopic expression in tomato activates phenylpropanoid metabolism without affecting volatiles (Dal Cin et al. 2011). In addition to the two SNPs leading to synonymous amino acid changes in *ODO1* coding region, the *MYB102-like ODORANT* gene from ‘Capitola’ LG3a carries several SNPs and INDELS in the 5’UTR and promoter regions, which opens the possibility that transcriptional regulation of this MYB gene (Fig. 4) may be responsible for variation in anthocyanin biosynthesis, regardless of its weak expression in the fruit (Hawkins et al. 2017). Altogether, these information warrant further investigation of *MYB102-like ODORANT* function in strawberry fruit.

In strawberry, ANR converts respectively the anthocyanidins pelargonidin and cyanidin to the flavan-3-ols, either to epiafzelechin, which can be derived to epiafzelechin-glucose, or to epicatechin (Fig. 1A). ANR is encoded by a single gene with two predicted alternative transcripts in *F. vesca* genome. Mining the *F. vesca* gene expression atlas database (Hawkins et al. 2017) further showed that *ANR* is very highly expressed in fruit cortex at early developing stages until white fruit stage (onset of ripening) and moderately thereafter. ANR is therefore a strong expressional candidate for the regulation of anthocyanins in strawberry fruit. It is to be noted that a single homoeoallele from the ‘CF1116’ *F. vesca*–derived subgenome out of the 8 homoeoalleles present in the octoploid genome would be responsible for the large effect of the M3a QTL. Because this homoeoallelic variant is likely overexpressed in individuals with low pelargonidin-3-glucoside content (Fig. 4) and that the deletion identified in ANR 5’UTR falls within a MYKS-MADS box binding domain, the candidate MADS box would be involved in the negative regulation of the ANR gene. During fruit ripening, the lack of negative regulation of ANR (no MADS box binding) would maintain a high ANR transcript level in the ripening fruit (Fig. 4), thereby keeping the phenylpropanoid flux channeled towards flavan-3-ols synthesis at the expense of anthocyanin formation (Fig. 1A). The down-regulation of ANR at earlier stages of fruit development should therefore trigger the enhanced accumulation of anthocyanins in the fruit before ripening. Conversely, its up-regulation in the ripening fruit would lead to reduced anthocyanin accumulation. Indeed, we previously demonstrated (Fischer et al. 2014) that ANR silencing in strawberry leads to the early accumulation of anthocyanins in the fruit, by re-routing the flux from flavan-3-ols to anthocyanins. At the opposite, Zhao et al. (2017) recently showed that overexpression of tea (*Camellia sinensis*) ANR genes in tobacco effectively leads to a significant loss of flower red-pigmentation by reducing their anthocyanin content. These results further support the hypothesis that the increased expression of a LG3a-located ‘CF1116’ ANR homoeoallele from the *F. vesca*–derived subgenome is responsible for the increased accumulation of pelargonidin-3-glucoside that likely underlies the M3a colour mQTL.

The involvement of MADS-box genes in anthocyanin accumulation have been reported in different species including Arabidopsis (Nesi et al. 2002) and other Rosaceae fruit species such as bilberry (Jaakola et al. 2010) and pear (Wang et al. 2017). Among the four MADS box candidates expressed in fruit cortex, two of them (*gene26119*, *gene06301*) that are expressed at late stages of fruit development (Hawkins et al. 2017) show a pattern of expression opposite to that of ANR, which makes them potential candidates for regulating ANR in strawberry fruit. Because the control of anthocyanin accumulation by MADS-box TFs remains unclear, the regulation of ANR by MADS-box genes in strawberry fruit is a working hypothesis worth exploring.

## Conclusions

We demonstrated for the first time in this study that, by breaking down fruit colour traits into smaller components corresponding to individual flavonoid compounds and by delineating the resulting mQTLs to narrow genomic regions corresponding to specific subgenomes of the cultivated strawberry, the identification of homoeoallelic variants likely underlying the flavonoid mQTLs is now feasible. By deciphering the genetic architecture of fruit colour, our results further support, at the genetic level, the hypothesis that the flavonoid metabolism is mainly controlled by the *F. vesca*-derived subgenome in the octoploid cultivated strawberry. Our findings also point out that other subgenomes from the cultivated strawberry effectively contribute to the control of fruit colour. We additionally provide complementary genetic and genomic evidences that natural diversity found in ANR plays a main role in the modulation of fruit anthocyanin content in cultivated strawberry, the key function of ANR being supported by previous reverse genetic studies. Moreover, our study opens new research avenues on the hitherto unknown role in the regulation of anthocyanin accumulation in strawberry fruit of several TFs belonging to the MADS-box and MYB families. From a more applied perspective, the possibility to identify genetic variations underlying major anthocyanins mQTLs in strawberry provides the means to develop MAS markers closely linked to fruit colour, which is a main breeding target from the consumer’s perspective. An important step toward reaching that end is the identification of a SNP marker close to the ANR gene underlying the major pelargonidin-3-glucoside mQTL located on M3a, which explains > 20% of the phenotypic variance.

## Experimental procedures

### Plant materials and preparation

A pseudo full-sibling F_1_ population of 165 individuals obtained from a cross between the variety ‘Capitola’ (CA75.121-101 x Parker, University of California, Davis, USA) and the advanced line ‘CF1116’ ([‘Pajaro’ x (‘Earlyglow’ x ‘Chandler’], reference from the Ciref, France) was developed. The ‘Capitola’ and ‘CF1116’ parents display contrasting fruit colour together with differences in fruit shape and weight, firmness, sweetness and acidity (Lerceteau-Köhler et al. 2012). For each of the two consecutive study years (2010 and 2011), cold-stored strawberry plants planted in 2009 and 2010 were grown in soil-free pine bark substrate under plastic tunnel with daily ferti-irrigation and control of biotic stresses. Mapping population included a total of 165 individuals over the two study years. Within this progeny, 72 and 131 individuals, including parents, were respectively phenotyped in 2010 and in 2011. Fruits were harvested at red ripe stage, when red colouration of the fruit is homogeneous, and processed as previously indicated (Labadie et al. 2020) to produce frozen powder samples that were further stored at −80°C until use for chemical analyses.

### Fruit colour evaluation and flavonoid chemical analyses

In 2011, for both harvests, red ripe fruits (4-5 fruits per harvest) were also photographed side-by-side with the strawberry colour chart from Ctifl (http://www.ctifl.fr/Pages/Kiosque/DetailsOuvrage.aspx?IdType=3&idouvrage=833). Fruit colour was then scored by two independent persons (2 replicates) on a scale from 0 (very pale red-orange) to 6 (very dark red) using the photographs. A mean score value was then obtained for each genotype (Supplemental Fig. 1).

Analysis of polyphenolic metabolites by LC–ESI-MS was done as previously described (Ring et al. 2013). A total of 16 traits encompassing individual phenolic metabolites [13 metabolites: anthocyanins (pelargonidin-3-glucoside; pelargonidin-3-glucoside-malonate; pelargonidin-3-rutinoside; (epi)afzelechin-pelargonidin-glucoside; cyanidin-3-glucoside), flavonols (kaempferol-glucuronide; kaempferol-glucoside; kaempferol-coumaroyl-glucoside; quercetin-glucuronide), flavan-3-ols (catechin; (Epi)catechin dimers; (epi)afzelechin-(epi)catechin dimers; (epi)afzelechin-glucoside) and their sum [3 traits: total anthocyanins, total flavonols, total flavan-3-ols] were measured for the two years. Analyses of pooled frozen powder samples were carried out in 2010 and 2011 on 6 replicates for parents and on 3 replicates for individuals from the progeny.

Extraction and measurement of total anthocyanin content (ANTHc) by colourimetric assay were as described in Labadie et al. (2020). Results are expressed as mg pelargonidin-3-glucoside equivalents/100 g fresh weight. For each genotype, 4 technical repeats from the pooled two-harvest-fruit-powder were performed.

### Genotyping

DNA from the parental lines ‘Capitola’ and ‘CF1116’ and from 165 individuals from the mapping population was extracted using DNeasy Plant Mini kit (Qiagen, Hilden, Germany) and hybridized with the Affymetrix® 90 K Axiom® SNP array (Affymetrix, CA, USA) at CeGen USC (Santiago de Compostela, Spain). IStraw90® SNP array includes 138,000 SNP markers corresponding to 90,000 localizations on wild diploid strawberry *F. vesca* ‘Hawai’ v1.1 genome (Bassil et al. 2015). Analysis was performed using Genotyping console™ and SNPpolisher© (Affymetrix, CA, USA) following manufacturers recommendations.

### Linkage maps and QTL analysis

Frequency distribution of each trait was represented using ggplot2 (v3.2.1; Wickham et al., 2019) r-package. Single dose markers (SD) from the Affymetrix array (Bassil et al. 2015) that were in backcross configuration and segregated 1:1 (Rousseau-Gueutin et al. 2008) were used in combination with previously mapped SSR, SSCP and AFLP markers (Gaston et al. 2013; Labadie et al. 2020) for map construction using JoinMap 5.1 software (Van Ooijen 2011). Grouping was performed using independence log of the odds (LOD) and the default settings in JoinMap® and linkage groups were chosen from a LOD higher than 10 for all of them. Map construction was performed using the maximum likelihood (ML) mapping algorithm and the following parameters: chain length 5,000, initial acceptance probability 0,250, cooling control parameter 0,001, stop after 30,000 chains without improvement, length of burn-in chain 10,000, number of Monte Carlo EM cycles 4, chain length per Monte Carlo EM cycle 2,000 and sampling period for recombination frequency matrix samples: 5.

For QTL analysis, the female and male linkage parental maps based on the 165 individuals were used separately. Phenotypic data of the 72 and 131 individuals for 2010 and 2011, respectively, were represented by the mean value of the three replicates. QTL detection was performed by simple interval mapping (SIM) using R/QTL (Broman et al. 2003). Permutation analysis, using 1000 permutations, was performed to calculate the critical LOD score. QTL with LOD values higher than the LOD threshold at *P* ≤ 0.05 were considered significant. When one QTL was found significant, we used composite interval mapping (CIM) (Zeng 1993, 1994) with one co-variable at the position of the significant QTL and reiterated the analysis until no new significant QTLs were detected. Bayesian credible interval was calculated using the function ‘bayesint’ at probability of 0.95. The proportion of phenotypic variance explained by a single QTL was calculated as the square of the partial correlation coefficient (R^2^). Mapping results are displayed using MapChart (Voorrips, 2002).

### Gene expression analysis

A custom-made oligonucleotide-based (60-mer length) platform designed (Roche NimbleGen) from non-redundant *Fragaria vesca* strawberry sequences (Fv_v1.0 Shulaev et al. 2011) representing a total of 18,152 unigenes (Ring et al. 2013) was used for the analysis of differentially-expressed genes between 21 F_1_ genotypes from the ‘Capitola’ x ‘CF1116’ population displaying contrasted phenotypic values for flavonoids and colour-related traits. RNA extraction from red ripe fruits and microarray processing and data analysis were as previously described (Ring et al. 2013). Student’s t test was used with a confidence of *P* < 0.05 to detect statistically significant differences.

### Whole genome sequencing

Whole genome sequencing of the two parents ‘Capitola’ and ‘CF1116’ was performed using paired-end Illumina sequencing with a ~ 50X coverage of the *F. x ananassa* genome. Illumina paired-end shotgun-indexed libraries were prepared and sequenced using an Illumina HiSeq 3000 at the Institut National de la Recherche Agronomique GeT-PlaGe facility (Toulouse, France), operating in a 150-bp paired-end run mode. Raw fastq files were mapped to the FvH4 reference genome sequence *F. vesca* (Edger et al. 2018) using bwa mem algorithm for the alignment of paired-end (150 bp) Illumina reads. Polymorphisms between ‘Capitola’ and ‘CF1116’ were identified using Integrative Genomics Viewer (IGV) (Robinson et al. 2011). All identified polymorphisms were tested for the 1:7 (mutant allele:WT alleles) segregation ratio for goodness-of-fit to theoretical ratio (Chi2 test) when considering the hypothesis that, out of the eight homoeoalleles expected in the octoploid *F. x ananassa*, one single homoeoallele (mutant allele) specific to ‘Capitola’ or to ‘CF1116’ controls the trait.

## Acknowledgements

We thank Sujeet Verma and Vance Whitaker (University of Florida) for providing the assignation of Affymetrix markers to the reference octoploid genome. We thank Karine Tallès and Gabriel Jousseaume for fruit harvests and colourimetric measurements. The authors gratefully acknowledge support from Région Nouvelle Aquitaine (AgirClim project N°2018-1R20202), the EU ERANET (FraGenomics N°PCS-08-TRIL-00) and the European Union’s Horizon 2020 research and innovation program (GoodBerry project N° 679303). NGS data were produced by GeT-PlaGe Toulouse, France.

## Accession numbers

*Anthocyanidin reductase: FvH4_3g02980 (gene24665); MYB102-like ODORANT: FvH4_3g03780 (gene30725); MYB58: FvH4_3g03680 (gene30736).*

## Online resource

**Supplemental Figure 1.** Colour score of ripe fruits.

**Supplemental Figure 2.** Distribution in 2011 of the progeny mean for flavonoid metabolites, total anthocyanins assessed by colourimetry and colour assessed visually.

**Supplemental Figure 3.** Localization of all detected QTLs on male (A) and female (B) linkage maps for flavonoid metabolites, total anthocyanins assessed by colourimetry and colour assessed visually.

**Supplemental Table 1.** COLOUR trait values for ‘Capitola’ and ‘Capitola’ x ‘CF1116’ progeny in 2011.

**Supplemental Table 2.** Correspondence between names of linkage groups from female (‘Capitola’) and male (‘CF1116’) parents and the *F. vesca*, *F. viridis*, *F. iinumae* and *F. nipponica*–derived subgenomes.

**Supplemental Table 3.** Pointwise of male linkage map.

**Supplemental Table 4.** Pointwise of female linkage map.

**Supplemental Table 5.** Significant QTLs detected in male and female linkage maps for all traits and for two years based on CIM analysis with LOD> LOD threshold 5 and 10% for each trait.

**Supplemental Table 6.** Values of the thresholds at 5 and 10% used for CIM QTL analysis. The thresholds were calculated on 1000 permutations for each trait for male and female in 2010 and 2011.

**Supplemental Table 7.** List of genes from the FvH4_v4.0.a2 version of the *Fragaria vesca* reference genome (Li et al. 2019) located in the male M3a colour-related QTLs.

**Supplemental Table 8.** Phenotypic data of the two sets of individual with extremes phenotypes for pelargonidin-3-glucoside that were studied using microarray.

**Supplemental Table 9.** List of genes included in the Bayesian credible interval of M3a colour-related QTLs that were tested by microarray.

