## Supplemental Fig 1 to 3 for "The genetic architecture of fruit colour in strawberry (*Fragaria × ananassa*) uncovers the predominant contribution of the *F. vesca* subgenome to anthocyanins and reveals underlying genetic variations"

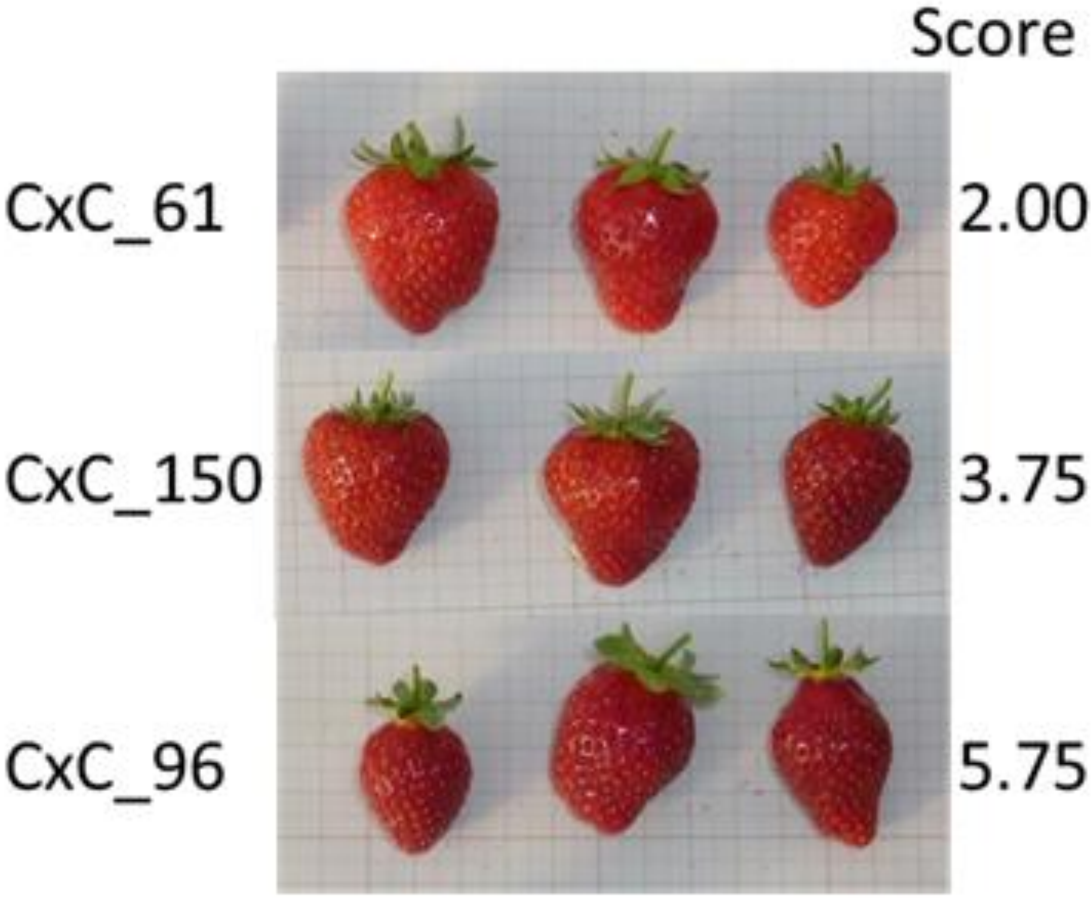


**Supplemental Fig 1. Colour score of ripe fruits.**
Visual evaluation of colour on a scale from 0 (white) to 6 (dark red) on ripe fruits harvested in 2011. Three individuals from the progeny issued from the cross between ‘Capitola’ and ‘CF1116’ (CxC) are shown. Colour score is indicated on the right.

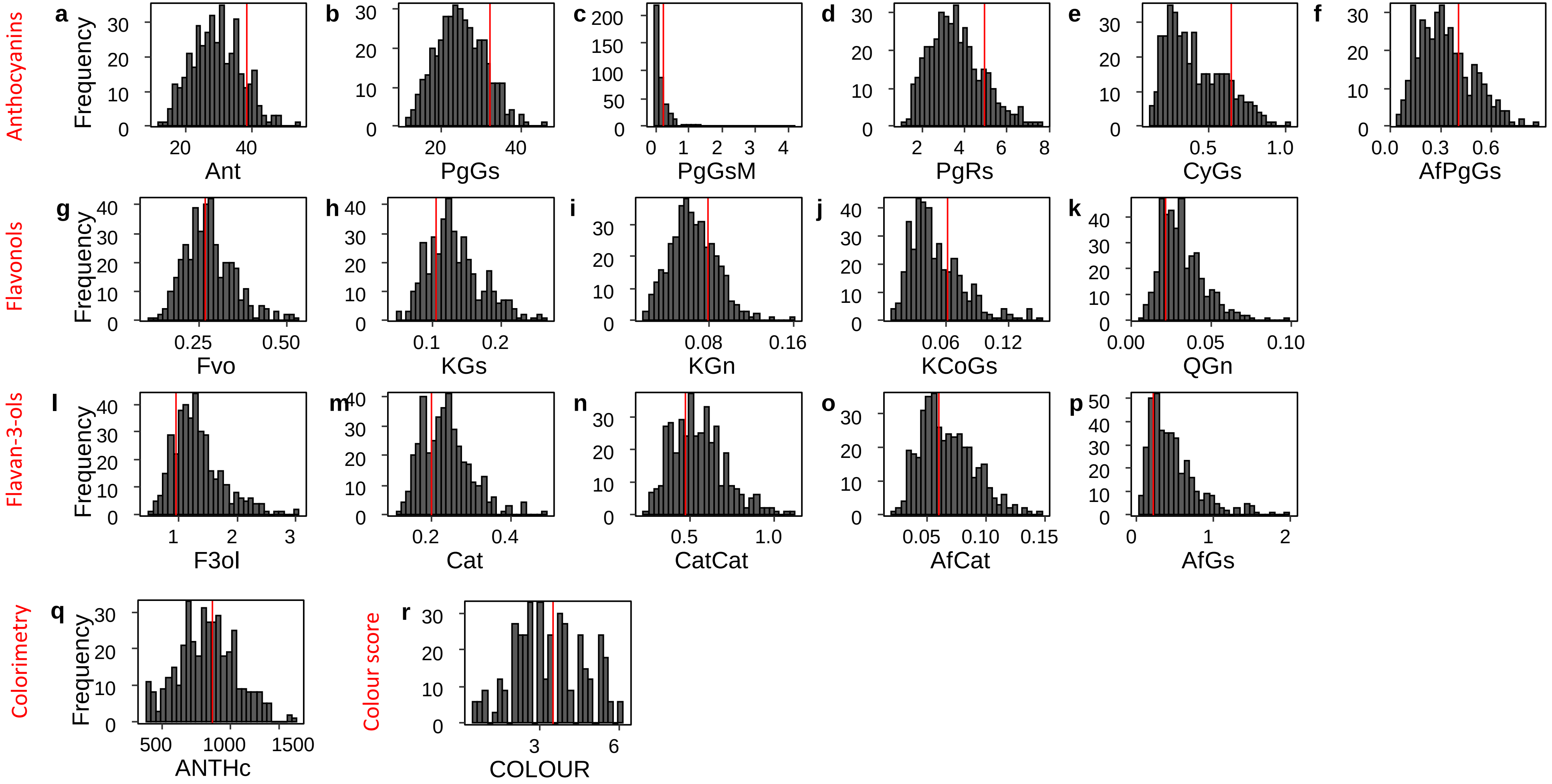


**Supplemental Figure 2. Distribution in 2011 of the progeny mean for flavonoid metabolites, total anthocyanins assessed by colourimetry and colour assessed visually.**

Mean phenotypic values from parents are represented in red for 'Capitola'. Ant, Fvo, F3ol values were obtained by summation of total anthocyanins, total flavonols and total flavan-3-ols, respectively; PgGs, Pelargonidin-3-glucoside; PgGsM, Pelargonidin-3-glucoside-malonate; PgRs, Pelargonidin-3-rutinoside; CyGs, Cyanidin-3-glucoside; AfPgGs, (epi)Afzelechin-pelargonidin-3-glucoside; KGs, Kaempferol-glucoside; KGn, Kaempferol-glucuronide; KCoGs, Kaempferol-coumaryl-glucoside; QGn, Quercetin-glucuronide; Cat, Catechin; CatCat, (epi)Catechin dimers; AfCat, (epi)Afzelechin-(epi)catechin dimers; AfGs, (epi)Afzelechin-glucoside; ANTHc, anthocyanins (colourimetry). The flavonoid metabolites values are expressed as mg-equ/100 g FW assuming a response factor of 1. ANTHc results are expressed as mg pelargonidin-3-glucoside equivalents/100 g FW. COLOUR values were assessed on a 0 to 6 scale. Values are the means of n = 3 replicates per genotype, except for COLOUR (n = 2).

#### A. QTLs on male linkage map

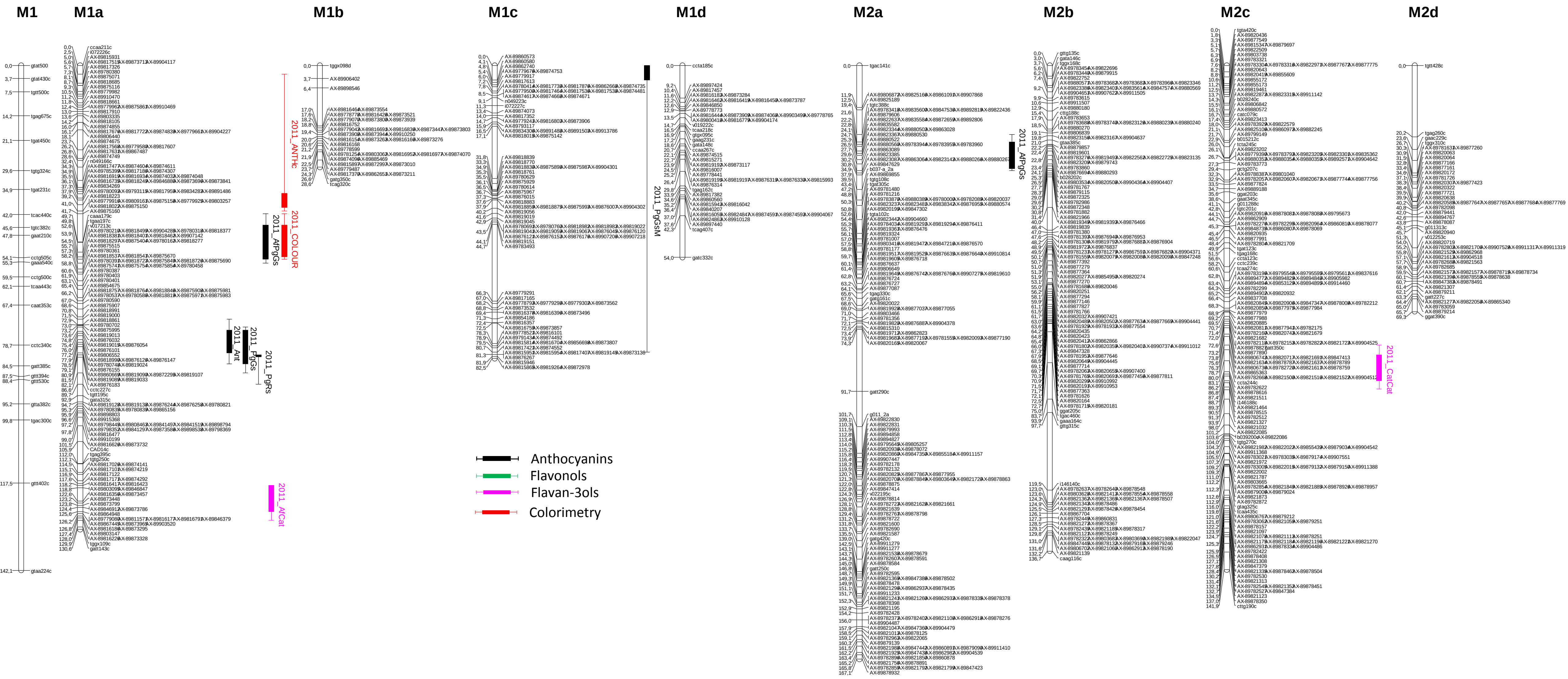


**Supplemental Figure 3. Localization of all detected QTLs on male (A) and female (B) linkage maps for flavonoid metabolites, total anthocyanins assessed by colourimetry and colour assessed visually**.
The QTLs for the various flavonoid metabolites, for the total anthocyanins measured by colorimetric assays or for colour assessed visually have a colour code that is shown on the bottom left of the figure. The QTL name is suffixed according to the year and the trait. The linkage groups are represented in MapChart 2.3 (R. E. Voorrips, 2002) with a space of 3mm by cM. Each boxplot corresponds to a QTL identified with a threshold of 10%. Bayesian credible interval of QTL is indicated at 5%. Ant, total anthocyanins; PgGs, Pelargonidin-3-glucoside; PgGsM, Pelargonidin-3-glucoside-malonate; PgRs, Pelargonidin-3-rutinoside; CyGs, Cyanidin-3-glucoside; AfPgGs, (epi)Afzelechin-pelargonidin-3-glucoside; Fvo, total flavonols; KGs, Kaempferol-glucoside; KGn, Kaempferol-glucuronide; KCoGs, Kaempferol-coumaryl-glucoside; QGn, Quercetin-glucuronide; F3ol, total flavan-3-ols; Cat, Catechin; CatCat, (epi)Catechin dimers; AfCat, (epi)Afzelechin-(epi)catechin dimers; AfGs, (epi)Afzelechin-glucoside; ANTHc, anthocyanins (colourimetry); COLOUR, colour (assessed visually).

### A. QTLs on male linkage map - continued

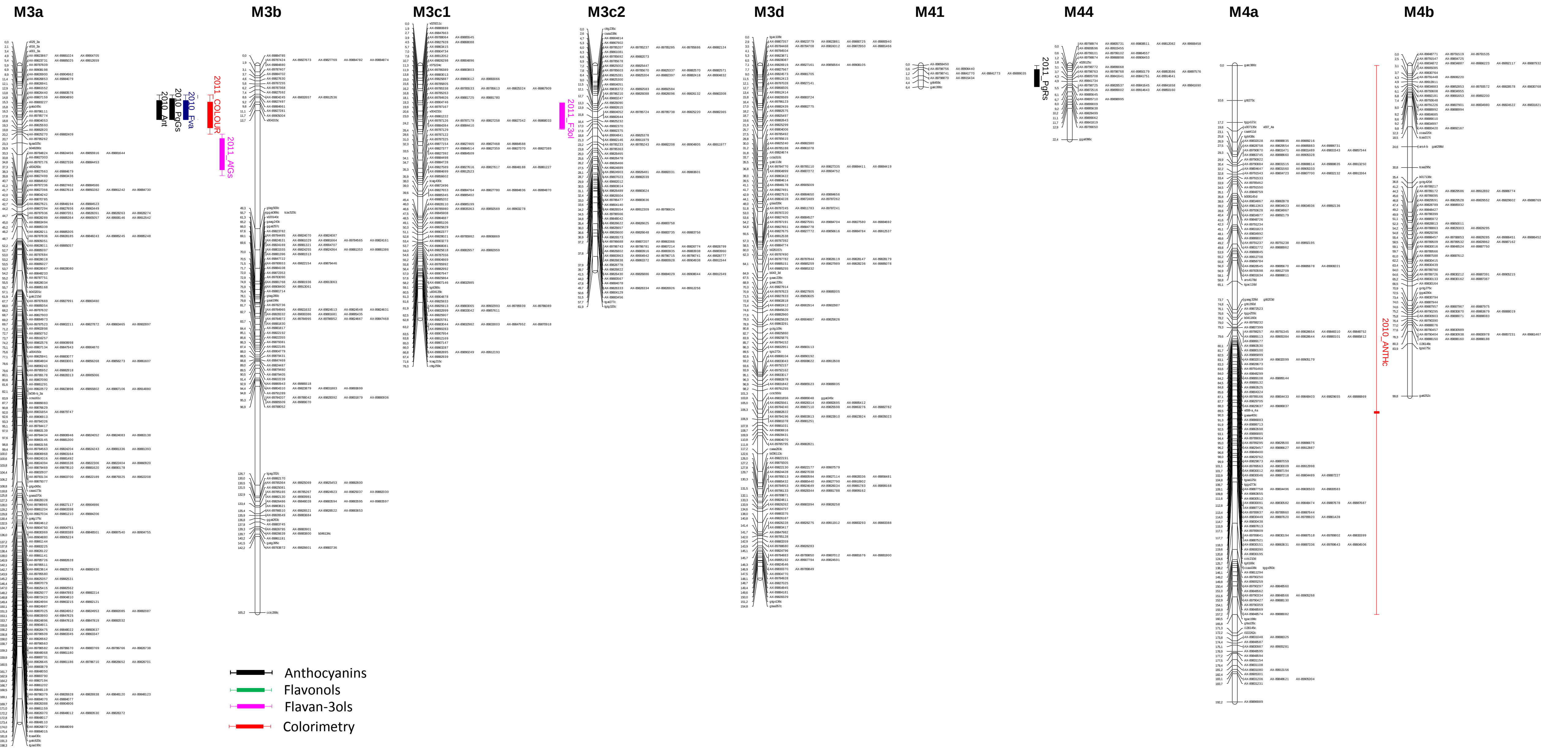

### A. QTLs on male linkage map - continued

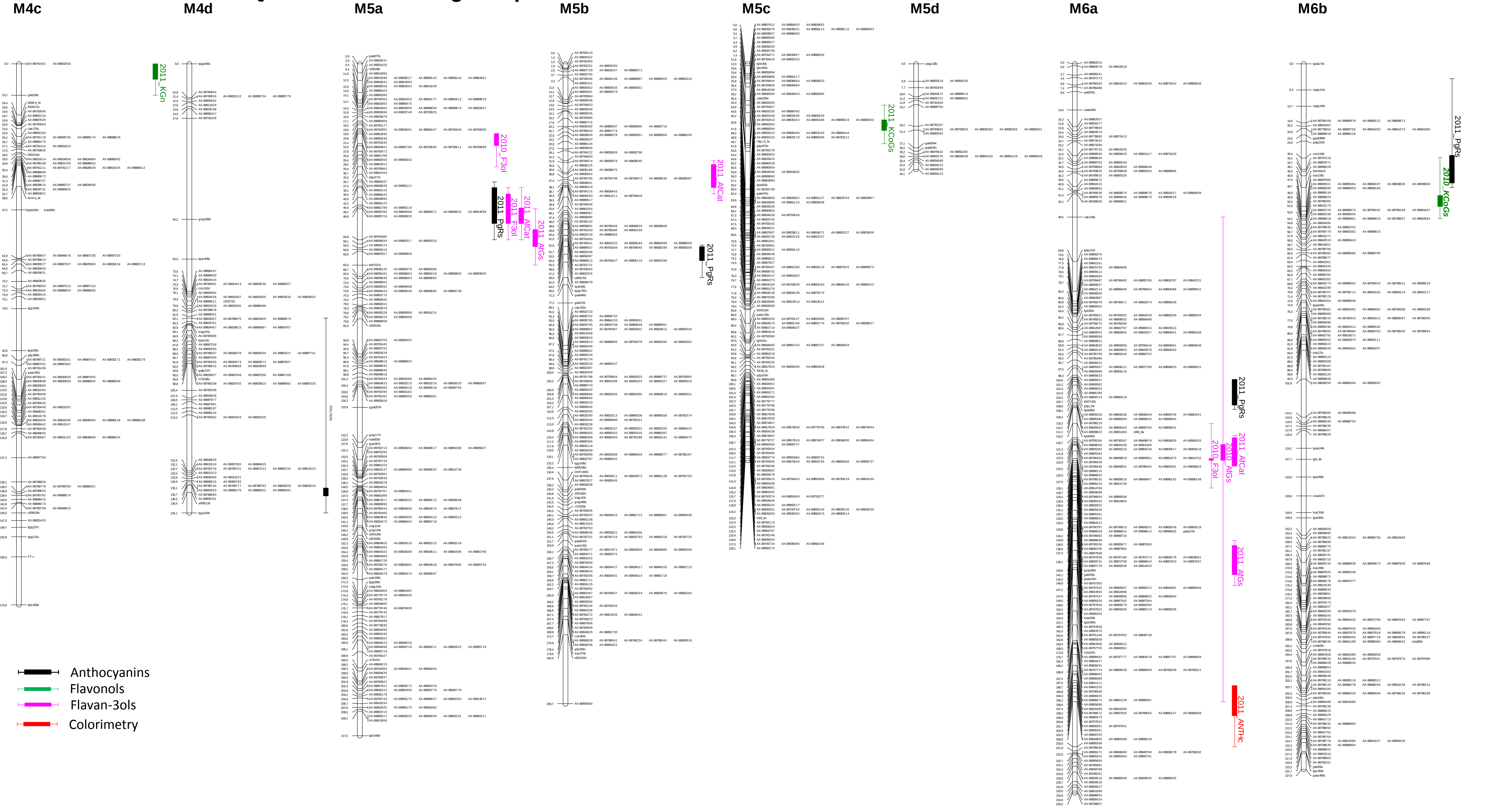

### A. QTLs on male linkage map - continued

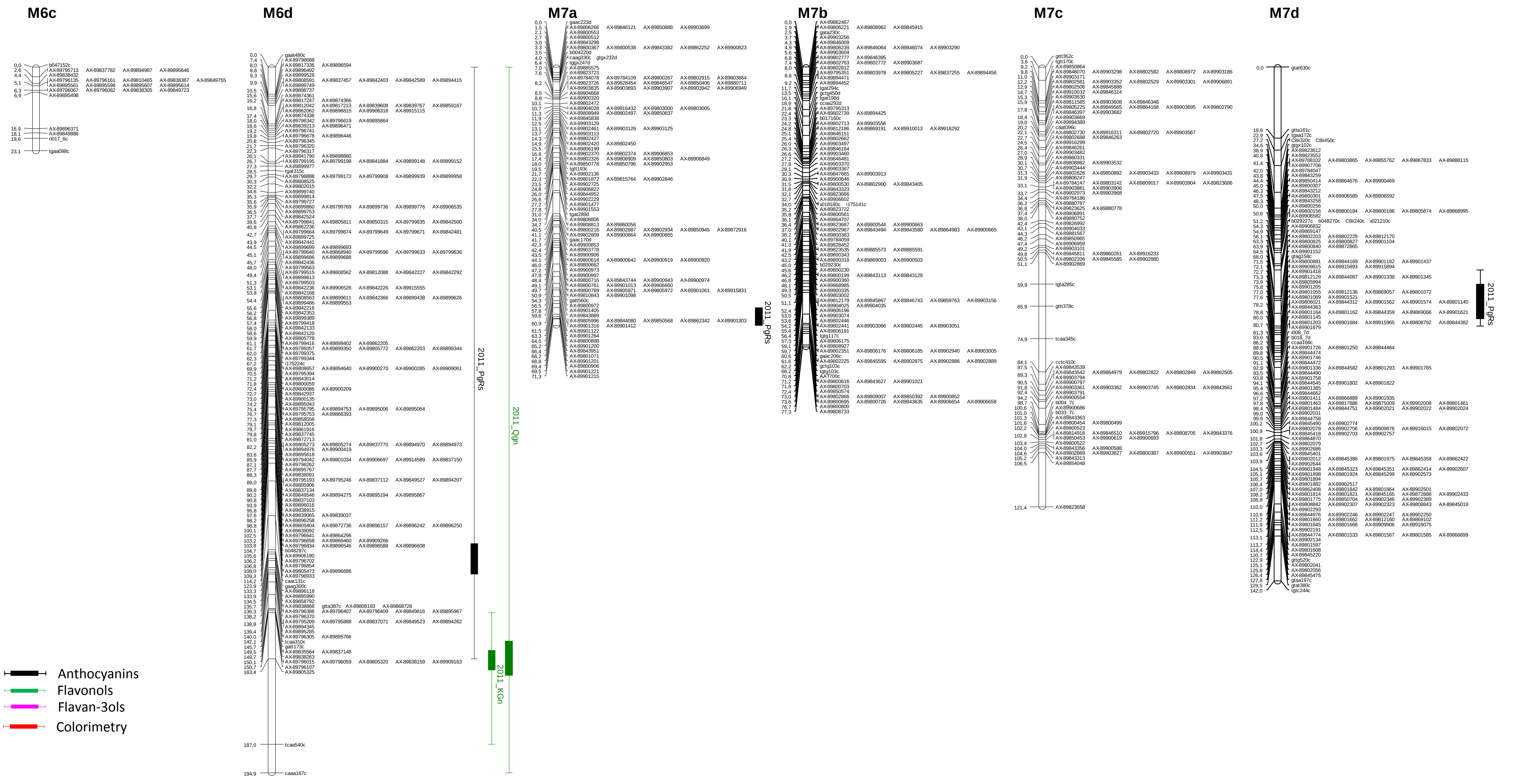

### B. QTLs on female linkage map

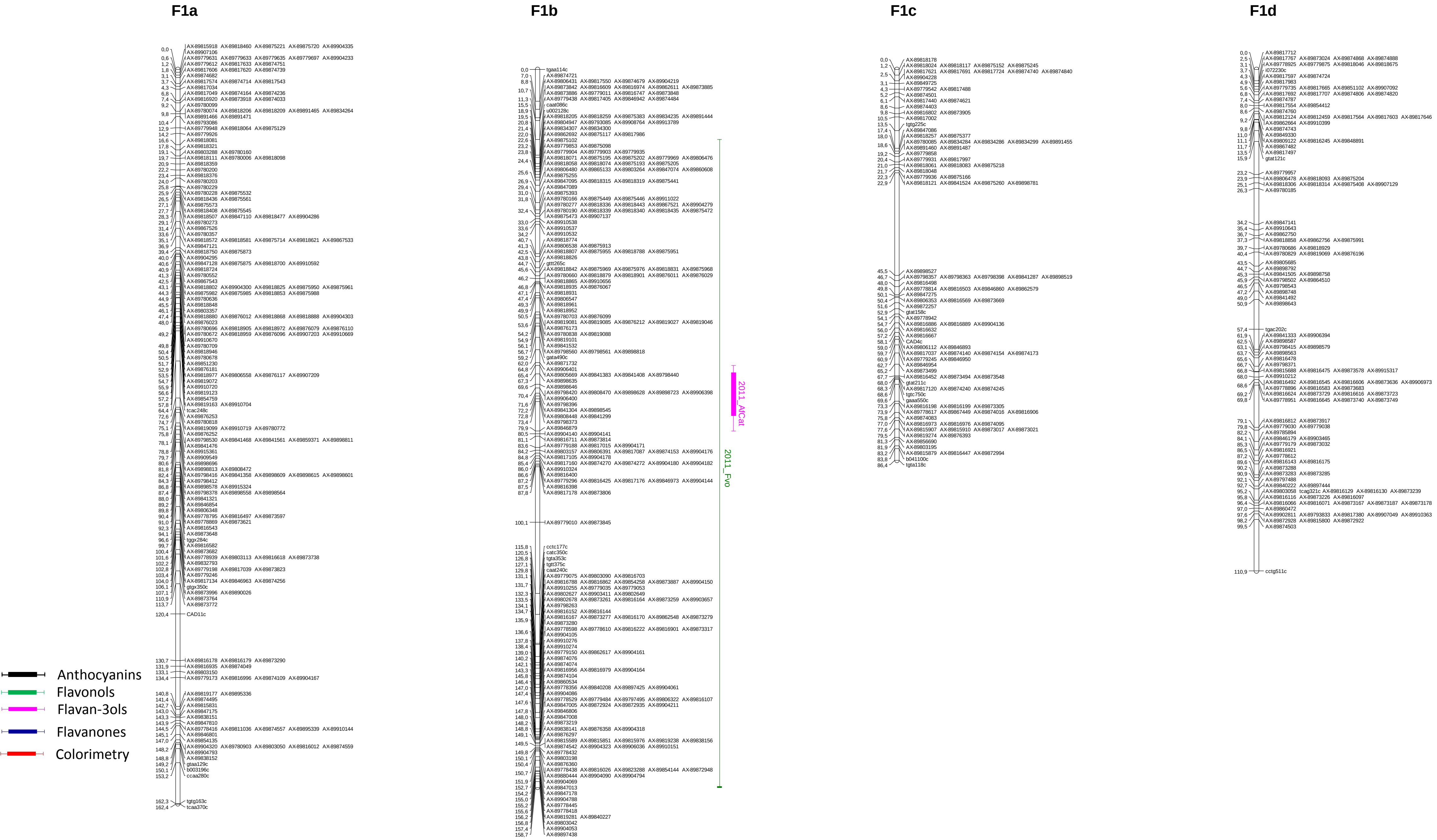

### **B. QTLs on female linkage map** - continued

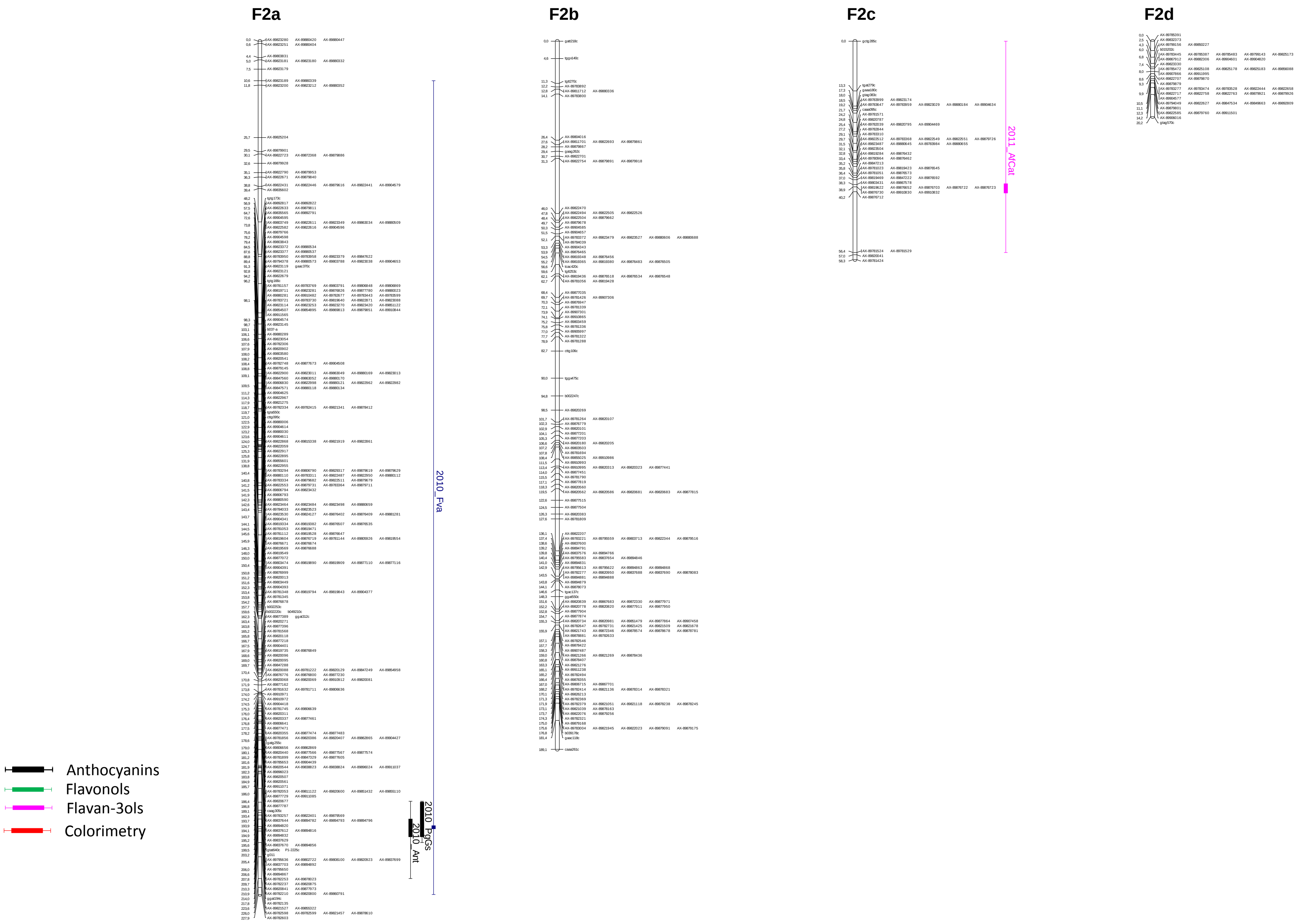

### B. QTLs on female linkage map - continued

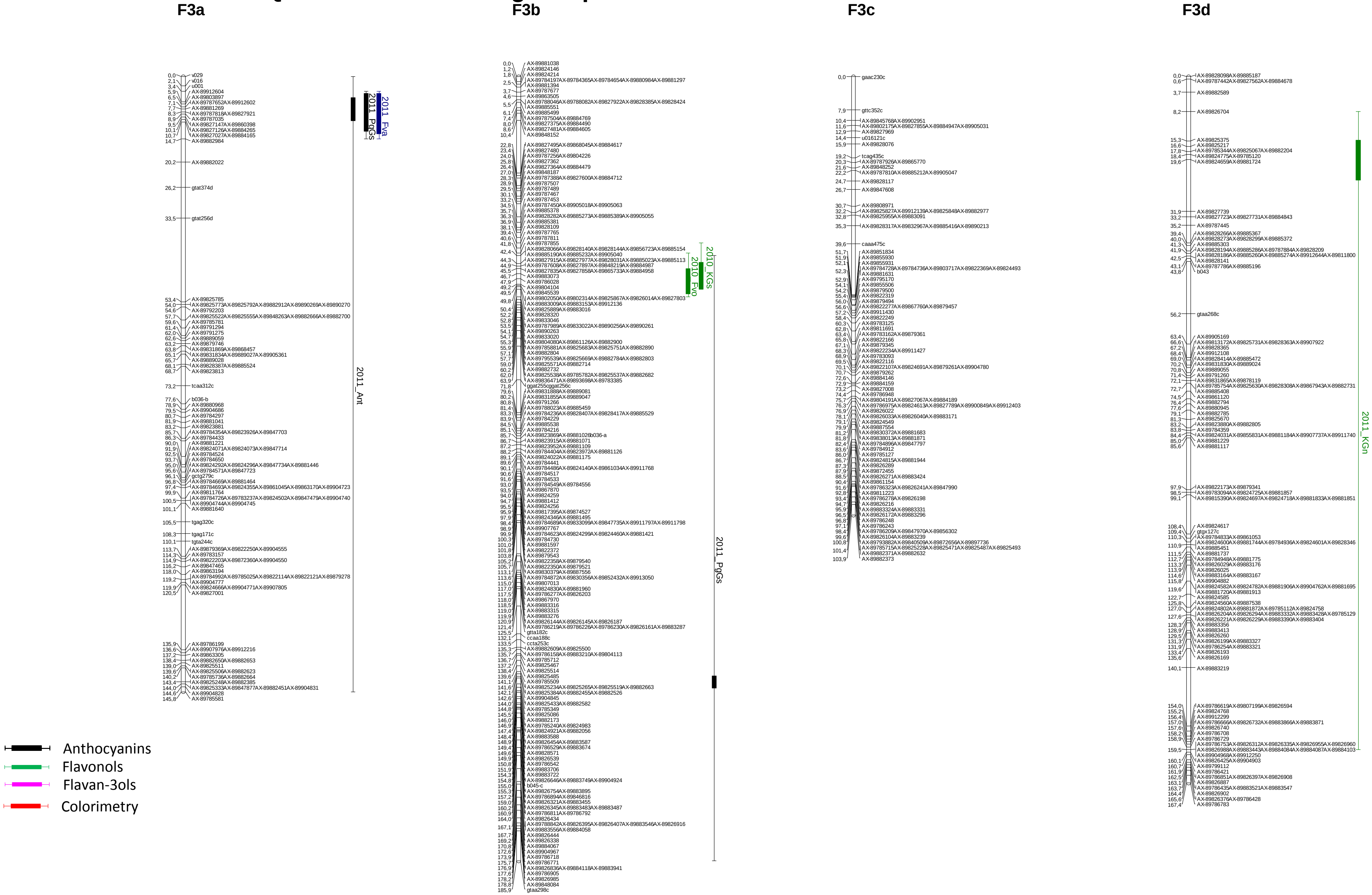

### B. QTLs on female linkage map - continued

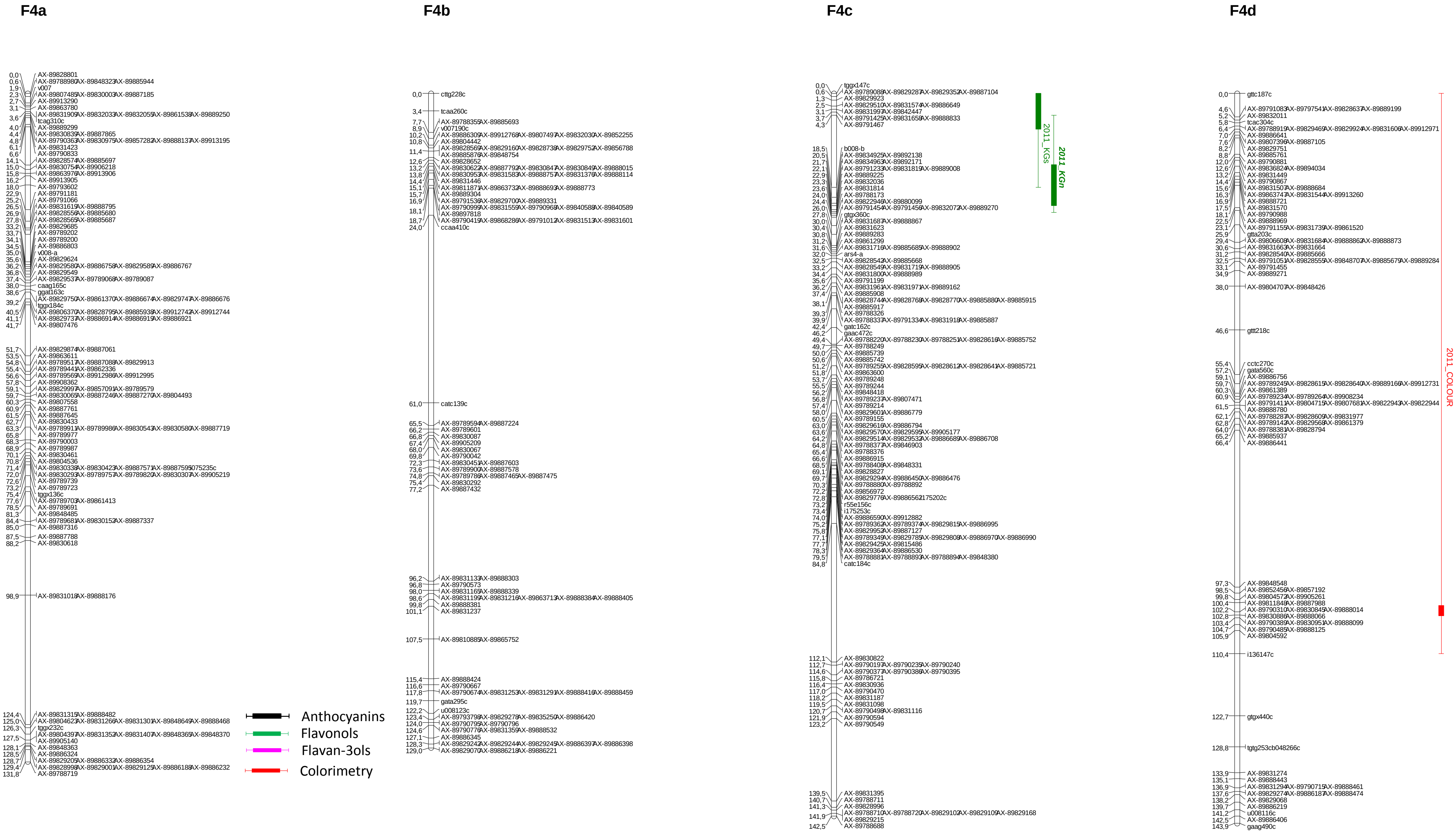

### B. QTLs on female linkage map - continued

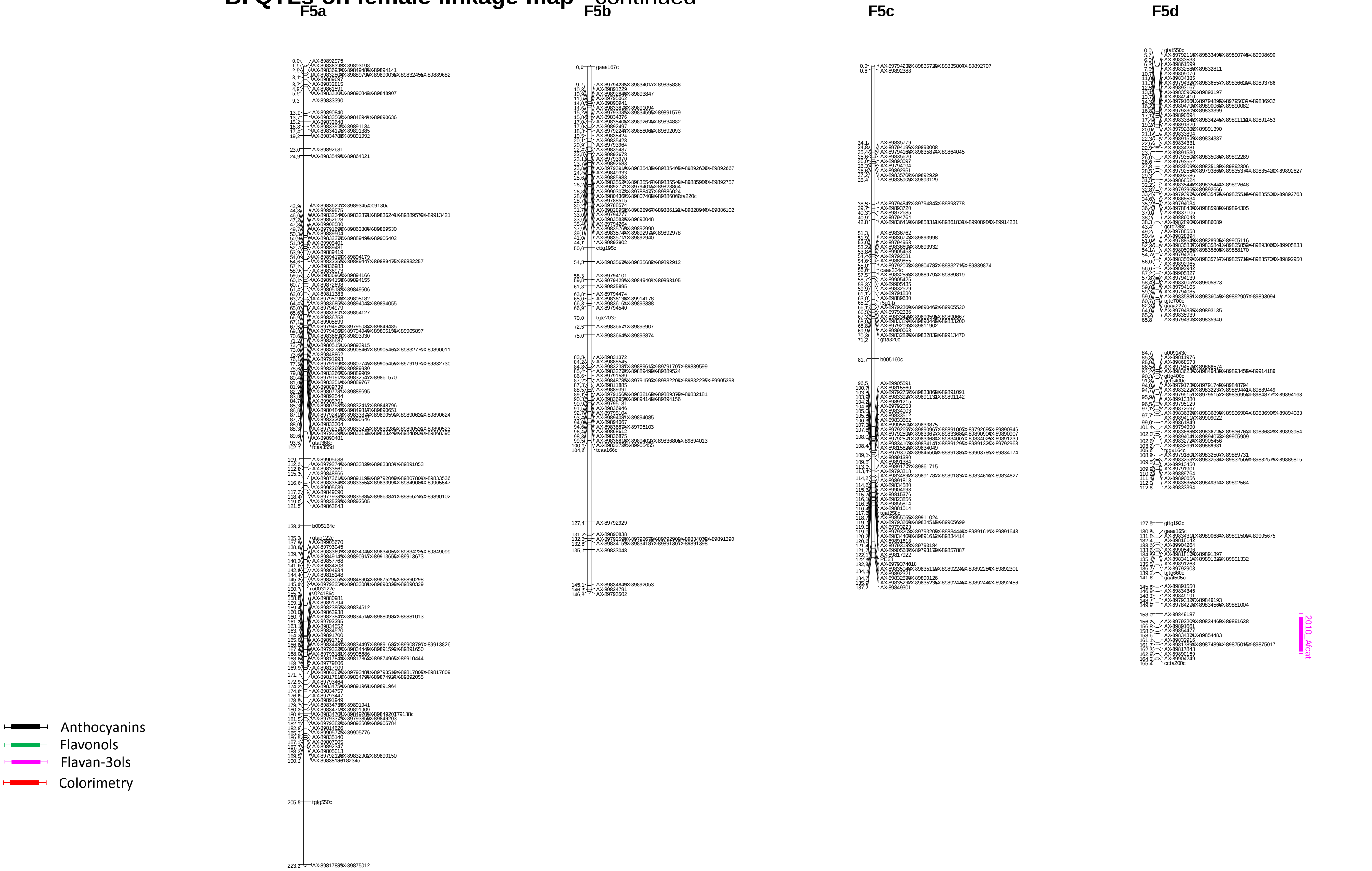

### B. QTLs on female linkage map - continued

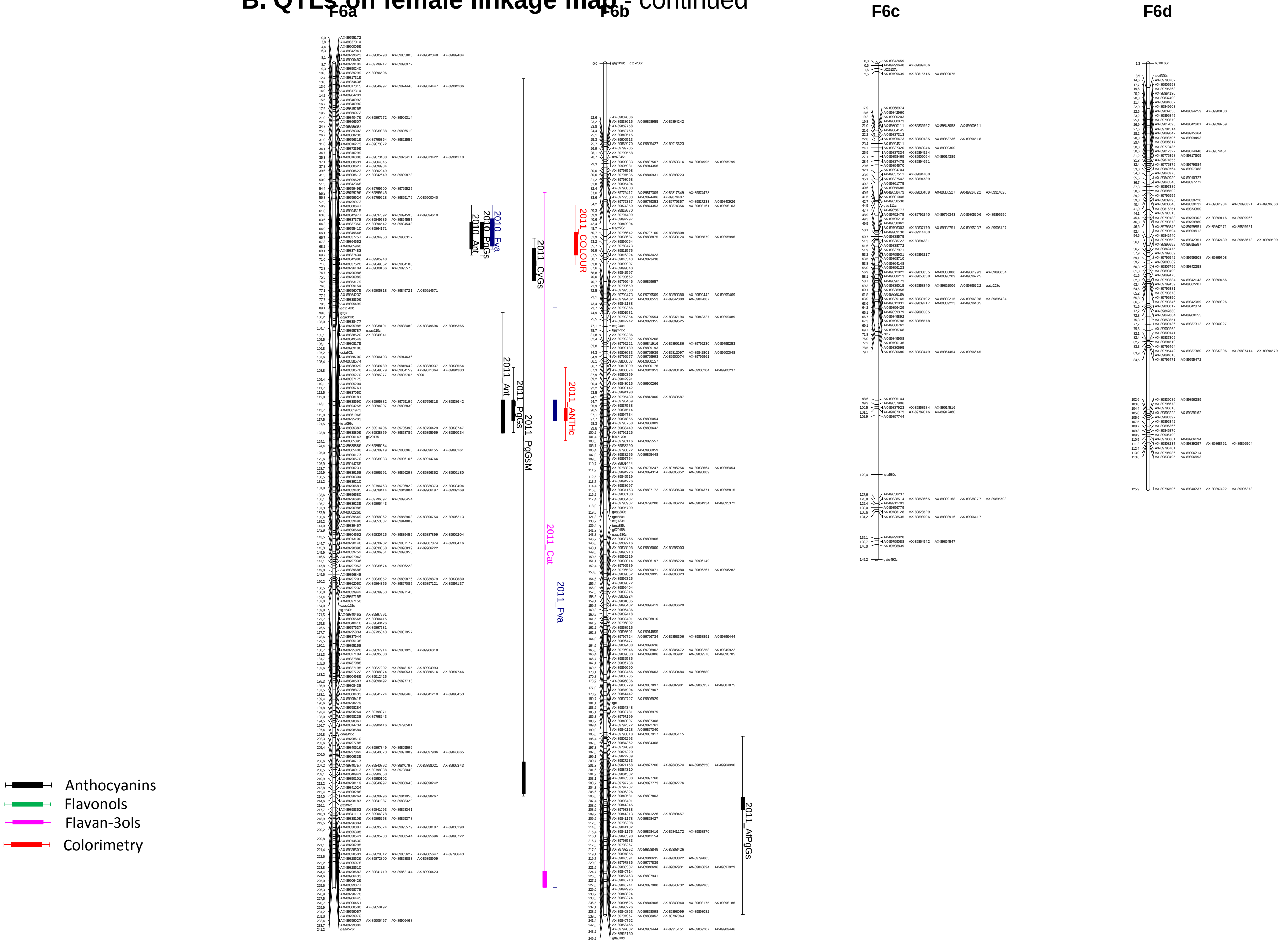

### **B. QTLs on female linkage map** - continued

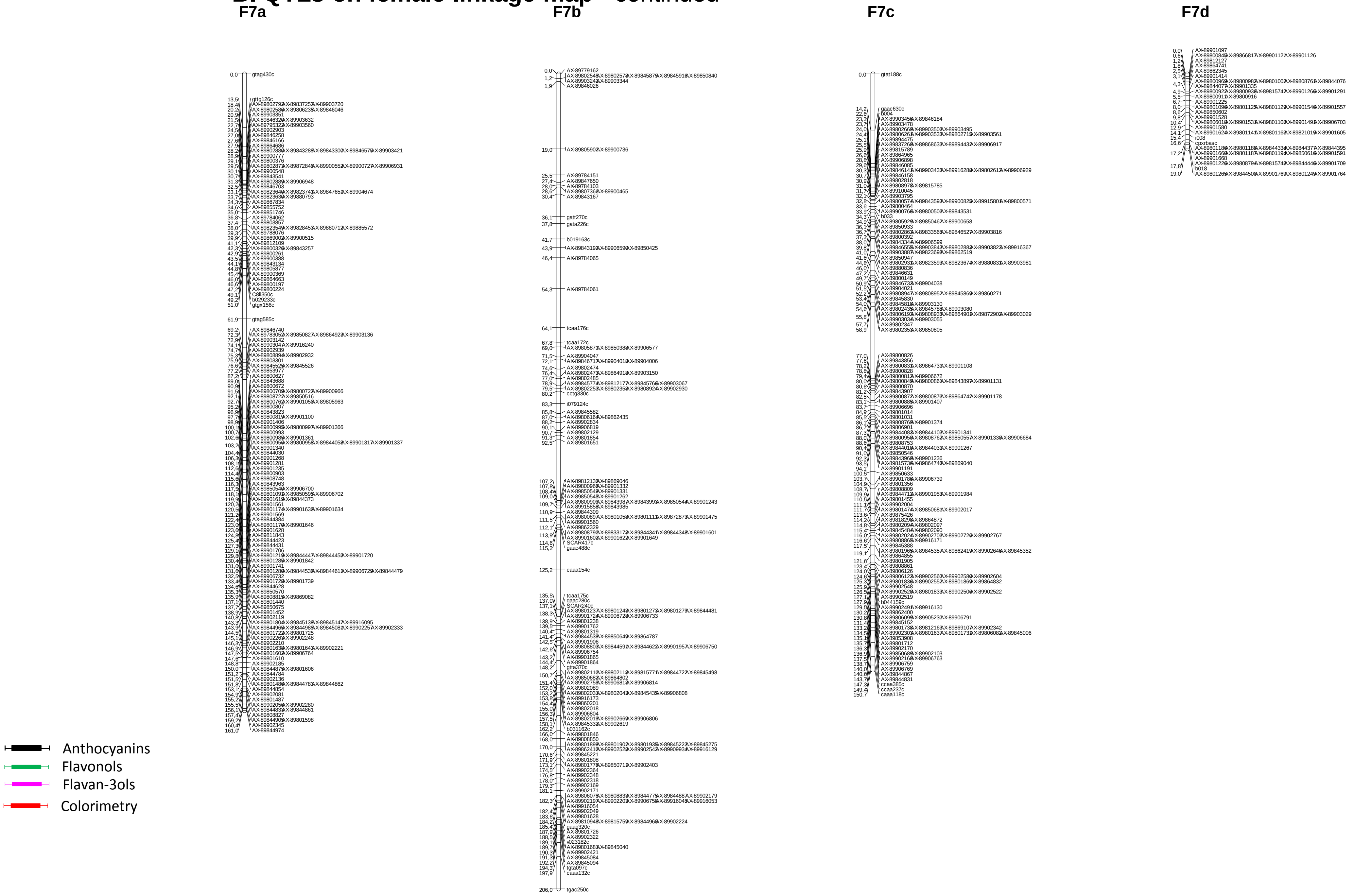
