## Supplemental Table 1 to 9 for "The genetic architecture of fruit colour in strawberry (*Fragaria × ananassa*) uncovers the predominant contribution of the *F. vesca* subgenome to anthocyanins and reveals underlying genetic variations"

**Supplemental Table 1.** COLOUR trait values for ‘Capitola’ and ‘Capitola’ x ‘CF1116’ progeny in 2011.

| Traits | Abbr. | Capitola | | | CF1116 | | | | Progeny | | |
|---|---|---|---|---|---|---|---|---|---|---|---|
| | | mean | s.d. | range | mean | s.d. | range | Cap vs CF | mean | s.d. | range |
| Colour | COLOUR | 3.50 | 0.00 | 3.50-3.50 | NA | NA | NA | NA | 3.38 | 1.33 | 0.50-6.00 |

The mean value, standard deviation (s.d.) and range are described. ‘CF1116’ was not studied in 2011.
COLOUR values were assessed on a 0 to 6 scale. Values are the means of n = 3 replicates per genotype.

**Supplemental Table 2.** Correspondence between names of linkage groups from female ('Capitola') and male ('CF1116') parents and the *F. vesca*, *F. viridis*, *F. iinumae* and *F. nipponica* subgenomes according to positions of SNP markers from Axiom 90K SNP array (Whitaker and Verma, unpublished results) on the *F. x ananassa* reference genome (Edger et al., 2019).

| Female | *Affyx-for-correspondancy-Camarosa-Genome* | *Camarosa genome* | | affyx | gene | rapd | scar | ssr | Total markers | LG size (cM) |
|---|---|---|---|---|---|---|---|---|---|---|
| F1a | *9* | *Fvb1-2* | *F. iinumae* | 214 | 1 | 7 | | 1 | 223 | 162.42 |
| F1b | *36* | *Fvb1-4* | *F. vesca* | 269 | | 9 | | 1 | 279 | 158.67 |
| F1c | *18* | *Fvb1-3* | *F. nipponica* | 104 | 1 | 6 | | 1 | 112 | 86.39 |
| F1d | *7* | *Fvb1-1* | *F. viridis* | 138 | | 4 | | 1 | 143 | 110.87 |
| F2a | *4* | *Fvb2-2* | *F. vesca* | 329 | | 10 | | 6 | 345 | 227.93 |
| F2b | *3* | *Fvb2-3* | *F. viridis* | 174 | | 12 | | 2 | 188 | 189.11 |
| F2c | *0* | *Fvb2-1* | *F. nipponica* | 52 | | 5 | | | 57 | 58.29 |
| F2d | *12* | *Fvb2-1* | *F. nipponica* | 45 | | 1 | | 1 | 47 | 20.15 |
| F3a | *2* | *Fvb3-4* | *F. vesca* | 117 | | 7 | | 4 | 128 | 145.80 |
| F3b | *32* | *Fvb3-1* | *F. viridis* | 274 | | 6 | | 2 | 282 | 185.90 |
| F3c | *5* | *Fvb3-3* | *F. nipponica* | 124 | | 4 | | 1 | 129 | 103.93 |
| F3d | *13* | *Fvb3-2* | *F. iinumae* | 175 | | 2 | | 1 | 178 | 167.43 |
| F4a | *0* | *Fvb4-3* | *F. vesca* | 148 | | 6 | | 3 | 157 | 131.83 |
| F4b | *0* | *Fvb4-4* | *F. iinumae* | 101 | | 6 | | 1 | 108 | 128.95 |
| F4c | *12* | *Fvb4-2* | *F. nipponica* | 156 | | 5 | | 5 | 166 | 142.54 |
| F4d | *12* | *Fvb4-1* | *F. viridis* | 104 | | 9 | | 3 | 116 | 143.86 |
| F5a | *12* | *Fvb5-1* | *F. vesca* | 264 | | 4 | | 6 | 274 | 223.16 |
| F5b | *14* | *Fvb5-4* | *F. nipponica* | 142 | | 5 | | | 147 | 146.92 |
| F5c | *9* | *Fvb5-2* | *F. viridis* | 162 | | 3 | | 4 | 169 | 137.19 |
| F5d | *9* | *Fvb5-3* | *F. iinumae* | 208 | | 12 | | 1 | 221 | 165.44 |
| F6a | *40* | *Fvb6-1* | *F. vesca* | 404 | | 11 | | 2 | 417 | 241.22 |
| F6b | *21* | *Fvb6-4* | *F. viridis* | 349 | | 12 | | 3 | 364 | 249.23 |
| F6c | *20* | *Fvb6-3* | *F. iinumae* | 139 | | 4 | | 2 | 145 | 145.18 |
| F6d | *8* | *Fvb6-2* | *F. nipponica* | 139 | | 1 | | 1 | 141 | 125.90 |
| F7a | *46* | *Fvb7-3* | *F. iinumae* | 194 | | 4 | | 2 | 200 | 161.04 |
| F7b | *6* | *Fvb7-4* | *F. viridis* | 174 | | 14 | 2 | 4 | 194 | 206.00 |
| F7c | *38* | *Fvb7-2* | *F. vesca* | 212 | | 5 | | 3 | 220 | 150.74 |
| F7d | *9* | *Fvb7-1* | *F. nipponica* | 63 | | | 1 | 2 | 66 | 18.98 |
| | | Total | | 4974 | 2 | 174 | 3 | 63 | 5216 | 4135.10 |

| Male | *Affyx-for-correspondancy-Camarosa-Genome* | *Camarosa genome* | | affyx | gene | rapd | scar | ssr | Total markers | LG size (cM) |
|---|---|---|---|---|---|---|---|---|---|---|
| M1a | *7* | *Fvb1-2* | *F. iinumae* | 184 | 1 | 10 | | 3 | 198 | 130.61 |

| | | | | | | | | | | |
|---|---|---|---|---|---|---|---|---|---|---|
| M1b | *4* | *Fvb1-4* | *F. vesca* | 39 | | 3 | | | 42 | 28.61 |
| M1c | *17* | *Fvb1-3* | *F. nipponica* | 104 | | | | 2 | 106 | 82.53 |
| M1d | *2* | *Fvb1-1* | *F. viridis* | 43 | | 9 | | 1 | 53 | 54.01 |
| M2a | *5* | *Fvb2-2* | *F. vesca* | 194 | | 10 | | 3 | 207 | 167.07 |
| M2b | *24* | *Fvb2-3* | *F. viridis* | 174 | | 10 | | 2 | 186 | 136.65 |
| M2c | *8* | *Fvb2-1* | *F. nipponica* | 200 | | 18 | | 5 | 223 | 141.88 |
| M2d | *9* | *Fvb2-4* | *F. iinumae* | 52 | | 6 | | 2 | 60 | 69.27 |
| M3a | *23* | *Fvb3-4* | *F. vesca* | 271 | | 11 | | 8 | 290 | 198.29 |
| M3b | *2* | *Fvb3-1* | *F. viridis* | 129 | | 11 | | 3 | 143 | 165.15 |
| M3c1&c2 | *8* | *Fvb3-3* | *F. nipponica* | 248 | | 8 | | 4 | 260 | 138.18 |
| M3d | *23* | *Fvb3-2* | *F. iinumae* | 216 | | 13 | | 3 | 232 | 154.77 |
| M4a | *13* | *Fvb4-3* | *F. vesca* | 191 | | 19 | | 8 | 218 | 192.23 |
| M4b | *0* | *Fvb4-4* | *F. iinumae* | 110 | | 9 | | 3 | 122 | 99.82 |
| M4c | *5* | *Fvb4-2* | *F. nipponica* | 107 | 1 | 12 | | 4 | 124 | 174.78 |
| M4d | *3* | *Fvb4-1* | *F. viridis* | 97 | | 8 | | 3 | 108 | 145.08 |
| M5a | *29* | *Fvb5-1* | *F. vesca* | 247 | | 12 | | 6 | 265 | 217.01 |
| M5b | *26* | *Fvb5-4* | *F. nipponica* | 257 | 1 | 14 | | 5 | 277 | 206.74 |
| M5c | *31* | *Fvb5-2* | *F. viridis* | 210 | | 9 | | 4 | 223 | 128.09 |
| M5d | *0* | *Fvb5-3* | *F. iinumae* | 27 | | 3 | | | 30 | 34.12 |
| M6a | *32* | *Fvb6-1* | *F. vesca* | 312 | | 15 | | 2 | 329 | 235.46 |
| M6b | *26* | *Fvb6-4* | *F. viridis* | 222 | | 22 | | 1 | 245 | 227.90 |
| M6c | *1* | *Fvb6-3* | *F. iinumae* | 21 | | 1 | | 2 | 24 | 23.11 |
| M6d | *2* | *Fvb6-2* | *F. nipponica* | 230 | | 9 | | 2 | 241 | 194.93 |
| M7a | *17* | *Fvb7-3* | *F. iinumae* | 124 | | 8 | | 1 | 133 | 71.34 |
| M7b | *18* | *Fvb7-4* | *F. viridis* | 122 | | 9 | | 5 | 136 | 77.33 |
| M7c | *10* | *Fvb7-2* | *F. vesca* | 115 | | 7 | | 2 | 124 | 121.35 |
| M7d | *13* | *Fvb7-1* | *F. nipponica* | 183 | | 10 | | 8 | 201 | 142.01 |
| | | Sub-Total | | 4429 | 3 | 276 | 0 | 92 | 4800 | |
| Small LGs | | | | | | | | | | |
| M41 | | | | 9 | | 2 | | | 11 | 6.41 |
| M44 | 3 | Fvb6-2 | *F. nipponica* | 43 | | 1 | | 1 | 45 | 22.41 |
| M1 | | | | | | 23 | | | 23 | 142.06 |
| | | Total | | 4481 | 3 | 302 | 0 | 93 | 4879 | |

**Supplemental Table 3.** Summary of male linkage map.
For each linkage group (LG), number of markers, length, mean spacing between markers and max spacing are indicated.

| LG | Nb of Markers | length (cM) | mean spacing (cM) | max spacing (cM) |
|---|---|---|---|---|
| M1 | 23 | 142.1 | 6.5 | 24.6 |
| M1a | 198 | 130.6 | 0.7 | 8.1 |
| M1b | 42 | 28.6 | 0.7 | 10.6 |
| M1c | 106 | 82.5 | 0.8 | 21.7 |
| M1d | 53 | 54 | 1 | 11.8 |
| M2a | 207 | 167.1 | 0.8 | 17.4 |
| M2b | 186 | 136.6 | 0.7 | 21.8 |
| M2c | 223 | 141.9 | 0.6 | 5.1 |
| M2d | 60 | 69.3 | 1.2 | 20.2 |
| M3a | 290 | 198.3 | 0.7 | 10 |
| M3b | 143 | 165.2 | 1.2 | 35.6 |
| M3c1 | 131 | 76.3 | 0.6 | 5.8 |
| M3c2 | 129 | 61.9 | 0.5 | 6.2 |
| M3d | 232 | 154.8 | 0.7 | 5.4 |
| M41 | 11 | 6.4 | 0.6 | 2 |
| M44 | 45 | 22.4 | 0.5 | 9.5 |
| M4a | 218 | 192.2 | 0.9 | 12.6 |
| M4b | 122 | 99.8 | 0.8 | 15.9 |
| M4c | 124 | 174.8 | 1.4 | 15.6 |
| M4d | 108 | 145.1 | 1.4 | 32.7 |
| M5a | 265 | 217 | 0.8 | 11.5 |
| M5b | 277 | 206.7 | 0.7 | 22.3 |
| M5c | 223 | 128.1 | 0.6 | 12.2 |
| LG5d | 30 | 34.1 | 1.2 | 7 |
| LG6a | 329 | 235.5 | 0.7 | 14.6 |
| LG6b | 245 | 227.9 | 0.9 | 12.3 |
| LG6c | 24 | 23.1 | 1 | 10 |
| LG6d | 241 | 194.9 | 0.8 | 23.5 |
| LG7a | 133 | 71.3 | 0.5 | 5.8 |
| LG7b | 136 | 77.3 | 0.6 | 6 |
| LG7c | 124 | 121.4 | 1 | 14.9 |
| M7d | 201 | 142 | 0.7 | 19.6 |
| Total or Mean | 4879 | 3929.2 | 0.8 | 35.6 |

**Supplemental Table 4.** Summary of female linkage map.
For each linkage group (LG), number of markers, length, mean spacing between marker and max spacing are indicated.

| LG | Nb of Markers | length (cM) | mean spacing (cM) | max spacing (cM) |
|---|---|---|---|---|
| F1a | 223 | 162.4 | 0.7 | 10.3 |
| F1b | 279 | 158.7 | 0.6 | 15.7 |
| F1c | 112 | 86.4 | 0.8 | 22.6 |
| F1d | 143 | 110.9 | 0.8 | 11.4 |
| F2a | 345 | 227.9 | 0.7 | 13.9 |
| F2b | 188 | 189.1 | 1 | 14.7 |
| F2c | 57 | 58.3 | 1 | 16.2 |
| F2d | 47 | 20.2 | 0.4 | 6 |
| F3a | 128 | 145.8 | 1.1 | 19.9 |
| F3b | 282 | 185.9 | 0.7 | 12.3 |
| F3c | 129 | 103.9 | 0.8 | 12.1 |
| F3d | 178 | 167.4 | 0.9 | 13.9 |
| F4a | 157 | 131.8 | 0.8 | 25.5 |
| F4b | 108 | 129 | 1.2 | 37 |
| F4c | 166 | 142.5 | 0.9 | 27.3 |
| F4d | 116 | 143.9 | 1.3 | 30.9 |
| F5a | 274 | 223.2 | 0.8 | 18.1 |
| F5b | 147 | 146.9 | 1 | 22.8 |
| F5c | 169 | 137.2 | 0.8 | 23.5 |
| F5d | 221 | 165.4 | 0.8 | 18.9 |
| F6a | 417 | 241.2 | 0.6 | 15.8 |
| F6b | 364 | 249.2 | 0.7 | 22.6 |
| F6c | 145 | 145.2 | 1 | 18.9 |
| F6d | 141 | 124.6 | 0.9 | 18.1 |
| F7a | 200 | 161 | 0.8 | 13.5 |
| F7b | 194 | 206 | 1.1 | 17.2 |
| F7c | 220 | 150.7 | 0.7 | 18.1 |
| F7d | 66 | 19 | 0.3 | 2.5 |
| Total or Mean | 5216 | 4133.8 | 0.8 | 37 |

**Supplemental Table 5.** Significant QTLs detected in male and female linkage maps for all traits and for two years based on CIM analysis with LOD> LOD threshold 10% for each trait.
For each trait, the Abbreviation (Abbr.), years, linkage groups (LGs), number of covariable (nb cov) for CIM QTL analyses, QTLs Marker names (Marker) and positions (pos.), LOD score (LOD), effect and $R^2$, and flanking QTL markers names (Left and Right) and positions (pos.) are indicated.

| Traits | Abbr. | years | LGs | nb cov | Marker | Pos. | LOD | Effect | $R^2$ | Left flanking marker | Pos. | Right flanking marker | Pos. |
|---|---|---|---|---|---|---|---|---|---|---|---|---|---|
| **Anthocyanins** | | | | | | | | | | | | | |
| Total anthocyanins | Ant | 2011 | M1a | 1 | AX-89780748 | 78.46 | 4.42 | 3.97 | 14.38 | AX-89819000 | 71.52 | AX-89860669 | 80.90 |
| | | 2010 | F2a | 2 | AX-89782237 | 209.65 | 3.87 | -4.44 | 14.94 | AX-89878023 | 207.80 | AX-89821527 | 223.56 |
| | | 2010 | M3a | 1 | AX-89826853 | 11.37 | 3.27 | 4.98 | 18.89 | AX-89904962 | 8.86 | AX-89786111 | 16.29 |
| | | 2011 | F3a | 2 | AX-89787035 | 8.89 | 2.66 | -3.15 | 8.77 | v029_3a | 0.00 | AX-89785581 | 145.80 |
| | | 2010 | F6a | 2 | cF6a.loc49.5 | 49.50 | 5.02 | -5.45 | 22.17 | AX-89899878 | 41.45 | AX-89799499 | 54.38 |
| | | 2011 | F6a | 2 | cF6a.loc101 | 101.00 | 2.99 | -3.40 | 10.49 | AX-89895575 | 72.81 | AX-89838574 | 108.42 |
| Pelargonidin-3-glucoside | PgGs | 2011 | M1a | 2 | AX-89780748 | 78.46 | 4.34 | 2.98 | 11.65 | AX-89819013 | 73.56 | cctc227c | 86.58 |
| | | 2010 | F2a | 2 | cF2a.loc208.5 | 208.50 | 4.07 | -4.22 | 14.98 | g011_2a | 203.19 | ggat194c | 214.00 |
| | | 2010 | M3a | 1 | AX-89826853 | 11.37 | 3.60 | 4.89 | 20.59 | AX-89904962 | 8.86 | AX-89786111 | 16.29 |
| | | 2011 | F3a | 4 | AX-89787035 | 8.89 | 3.43 | -2.58 | 8.46 | u001_3a | 3.40 | AX-89882984 | 14.74 |
| | | 2011 | F3b | 4 | cF3b.loc43.5 | 143.50 | 2.70 | -2.56 | 8.39 | AX-89905040 | 42.42 | gtaa298c | 185.90 |
| | | 2011 | M4d | 2 | u008113c | 138.35 | 3.05 | -2.43 | 7.56 | AX-89886574 | 82.08 | tggx104c | 145.08 |
| | | 2010 | F6a | 2 | cF6a.loc49.5 | 49.50 | 4.47 | -4.84 | 19.70 | AX-89899878 | 41.45 | AX-89799499 | 54.38 |
| | | 2011 | F6a | 4 | cF6a.loc103.5 | 103.50 | 4.66 | -2.66 | 9.26 | gctg280c | 89.11 | gaaa610c | 104.70 |
| Pelargonidin-3-glucoside-malonate | PgGsM | 2011 | M1c | 1 | AX-89860573 | 0.00 | 1.93 | -0.18 | 6.54 | AX-89860573 | 0.00 | AX-89874552 | 80.70 |
| | | | F6a | 1 | AX-89840941 | 209.07 | 2.65 | 0.20 | 8.91 | AX-89900059 | 4.41 | AX-89798187 | 214.58 |
| Pelargonidin-3-rutinoside | PgRs | 2011 | M1a | 9 | cctc227c | 86.58 | 8.29 | 0.77 | 12.40 | AX-89819033 | 81.53 | tgtt195c | 89.71 |
| | | | M41 | 9 | gatc390c | 6.41 | 5.26 | -0.49 | 5.46 | AX-89899035 | 1.24 | gatc390c | 6.41 |
| | | | M5a | 9 | cM5a.loc48.5 | 48.50 | 6.61 | 0.65 | 9.53 | AX-89828890 | 39.96 | AX-89794385 | 56.84 |
| | | | M5b | 9 | tgat148c | 63.97 | 4.54 | -0.43 | 4.15 | u009170c | 58.96 | tgag700c | 68.97 |
| | | | M6a | 9 | cM6a.loc104 | 104.00 | 4.76 | 0.60 | 8.20 | AX-89895789 | 101.85 | AX-89796219 | 110.22 |
| | | | M6b | 9 | AX-89797216 | 34.09 | 3.85 | -0.61 | 8.16 | cctc218d | 30.42 | b010162d | 40.02 |
| | | | M6d | 9 | AX-89796386 | 136.33 | 3.36 | -0.71 | 11.54 | gaat480c | 0.00 | AX-89805325 | 163.43 |
| | | | M7a | 9 | AX-89901215 | 71.34 | 6.24 | -0.48 | 4.90 | AX-89843951 | 66.37 | AX-89901215 | 71.34 |
| | | | M7d | 9 | AX-89901152 | 64.51 | 4.16 | 0.32 | 2.32 | AX-89812170 | 56.13 | AX-89800881 | 71.47 |
| Cyanidin-3-glucoside | CyGs | 2011 | F6a | 1 | AX-89808647 | 58.92 | 4.36 | -0.13 | 14.20 | AX-89842368 | 51.25 | AX-89894557 | 63.63 |
| (epi)Afzelechin-pelargonidin-glucoside | AfPgGs | 2011 | M1a | 2 | tgaa197c | 49.77 | 3.74 | 0.10 | 12.73 | AX-89875160 | 41.65 | AX-89875515 | 55.71 |
| | | | M2a | 2 | AX-89823367 | 24.66 | 4.69 | 0.11 | 15.19 | AX-89783418 | 21.61 | AX-89823385 | 29.58 |
| | | | F6b | 1 | AX-89798583 | 216.66 | 2.73 | 0.08 | 8.17 | AX-89884362 | 196.99 | gtta310d | 249.23 |
| **Flavonols** | | | | | | | | | | | | | |
| Total Flavonols | Fvo | 2011 | F1b | 1 | AX-89804947 | 20.75 | 2.42 | -0.03 | 8.16 | caat086c | 15.45 | AX-89897438 | 158.67 |
| | | 2010 | F3b | 1 | AX-89827835 | 45.49 | 2.83 | -0.07 | 16.54 | AX-89787855 | 41.81 | AX-89828320 | 52.24 |
| Kaempferol-glucoside | KGs | 2010 | F3b | 1 | AX-89827835 | 45.49 | 3.01 | -0.03 | 17.50 | AX-89787765 | 39.35 | AX-89883016 | 50.39 |
| | | 2011 | F4c | 1 | AX-89829510 | 2.48 | 4.09 | 0.02 | 13.39 | tggx147c_F4c | 0.00 | b008-b_4c | 18.53 |
| Kaempferol-glucuronide | KGn | 2011 | F3d | 2 | cF3d.loc21 | 21.00 | 2.66 | 0.01 | 6.20 | AX-89826704 | 8.17 | AX-89786753 | 159.46 |
| | | | M4c | 2 | cM4c.loc1 | 1.00 | 5.08 | -0.02 | 16.37 | AX-89791452 | 0.00 | gtat238c | 10.14 |
| | | | F4c | 2 | cF4c.loc18.5 | 18.50 | 5.56 | -0.01 | 15.39 | AX-89791467 | 4.31 | AX-89889008 | 22.08 |
| | | | M6d | 2 | AX-89805325 | 163.43 | 3.11 | -0.01 | 11.01 | AX-89796107 | 150.67 | tcaa540c | 186.97 |
| | KCoGs | 2011 | M5c | 1 | AX-89835889 | 19.97 | 3.08 | -0.01 | 10.26 | tgtt140c | 13.29 | AX-89794844 | 25.75 |

| | | | | | | | | | | | | | |
|---|---|---|---|---|---|---|---|---|---|---|---|---|---|
| Kaempferol-coumaryl-glucoside | | 2010 | M6b | 1 | AX-89896029 | 36.12 | 3.27 | -0.02 | 18.88 | cctc218d | 30.42 | tcac235c | 45.98 |
| Quercetin-glucuronide | QGn | 2011 | M6d | 1 | AX-89805325 | 163.43 | 2.54 | -0.01 | 8.53 | gaat480c | 0.00 | caaa167c | 194.93 |
| **Flavan-3-ols** | | | | | | | | | | | | | |
| Total Flavan-3-ols | F3ol | 2011 | M3c1 | 2 | AX-89785338 | 16.23 | 3.30 | 0.24 | 10.33 | i075224c | 11.32 | AX-89904746 | 19.29 |
| | | 2010 | M5a | 2 | cM5a.loc23.5 | 23.50 | 2.85 | -0.25 | 13.19 | AX-89793472 | 22.63 | gtgx171c | 33.17 |
| | | 2011 | M5a | 2 | AX-89835724 | 46.80 | 5.08 | -0.30 | 15.75 | AX-89828890 | 39.96 | AX-89794385 | 56.84 |
| | | 2010 | M6a | 2 | AX-89796438 | 123.15 | 4.22 | -0.28 | 16.99 | tgat340c | 116.22 | AX-89914769 | 128.09 |
| Catechin | Cat | 2011 | F6a | 1 | cF6a.loc240.5 | 240.50 | 2.61 | 0.03 | 8.76 | AX-89899984 | 37.75 | gaaa523c | 241.22 |
| (epi)Catechin dimers | CatCat | 2011 | M2c | 1 | AX-89782622 | 86.22 | 3.79 | -0.09 | 12.47 | AX-89865363 | 78.73 | i146188c | 88.67 |
| (epi)Afzelechin-(epi)catechin dimers | AfCat | 2011 | M1a | 4 | AX-89873448 | 123.16 | 5.51 | 0.02 | 18.42 | AX-89816417 | 118.16 | AX-89864948 | 125.60 |
| | | 2011 | F1b | 2 | AX-89798396 | 71.58 | 3.18 | 0.01 | 9.39 | AX-89798440 | 65.40 | AX-89846879 | 79.87 |
| | | 2011 | F2c | 2 | AX-89819622 | 38.88 | 2.05 | -0.01 | 5.69 | gctg285c | 0.00 | AX-89781524 | 56.43 |
| | | 2011 | M5a | 4 | cM5a.loc47 | 47.00 | 3.16 | -0.01 | 5.09 | AX-89828890 | 39.96 | AX-89794385 | 56.84 |
| | | 2011 | M5b | 4 | AX-89836519 | 38.05 | 5.52 | 0.02 | 18.18 | AX-89794083 | 31.26 | AX-89794556 | 39.88 |
| | | 2010 | F5d | 1 | cF5d.loc158.5 | 158.50 | 3.16 | -0.01 | 18.26 | AX-89849187 | 153.04 | AX-89904249 | 164.15 |
| | | 2011 | M6a | 4 | AX-89914769 | 128.09 | 5.38 | 0.02 | 15.84 | AX-89795266 | 120.71 | AX-89914804 | 129.92 |
| (epi)Afzelechin-glucoside | AfGs | 2011 | M3a | 3 | cM3a.loc30.5 | 30.50 | 4.03 | 0.19 | 9.70 | AX-89785262 | 20.73 | AX-89784824 | 33.27 |
| | | 2011 | M5a | 3 | cM5a.loc58 | 58.00 | 3.50 | -0.19 | 9.68 | AX-89905829 | 46.80 | b037227c | 64.95 |
| | | 2010 | M6a | 1 | cM6a.loc126 | 126.00 | 2.71 | -0.24 | 15.96 | catc138c | 49.50 | AX-89840892 | 205.99 |
| | | 2011 | M6a | 3 | cM6a.loc165 | 165.00 | 5.82 | -0.25 | 16.31 | tcaa168c | 153.98 | AX-89797755 | 168.49 |
| **Colorimetry** | | | | | | | | | | | | | |
| Anthocyanins (colorimetry) | ANTHc | 2011 | M1a | 2 | AX-89816735 | 36.09 | 3.09 | 130.41 | 9.91 | i072226c | 2.54 | AX-89875150 | 41.00 |
| | | 2010 | M4a | 1 | tgaa125c | 104.80 | 2.86 | 166.41 | 17.87 | gatc380c | 0.00 | gtta105c | 165.84 |
| | | 2011 | M6a | 2 | AX-89840892 | 205.99 | 5.19 | 167.22 | 16.31 | AX-89797823 | 200.76 | AX-89895308 | 220.62 |
| | | 2011 | F6a | 1 | cF6a.loc103.5 | 103.50 | 5.63 | -176.14 | 17.97 | gctg280c | 89.11 | AX-89795985 | 104.70 |
| Colour | COLOUR | 2011 | M1a | 2 | v017213c | 52.64 | 4.80 | 0.89 | 11.21 | AX-89875160 | 41.65 | AX-89780210 | 53.87 |
| | | | M3a | 2 | cM3a.loc9.5 | 9.50 | 4.08 | 0.85 | 10.10 | AX-89826900 | 8.86 | AX-89804053 | 18.77 |
| | | | F4d | 2 | AX-89790310 | 102.21 | 2.89 | 0.80 | 8.99 | gttc187c | 0.00 | i136147c | 110.44 |
| | | | F6a | 2 | cF6a.loc53.5 | 53.50 | 3.61 | -0.89 | 11.06 | AX-89899878 | 41.45 | AX-89899245 | 56.23 |

**Supplemental Table 6.** Values of the QTL thresholds at 5 and 10% used for CIM analysis. The thresholds were calculated on 1000 permutations for each trait for male and female in 2010 and 2011.

| Traits | Abbr. | Male threshold | | | | Female threshold | | | |
|---|---|---|---|---|---|---|---|---|---|
| | | 2010 | | 2011 | | 2010 | | 2011 | |
| | | 5% | 10% | 5% | 10% | 5% | 10% | 5% | 10% |
| **Anthocyanins** | | | | | | | | | |
| Total anthocyanins | Ant | 3.07 | 2.79 | 3.16 | 2.73 | 3.23 | 2.84 | 2.13 | 2.12 |
| Pelargonidin-3-glucoside | PgGs | 3.05 | 2.75 | 3.21 | 2.80 | 3.17 | 2.86 | 2.25 | 2.16 |
| Pelargonidin-3-glucoside-malonate | PgGsM | 2.04 | 1.87 | 1.92 | 1.78 | 2.13 | 1.97 | 1.78 | 1.75 |
| Pelargonidin-3-rutinoside | PgRs | 2.79 | 2.55 | 3.17 | 2.76 | 3.07 | 2.81 | 2.85 | 2.72 |
| Cyanidin-3-glucoside | CyGs | 2.80 | 2.54 | 3.13 | 2.89 | 2.89 | 2.64 | 2.61 | 2.60 |
| (epi)Afzelechin-pelargonidin-glucoside | AfPgGs | 3.23 | 2.89 | 3.09 | 2.78 | 3.29 | 2.94 | 2.51 | 2.45 |
| **Flavonols** | | | | | | | | | |

| | | | | | | | | | |
|---|---|---|---|---|---|---|---|---|---|
| Total Flavonols | Fvo | 2.93 | 2.64 | 3.08 | 2.77 | 2.95 | 2.73 | 2.02 | 2.01 |
| Kaempferol-glucoside | KGs | 3.11 | 2.73 | 3.01 | 2.75 | 3.17 | 2.84 | 2.04 | 2.01 |
| Kaempferol-glucuronide | KGn | 3.03 | 2.77 | 3.20 | 2.85 | 3.05 | 2.78 | 2.16 | 2.11 |
| Kaempferol-coumaryl-glucoside | KCoGs | 3.19 | 2.71 | 3.22 | 2.89 | 3.21 | 2.88 | 3.21 | 3.08 |
| Quercetin-glucuronide | QGn | 2.28 | 2.10 | 3.04 | 2.78 | 2.29 | 2.12 | 2.43 | 2.40 |
| **Flavan-3-ols** | | | | | | | | | |
| Total Flavan-3-ols | F3ol | 2.77 | 2.57 | 3.04 | 2.78 | 3.00 | 2.71 | 3.45 | 3.23 |
| Catechin | Cat | 3.02 | 2.65 | 3.02 | 2.80 | 3.25 | 2.80 | 2.68 | 2.63 |
| (epi)Catechin dimers | CatCat | 3.15 | 2.84 | 3.05 | 2.76 | 3.23 | 2.91 | 1.96 | 1.95 |
| (epi)Afzelechin-(epi)catechin dimers | AfCat | 2.97 | 2.67 | 3.12 | 2.79 | 3.22 | 2.89 | 1.89 | 1.87 |
| (epi)Afzelechin-glucoside | AfGs | 2.82 | 2.54 | 3.07 | 2.75 | 3.09 | 2.76 | 2.16 | 2.15 |
| **Colorimetry** | | | | | | | | | |
| Anthocyanins (colorimetry) | ANTHc | 3.11 | 2.77 | 3.15 | 2.80 | 3.19 | 2.87 | 5.15 | 4.77 |
| **Visual assessment** | | | | | | | | | |
| Colour | COLOUR | NA | NA | 3.07 | 2.79 | NA | NA | 2.42 | 2.41 |

**Supplemental Table 7.** List of genes from the FvH4_v4.0.a2 version of the Fragaria vesca reference genome located in the male M3a colour-related QTLs (PgGs etc): physical position from 1.213489 Mb to 2.673762 Mb (extract from Li et al., 2019, Sup Table S2)

| Gene ID (v2.0.a2) | Gene location | LG | Start | End | Gene ID (v4.0.a1) | Gene ID (v2.0.a2) | AED score | GO term | symbol | Description |
|---|---|---|---|---|---|---|---|---|---|---|
| gene36090 | Fvb3:1214342-1217361 | Fvb3 | 1214342 | 1217361 | FvH4_3g02420 | gene36090 | 0.11 | GO:0000741 | | |
| gene19584 | Fvb3:1217959-1218596 | Fvb3 | 1217959 | 1218596 | FvH4_3g02421 | gene19584 | 1 | | | |
| gene19583 | Fvb3:1218676-1220955 | Fvb3 | 1218676 | 1220955 | FvH4_3g02430 | gene19583 | 0.11 | GO:0008654,GO:0016020,GO:0016780 | CLS | cardiolipin synthase |
| gene19582 | Fvb3:1221719-1224257 | Fvb3 | 1221719 | 1224257 | FvH4_3g02440 | gene19582 | 0.2 | GO:0004672,GO:0005524,GO:0006468 | Protein kinase superfamily protein | |
| gene19581 | Fvb3:1223868-1224263 | Fvb3 | 1223868 | 1224263 | FvH4_3g02450 | gene19581 | 1 | | | |
| gene19580 | Fvb3:1225360-1232457 | Fvb3 | 1225360 | 1232457 | FvH4_3g02460 | gene19580 | 0.13 | GO:0005515,GO:0006629 | phospholipases;galactolipases | |
| gene19579 | Fvb3:1233807-1234424 | Fvb3 | 1233807 | 1234424 | FvH4_3g02470 | gene19579 | 1 | | | LA RNA-binding protein |
| gene19578 | Fvb3:1235098-1238130 | Fvb3 | 1235098 | 1238130 | FvH4_3g02490 | gene19578 | 0.26 | GO:0019239 | | adenosine/AMP deaminase family protein |
| gene19577 | Fvb3:1237674-1242642 | Fvb3 | 1237674 | 1242642 | FvH4_3g02480 | gene19577 | 0.09 | GO:0001522,GO:0003723,GO:0009982 | Pseudouridine synthase family protein | |
| gene36091 | Fvb3:1241491-1247952 | Fvb3 | 1241491 | 1247952 | FvH4_3g02481 | gene36091 | 0.11 | GO:0004553,GO:0005975 | BGLU47 | beta-glucosidase 47 |
| gene36092 | Fvb3:1248042-1251710 | Fvb3 | 1248042 | 1251710 | FvH4_3g02500 | gene36092 | 0.22 | GO:0016021,GO:0022857,GO:0055085 | ATGPT1, GPT1 | glucose 6-phosphate/phosphate translocator 1 |
| gene19573 | Fvb3:1255971-1262076 | Fvb3 | 1255971 | 1262076 | FvH4_3g02510 | gene19573 | 0.08 | GO:0003676 | PCFS4 | PCF11P-similar protein 4 |
| gene19572 | Fvb3:1267960-1273443 | Fvb3 | 1267960 | 1273443 | FvH4_3g02520 | gene19572 | 0.23 | GO:0006355,GO:0008289,GO:0043565 | PDF2 | protodermal factor 2 |
| gene19570 | Fvb3:1279701-1282170 | Fvb3 | 1279701 | 1282170 | FvH4_3g02530 | gene19570 | 0 | GO:0010215,GO:0016049,GO:0031225 | COBL11 | COBRA-like protein 11 precursor |
| gene19568 | Fvb3:1282882-1283827 | Fvb3 | 1282882 | 1283827 | FvH4_3g02550 | gene19568 | 0.25 | | NTL | NTF2-like |
| gene19567 | Fvb3:1284357-1286597 | Fvb3 | 1284357 | 1286597 | FvH4_3g02560 | gene19567 | 0.1 | GO:0004857,GO:0005618,GO:0030599,GO:0042545 | ATPMEPCRA,PMEPCRA | methylesterase PCR A |
| gene19566 | Fvb3:1289201-1292633 | Fvb3 | 1289201 | 1292633 | FvH4_3g02570 | gene19566 | 0.58 | | RIC7 | PAK-box/P21-Rho-binding family protein |
| gene19564 | Fvb3:1293444-1294926 | Fvb3 | 1293444 | 1294926 | FvH4_3g02571 | gene19564 | 1 | | | |
| gene19563 | Fvb3:1298132-1298446 | Fvb3 | 1298132 | 1298446 | FvH4_3g02572 | gene19563 | 1 | | ATFC-II,FC-II,FC2 | ferrochelatase 2 |
| gene19562 | Fvb3:1301418-1303514 | Fvb3 | 1301418 | 1303514 | FvH4_3g02580 | gene19562 | 1 | GO:0005515 | | F-box/RNI-like superfamily protein |
| gene19561 | Fvb3:1304903-1305370 | Fvb3 | 1304903 | 1305370 | FvH4_3g02581 | gene19561 | 1 | | ACS12 | 1-amino-cyclopropane-1-carboxylate synthase 12 |
| gene39837 | Fvb3:1306058-1307635 | Fvb3 | 1306058 | 1307635 | FvH4_3g02582 | gene39837 | 0.1 | GO:0003676 | | |
| gene19560 | Fvb3:1307821-1309174 | Fvb3 | 1307821 | 1309174 | FvH4_3g02590 | gene19560 | 0.44 | | | |
| gene19559 | Fvb3:1312993-1315394 | Fvb3 | 1312993 | 1315394 | FvH4_3g02591 | gene19559 | 0.15 | GO:0005515 | | F-box/RNI-like superfamily protein |
| gene39838 | Fvb3:1316535-1318003 | Fvb3 | 1316535 | 1318003 | FvH4_3g02592 | gene39838 | 1 | | | |
| gene19558 | Fvb3:1318229-1319213 | Fvb3 | 1318229 | 1319213 | FvH4_3g02593 | gene19558 | 0.43 | | | |
| gene19557 | Fvb3:1322663-1323853 | Fvb3 | 1322663 | 1323853 | FvH4_3g02594 | gene19557 | 0.14 | GO:0005515 | | F-box family protein |

| | | | | | | | | | | |
|---|---|---|---|---|---|---|---|---|---|---|
| gene19556 | Fvb3:1324979-1325998 | Fvb3 | 1324979 | 1325998 | FvH4_3g02600 | gene19556 | 0.33 | | | Oligosaccharyltransferase complex/magnesium transporter family protein |
| gene19555 | Fvb3:1326638-1327957 | Fvb3 | 1326638 | 1327957 | FvH4_3g02620 | gene19555 | 0.27 | GO:0005515 | | Tetratricopeptide repeat (TPR)-like superfamily protein |
| gene19553 | Fvb3:1331732-1333854 | Fvb3 | 1331732 | 1333854 | FvH4_3g02630 | gene19553 | 0.08 | GO:0004553,GO:0005618,GO:0010411,GO:0016762,GO:0042546,GO:0048046 | XTH8 | xyloglucan endotransglucosylase/hydrolase 8 |
| gene19552 | Fvb3:1334771-1336329 | Fvb3 | 1334771 | 1336329 | FvH4_3g02640 | gene19552 | 0 | | | postsynaptic protein-related |
| gene19551 | Fvb3:1337019-1340760 | Fvb3 | 1337019 | 1340760 | FvH4_3g02641 | gene19551 | 1 | GO:0005515 | UBQ10 | polyubiquitin 10 |
| gene19550 | Fvb3:1340958-1345868 | Fvb3 | 1340958 | 1345868 | FvH4_3g02650 | gene19550 | 0.11 | GO:0005515,GO:0005634,GO:0006281,GO:0008081 | TDP1 | tyrosyl-DNA phosphodiesterase-related |
| gene36093 | Fvb3:1347333-1351040 | Fvb3 | 1347333 | 1351040 | FvH4_3g02660 | gene36093 | 0.19 | | | Chaperone DnaJ-domain superfamily protein |
| gene36094 | Fvb3:1351967-1355057 | Fvb3 | 1351967 | 1355057 | FvH4_3g02670 | gene36094 | 0.02 | GO:0016491,GO:0055114 | ATGA2OX8,GA2OX8 | gibberellin 2-oxidase 8 |
| gene19547 | Fvb3:1360388-1360757 | Fvb3 | 1360388 | 1360757 | FvH4_3g02671 | gene19547 | 1 | | | |
| gene19546 | Fvb3:1365621-1367459 | Fvb3 | 1365621 | 1367459 | FvH4_3g02680 | gene19546 | 0.26 | GO:0005524,GO:0016772 | ATRP1,RP1 | PPDK regulatory protein |
| gene19545 | Fvb3:1368552-1371552 | Fvb3 | 1368552 | 1371552 | FvH4_3g02690 | gene19545 | 0.16 | | | |
| gene19544 | Fvb3:1378211-1380130 | Fvb3 | 1378211 | 1380130 | FvH4_3g02700 | gene19544 | 0.18 | GO:0004601,GO:0006979,GO:0020037,GO:0042744,GO:0055114 | Peroxidase superfamily protein | |
| gene39839 | Fvb3:1380840-1381773 | Fvb3 | 1380840 | 1381773 | FvH4_3g02710 | gene39839 | 1 | | | Late embryogenesis abundant (LEA) hydroxyproline-rich glycoprotein family |
| gene19543 | Fvb3:1383480-1384064 | Fvb3 | 1383480 | 1384064 | FvH4_3g02720 | gene19543 | 1 | | NDR1 | Late embryogenesis abundant (LEA) hydroxyproline-rich glycoprotein family |
| gene19542 | Fvb3:1384043-1386849 | Fvb3 | 1384043 | 1386849 | FvH4_3g02730 | gene19542 | 0.35 | GO:0009245,GO:0016410 | Trimeric LpxA-like enzymes superfamily protein | |
| gene19541 | Fvb3:1386762-1390102 | Fvb3 | 1386762 | 1390102 | FvH4_3g02731 | gene19541 | 0.47 | GO:0004672,GO:0005524,GO:0006468 | CRK25 | cysteine-rich RLK (RECEPTOR-like protein kinase) 25 |
| gene36095 | Fvb3:1390832-1393829 | Fvb3 | 1390832 | 1393829 | FvH4_3g02732 | gene36095 | 0.47 | GO:0004672,GO:0005524,GO:0006468 | CRK25 | cysteine-rich RLK (RECEPTOR-like protein kinase) 25 |
| gene39840 | Fvb3:1393775-1394278 | Fvb3 | 1393775 | 1394278 | FvH4_3g02750 | gene39840 | 0.23 | | | Trimeric LpxA-like enzymes superfamily protein |

| | | | | | | | | | | |
|---|---|---|---|---|---|---|---|---|---|---|
| gene36096 | Fvb3:1394904-1397882 | Fvb3 | 1394904 | 1397882 | FvH4_3g02751 | gene36096 | 0.47 | GO:0004672,GO:0005524,GO:0006468 | CRK25 | cysteine-rich RLK (RECEPTOR-like protein kinase) 25 |
| gene19539 | Fvb3:1398823-1401464 | Fvb3 | 1398823 | 1401464 | FvH4_3g02752 | gene19539 | 0.48 | GO:0004672,GO:0005524,GO:0006468 | CRK25 | cysteine-rich RLK (RECEPTOR-like protein kinase) 25 |
| gene19538 | Fvb3:1406724-1407994 | Fvb3 | 1406724 | 1407994 | FvH4_3g02753 | gene19538 | 1 | | | |
| gene19537 | Fvb3:1418298-1425301 | Fvb3 | 1418298 | 1425301 | FvH4_3g02754 | gene19537 | 0.47 | GO:0004672,GO:0005524,GO:0006468 | CRK25 | cysteine-rich RLK (RECEPTOR-like protein kinase) 25 |
| gene19534 | Fvb3:1426375-1430307 | Fvb3 | 1426375 | 1430307 | FvH4_3g02770 | gene19534 | 0.47 | GO:0004672,GO:0005524,GO:0006468 | CRK25 | cysteine-rich RLK (RECEPTOR-like protein kinase) 25 |
| gene19533 | Fvb3:1431110-1435446 | Fvb3 | 1431110 | 1435446 | FvH4_3g02771 | gene19533 | 0.45 | GO:0004672,GO:0005524,GO:0006468 | CRK10,RLK4 | cysteine-rich RLK (RECEPTOR-like protein kinase) 10 |
| gene19532 | Fvb3:1437798-1443554 | Fvb3 | 1437798 | 1443554 | FvH4_3g02772 | gene19532 | 0.46 | GO:0004672,GO:0005524,GO:0006468 | CRK10,RLK4 | cysteine-rich RLK (RECEPTOR-like protein kinase) 10 |
| gene19530 | Fvb3:1449956-1453571 | Fvb3 | 1449956 | 1453571 | FvH4_3g02773 | gene19530 | 0.51 | GO:0004672,GO:0005524,GO:0006468 | CRK25 | cysteine-rich RLK (RECEPTOR-like protein kinase) 25 |
| gene19528 | Fvb3:1456071-1458661 | Fvb3 | 1456071 | 1458661 | FvH4_3g02774 | gene19528 | 0.47 | GO:0004672,GO:0005524,GO:0006468 | CRK25 | cysteine-rich RLK (RECEPTOR-like protein kinase) 25 |
| gene19527 | Fvb3:1465643-1466101 | Fvb3 | 1465643 | 1466101 | FvH4_3g02790 | gene19527 | 1 | GO:0006855,GO:0015238,GO:0015297,GO:0016020 | MATE efflux family protein | |
| gene39841 | Fvb3:1471384-1473237 | Fvb3 | 1471384 | 1473237 | FvH4_3g02800 | gene39841 | 0.44 | GO:0004672,GO:0005524,GO:0006468 | CRK25 | cysteine-rich RLK (RECEPTOR-like protein kinase) 25 |
| gene19526 | Fvb3:1473385-1473941 | Fvb3 | 1473385 | 1473941 | FvH4_3g02801 | gene19526 | 0.48 | GO:0004672,GO:0006468 | CRK33 | cysteine-rich RLK (RECEPTOR-like protein kinase) 33 |
| gene19525 | Fvb3:1475940-1479008 | Fvb3 | 1475940 | 1479008 | FvH4_3g02820 | gene19525 | 0.41 | GO:0004672,GO:0005524,GO:0006468 | CRK25 | cysteine-rich RLK (RECEPTOR-like protein kinase) 25 |
| gene19524 | Fvb3:1479970-1482127 | Fvb3 | 1479970 | 1482127 | FvH4_3g02830 | gene19524 | 0.57 | GO:0004672,GO:0005524,GO:0006468 | CRK25 | cysteine-rich RLK (RECEPTOR-like protein kinase) 25 |
| gene19523 | Fvb3:1482849-1484843 | Fvb3 | 1482849 | 1484843 | FvH4_3g02831 | gene19523 | 1 | GO:0006855,GO:0015238,GO:0015297,GO:0016020 | MATE efflux family protein | |
| gene36098 | Fvb3:1488257-1490637 | Fvb3 | 1488257 | 1490637 | FvH4_3g02832 | gene36098 | 0.45 | | CRK25 | cysteine-rich RLK (RECEPTOR-like protein kinase) 25 |
| gene36099 | Fvb3:1491475-1492987 | Fvb3 | 1491475 | 1492987 | FvH4_3g02833 | gene36099 | 0.53 | | CRK29 | cysteine-rich RLK (RECEPTOR-like protein kinase) 29 |
| gene36100 | Fvb3:1494127-1497293 | Fvb3 | 1494127 | 1497293 | FvH4_3g02834 | gene36100 | 0.42 | GO:0004672,GO:0005524,GO:0006468 | CRK29 | cysteine-rich RLK (RECEPTOR-like protein kinase) 29 |
| gene36103 | Fvb3:1498681-1511405 | Fvb3 | 1498681 | 1511405 | FvH4_3g02850 | gene36103 | 0.39 | | ATRPA70C,RPA70C | Replication factor-A protein 1-related |

| | | | | | | | | | | |
|---|---|---|---|---|---|---|---|---|---|---|
| gene19521 | Fvb3:1512633-1513880 | Fvb3 | 1512633 | 1513880 | FvH4_3g02851 | gene19521 | 1 | GO:0003676 | | |
| gene24648 | Fvb3:1528115-1528692 | Fvb3 | 1528115 | 1528692 | FvH4_3g02852 | gene24648 | 1 | | | |
| gene24649 | Fvb3:1531980-1534897 | Fvb3 | 1531980 | 1534897 | FvH4_3g02870 | gene24649 | 0.12 | GO:0004672,GO:0005524,GO:0006468 | CRK29 | cysteine-rich RLK (RECEPTOR-like protein kinase) 29 |
| gene36104 | Fvb3:1535226-1536646 | Fvb3 | 1535226 | 1536646 | FvH4_3g02871 | gene36104 | 0 | | | |
| gene24650 | Fvb3:1536971-1537610 | Fvb3 | 1536971 | 1537610 | FvH4_3g02880 | gene24650 | 1 | GO:0003677,GO:0046983 | AGL29 | AGAMOUS-like 29 |
| gene24651 | Fvb3:1537709-1539815 | Fvb3 | 1537709 | 1539815 | FvH4_3g02881 | gene24651 | 1 | | | |
| gene39842 | Fvb3:1540876-1541927 | Fvb3 | 1540876 | 1541927 | FvH4_3g02882 | gene39842 | 1 | | ATK5 | kinesin 5 |
| gene24652 | Fvb3:1542978-1545247 | Fvb3 | 1542978 | 1545247 | FvH4_3g02890 | gene24652 | 0.01 | | | RNI-like superfamily protein |
| gene24653 | Fvb3:1546731-1549678 | Fvb3 | 1546731 | 1549678 | FvH4_3g02900 | gene24653 | 0.04 | GO:0016021 | | Sulfite exporter TauE/SafE family protein |
| gene24654 | Fvb3:1549701-1552477 | Fvb3 | 1549701 | 1552477 | FvH4_3g02910 | gene24654 | 0.16 | GO:0046983 | bHLH105,ILR3 | basic helix-loop-helix (bHLH) DNA-binding superfamily protein |
| gene24655 | Fvb3:1553281-1554668 | Fvb3 | 1553281 | 1554668 | FvH4_3g02911 | gene24655 | 0.04 | GO:0006662,GO:0015035,GO:0045454 | ATCXXS1,CXXS1 | C-terminal cysteine residue is changed to a serine 1 |
| gene24656 | Fvb3:1554833-1559824 | Fvb3 | 1554833 | 1559824 | FvH4_3g02912 | gene24656 | 0.09 | GO:0003777,GO:0005524,GO:0007018,GO:0008017 | ATK1,KATA,KATAP | kinesin 1 |
| gene24657 | Fvb3:1561438-1562904 | Fvb3 | 1561438 | 1562904 | FvH4_3g02920 | gene24657 | 0.17 | GO:0005509,GO:0009654,GO:0015979,GO:0019898 | PSBQ,PSBQ-2,PSII-Q | photosystem II subunit Q-2 |
| gene24658 | Fvb3:1564146-1567056 | Fvb3 | 1564146 | 1567056 | FvH4_3g02930 | gene24658 | 1 | GO:0006355 | | DNA-binding storekeeper protein-related transcriptional regulator |
| gene24659 | Fvb3:1569121-1570130 | Fvb3 | 1569121 | 1570130 | FvH4_3g02940 | gene24659 | 0.15 | GO:0000786,GO:0003677,GO:0005634,GO:0046982 | HTA2 | histone H2A 2 |
| gene24660 | Fvb3:1569878-1570816 | Fvb3 | 1569878 | 1570816 | FvH4_3g02950 | gene24660 | 0.03 | | | Protein of unknown function (DUF1218) |
| gene24661 | Fvb3:1571146-1573912 | Fvb3 | 1571146 | 1573912 | FvH4_3g02960 | gene24661 | 0.19 | GO:0003824 | HSA32 | Aldolase-type TIM barrel family protein |
| gene24663 | Fvb3:1576006-1578248 | Fvb3 | 1576006 | 1578248 | FvH4_3g02970 | gene24663 | 0.36 | | | |
| gene24664 | Fvb3:1578846-1582265 | Fvb3 | 1578846 | 1582265 | FvH4_3g02971 | gene24664 | 1 | | | |
| gene24665 | Fvb3:1583755-1586819 | Fvb3 | 1583755 | 1586819 | FvH4_3g02980 | gene24665 | 0.21 | GO:0003824,GO:0050662 | BAN | NAD(P)-binding Rossmann-fold superfamily protein |
| gene24666 | Fvb3:1585474-1588821 | Fvb3 | 1585474 | 1588821 | FvH4_3g02990 | gene24666 | 0.06 | GO:0004672,GO:0005524,GO:0006468 | CRLK1 | Protein kinase superfamily protein |
| gene24667 | Fvb3:1590747-1593542 | Fvb3 | 1590747 | 1593542 | FvH4_3g03000 | gene24667 | 0.12 | GO:0000287,GO:0008152,GO:0010333 | ATTPS03,TPS03 | terpene synthase 03 |
| gene24668 | Fvb3:1594603-1595709 | Fvb3 | 1594603 | 1595709 | FvH4_3g03010 | gene24668 | 0.27 | GO:0004842,GO:0016567 | RING/U-box superfa | |

| | | | | | | | | | | |
|---|---|---|---|---|---|---|---|---|---|---|
| | | | | | | | | | mily protein | |
| gene24669 | Fvb3:1599666-1604244 | Fvb3 | 1599666 | 1604244 | FvH4_3g03030 | gene24669 | 0.18 | GO:0003743 | | eukaryotic translation initiation factor-related |
| gene24670 | Fvb3:1604517-1607349 | Fvb3 | 1604517 | 1607349 | FvH4_3g03040 | gene24670 | 0 | GO:0005515 | | Phototropic-responsive NPH3 family protein |
| gene24671 | Fvb3:1607881-1612771 | Fvb3 | 1607881 | 1612771 | FvH4_3g03041 | gene24671 | 0.04 | GO:0005515,GO:0046872 | phosphoinositide binding | |
| gene24672 | Fvb3:1613290-1615710 | Fvb3 | 1613290 | 1615710 | FvH4_3g03042 | gene24672 | 0.43 | GO:0000287,GO:0008152,GO:0010333 | ATTPS14,TPS14 | terpene synthase 14 |
| gene34030 | Fvb3:1615982-1617349 | Fvb3 | 1615982 | 1617349 | FvH4_3g03043 | gene34030 | 1 | | | Tetratricopeptide repeat (TPR)-like superfamily protein |
| gene36105 | Fvb3:1623630-1624412 | Fvb3 | 1623630 | 1624412 | FvH4_3g03044 | gene36105 | 0.66 | | | |
| gene39843 | Fvb3:1624708-1629243 | Fvb3 | 1624708 | 1629243 | FvH4_3g03045 | gene39843 | 0.44 | GO:0000287,GO:0008152,GO:0010333 | ATTPS14,TPS14 | terpene synthase 14 |
| gene24675 | Fvb3:1629270-1629924 | Fvb3 | 1629270 | 1629924 | FvH4_3g03060 | gene24675 | 0.46 | GO:0005515 | | Tetratricopeptide repeat (TPR)-like superfamily protein |
| gene24676 | Fvb3:1633206-1635362 | Fvb3 | 1633206 | 1635362 | FvH4_3g03061 | gene24676 | 0.44 | GO:0000287,GO:0008152,GO:0010333 | ATTPS14,TPS14 | terpene synthase 14 |
| gene24677 | Fvb3:1635670-1636316 | Fvb3 | 1635670 | 1636316 | FvH4_3g03070 | gene24677 | 0.41 | GO:0005515 | | Tetratricopeptide repeat (TPR)-like superfamily protein |
| gene24678 | Fvb3:1636706-1637892 | Fvb3 | 1636706 | 1637892 | FvH4_3g03071 | gene24678 | 0.66 | | | phosphoinositide binding |
| gene39844 | Fvb3:1642571-1645757 | Fvb3 | 1642571 | 1645757 | FvH4_3g03072 | gene39844 | 0.6 | GO:0000287,GO:0008152,GO:0010333 | ATTPS14,TPS14 | terpene synthase 14 |
| gene24673 | Fvb3:1645119-1647527 | Fvb3 | 1645119 | 1647527 | FvH4_3g03080 | gene24673 | 0.37 | GO:0005515 | | Tetratricopeptide repeat (TPR)-like superfamily protein |
| gene39845 | Fvb3:1649467-1654658 | Fvb3 | 1649467 | 1654658 | FvH4_3g03081 | gene39845 | 0.42 | GO:0000287,GO:0008152,GO:0010333 | ATTPS14,TPS14 | terpene synthase 14 |
| gene39846 | Fvb3:1654662-1655297 | Fvb3 | 1654662 | 1655297 | FvH4_3g03090 | gene39846 | 0.45 | GO:0005515 | | Tetratricopeptide repeat (TPR)-like superfamily protein |
| gene39847 | Fvb3:1657959-1658828 | Fvb3 | 1657959 | 1658828 | FvH4_3g03100 | gene39847 | 1 | GO:0009055,GO:0015035,GO:0045454 | Glutaredoxin family protein | |
| gene36106 | Fvb3:1664540-1666004 | Fvb3 | 1664540 | 1666004 | FvH4_3g03101 | gene36106 | 1 | GO:0003677,GO:0046983 | BED zinc finger ;hAT family dimerisation domain | |
| gene39848 | Fvb3:1673389-1676118 | Fvb3 | 1673389 | 1676118 | FvH4_3g03110 | gene39848 | 0.44 | | | |
| gene36107 | Fvb3:1677359-1678352 | Fvb3 | 1677359 | 1678352 | FvH4_3g03120 | gene36107 | 1 | | | |
| gene30670 | Fvb3:1678430-1680235 | Fvb3 | 1678430 | 1680235 | FvH4_3g03130 | gene30670 | 0 | GO:0008168 | ATBSMT1,BSMT1 | S-adenosyl-L-methionine-dependent methyltransferases superfamily protein |

| | | | | | | | | | | |
|---|---|---|---|---|---|---|---|---|---|---|
| gene30669 | Fvb3:1680583-1685225 | Fvb3 | 1680583 | 1685225 | FvH4_3g03150 | gene30669 | 0.2 | GO:0000287,GO:0008152,GO:0010333 | ATTPS14,TPS14 | terpene synthase 14 |
| gene30668 | Fvb3:1685456-1686592 | Fvb3 | 1685456 | 1686592 | FvH4_3g03160 | gene30668 | 0 | GO:0005515 | | Tetratricopeptide repeat (TPR)-like superfamily protein |
| gene30667 | Fvb3:1687520-1688818 | Fvb3 | 1687520 | 1688818 | FvH4_3g03170 | gene30667 | 0.37 | GO:0046983 | DYT1 | basic helix-loop-helix (bHLH) DNA-binding superfamily protein |
| gene36108 | Fvb3:1690369-1690947 | Fvb3 | 1690369 | 1690947 | FvH4_3g03180 | gene36108 | 0.32 | | | Protein of unknown function, DUF538 |
| gene36109 | Fvb3:1691861-1692829 | Fvb3 | 1691861 | 1692829 | FvH4_3g03190 | gene36109 | 1 | GO:0070300 | | |
| gene30665 | Fvb3:1701110-1704158 | Fvb3 | 1701110 | 1704158 | FvH4_3g03200 | gene30665 | 0.09 | GO:0046983 | | basic helix-loop-helix (bHLH) DNA-binding superfamily protein |
| gene30664 | Fvb3:1707024-1710101 | Fvb3 | 1707024 | 1710101 | FvH4_3g03210 | gene30664 | 0.21 | GO:0003955,GO:0010181 | FQR1 | flavodoxin-like quinone reductase 1 |
| gene30663 | Fvb3:1712123-1715078 | Fvb3 | 1712123 | 1715078 | FvH4_3g03220 | gene30663 | 0.08 | GO:0004842,GO:0005515,GO:0016567 | B80,PUB8 | plant U-box 8 |
| gene30662 | Fvb3:1717851-1721952 | Fvb3 | 1717851 | 1721952 | FvH4_3g03230 | gene30662 | 0.43 | GO:0004674,GO:0005524,GO:0006468,GO:0048544 | S-locus lectin protein kinase family protein | |
| gene30661 | Fvb3:1729201-1732990 | Fvb3 | 1729201 | 1732990 | FvH4_3g03231 | gene30661 | 0.42 | GO:0004674,GO:0005524,GO:0006468,GO:0048544 | S-locus lectin protein kinase family protein | |
| gene30660 | Fvb3:1734251-1737670 | Fvb3 | 1734251 | 1737670 | FvH4_3g03240 | gene30660 | 0.44 | GO:0004674,GO:0005524,GO:0006468,GO:0048544 | S-locus lectin protein kinase family protein | |
| gene30659 | Fvb3:1740132-1743982 | Fvb3 | 1740132 | 1743982 | FvH4_3g03241 | gene30659 | 0.44 | GO:0004674,GO:0005524,GO:0006468,GO:0048544 | S-locus lectin protein kinase family protein | |
| gene30658 | Fvb3:1746423-1750099 | Fvb3 | 1746423 | 1750099 | FvH4_3g03242 | gene30658 | 0.44 | GO:0004674,GO:0005524,GO:0006468,GO:0048544 | S-locus lectin protein kinase family protein | |
| gene30657 | Fvb3:1754164-1757586 | Fvb3 | 1754164 | 1757586 | FvH4_3g03243 | gene30657 | 0.48 | GO:0004674,GO:0005524,GO:0006468,GO:0048544 | S-locus lectin protein kinase family protein | |
| gene30656 | Fvb3:1757967-1760459 | Fvb3 | 1757967 | 1760459 | FvH4_3g03250 | gene30656 | 1 | GO:0005515,GO:0006396 | crooked neck protein, putative / cell cycle protein, putative | |

| | | | | | | | | | | |
|---|---|---|---|---|---|---|---|---|---|---|
| gene30655 | Fvb3:1762502-1763232 | Fvb3 | 1762502 | 1763232 | FvH4_3g03251 | gene30655 | 0.35 | | | |
| gene30654 | Fvb3:1764174-1766346 | Fvb3 | 1764174 | 1766346 | FvH4_3g03260 | gene30654 | 0.14 | GO:0009690,GO:0019139,GO:0055114,GO:0071949 | ATCKX6, ATCKX7, CKX6 | cytokinin oxidase/dehydrogenase 6 |
| gene30653 | Fvb3:1768450-1769979 | Fvb3 | 1768450 | 1769979 | FvH4_3g03280 | gene30653 | 0.13 | GO:0005524,GO:0016887 | ATRLI2, RLI2 | RNAse l inhibitor protein 2 |
| gene30652 | Fvb3:1771083-1774428 | Fvb3 | 1771083 | 1774428 | FvH4_3g03300 | gene30652 | 0.38 | GO:0004674,GO:0005524,GO:0006468,GO:0048544 | S-locus lectin protein kinase family protein | |
| gene30651 | Fvb3:1775603-1779237 | Fvb3 | 1775603 | 1779237 | FvH4_3g03301 | gene30651 | 0.36 | GO:0004674,GO:0005524,GO:0006468,GO:0048544 | S-locus lectin protein kinase family protein | |
| gene30650 | Fvb3:1779994-1783527 | Fvb3 | 1779994 | 1783527 | FvH4_3g03310 | gene30650 | 0.37 | GO:0004674,GO:0005524,GO:0006468,GO:0048544 | S-locus lectin protein kinase family protein | |
| gene30649 | Fvb3:1786992-1790349 | Fvb3 | 1786992 | 1790349 | FvH4_3g03320 | gene30649 | 0.09 | GO:0004674,GO:0005524,GO:0006468,GO:0048544 | S-locus lectin protein kinase family protein | |
| gene39849 | Fvb3:1796903-1797935 | Fvb3 | 1796903 | 1797935 | FvH4_3g03321 | gene39849 | 1 | | | |
| gene39850 | Fvb3:1802085-1802574 | Fvb3 | 1802085 | 1802574 | FvH4_3g03322 | gene39850 | 0.47 | | B120 | S-locus lectin protein kinase family protein |
| gene39851 | Fvb3:1802838-1805248 | Fvb3 | 1802838 | 1805248 | FvH4_3g03323 | gene39851 | 0.35 | GO:0004672,GO:0005524,GO:0006468,GO:0048544 | S-locus lectin protein kinase family protein | |
| gene30646 | Fvb3:1805959-1806562 | Fvb3 | 1805959 | 1806562 | FvH4_3g03324 | gene30646 | 0.45 | | | |
| gene30645 | Fvb3:1812569-1815462 | Fvb3 | 1812569 | 1815462 | FvH4_3g03325 | gene30645 | 1 | | | |
| gene34404 | Fvb3:1815809-1816872 | Fvb3 | 1815809 | 1816872 | FvH4_3g03340 | gene34404 | 0.58 | GO:0048544 | | S-locus lectin protein kinase family protein |
| gene30644 | Fvb3:1818089-1821332 | Fvb3 | 1818089 | 1821332 | FvH4_3g03350 | gene30644 | 0.41 | GO:0004674,GO:0005524,GO:0006468,GO:0048544 | S-locus lectin protein kinase family protein | |
| gene39852 | Fvb3:1821652-1821882 | Fvb3 | 1821652 | 1821882 | FvH4_3g03351 | gene39852 | 0.49 | | | S-locus lectin protein kinase family protein |
| gene39853 | Fvb3:1821888-1822607 | Fvb3 | 1821888 | 1822607 | FvH4_3g03360 | gene39853 | 0.47 | | | |
| gene30643 | Fvb3:1829057-1832851 | Fvb3 | 1829057 | 1832851 | FvH4_3g03370 | gene30643 | 0.05 | GO:0004674,GO:0005524,GO:0006468,GO:0048544 | S-locus lectin protein kinase family protein | |
| gene30642 | Fvb3:1832450-1838776 | Fvb3 | 1832450 | 1838776 | FvH4_3g03390 | gene30642 | 0.35 | GO:0004674,GO:0005524,GO:0006468,GO:0048544 | S-locus lectin protein kinase family protein | |

| | | | | | | | | | | |
|---|---|---|---|---|---|---|---|---|---|---|
| gene30641 | Fvb3:1838900-1842240 | Fvb3 | 1838900 | 1842240 | FvH4_3g03400 | gene30641 | 0.08 | | | lysine decarboxylase family protein |
| gene30640 | Fvb3:1842526-1847095 | Fvb3 | 1842526 | 1847095 | FvH4_3g03410 | gene30640 | 0.37 | GO:0004674,GO:0005524,GO:0006468,GO:0048544 | S-locus lectin protein kinase family protein | |
| gene30639 | Fvb3:1846715-1850040 | Fvb3 | 1846715 | 1850040 | FvH4_3g03420 | gene30639 | 0.51 | GO:0004674,GO:0005524,GO:0006468,GO:0048544 | ARK3,RK3 | receptor kinase 3 |
| gene30638 | Fvb3:1850406-1855533 | Fvb3 | 1850406 | 1855533 | FvH4_3g03430 | gene30638 | 0.47 | GO:0004674,GO:0005524,GO:0006468,GO:0048544 | ARK3,RK3 | receptor kinase 3 |
| gene30637 | Fvb3:1857375-1862568 | Fvb3 | 1857375 | 1862568 | FvH4_3g03431 | gene30637 | 0.53 | GO:0004674,GO:0005524,GO:0006468,GO:0048544 | S-locus lectin protein kinase family protein | |
| gene30636 | Fvb3:1864042-1867792 | Fvb3 | 1864042 | 1867792 | FvH4_3g03432 | gene30636 | 0.56 | GO:0004674,GO:0005524,GO:0006468,GO:0048544 | S-locus lectin protein kinase family protein | |
| gene30635 | Fvb3:1868486-1872250 | Fvb3 | 1868486 | 1872250 | FvH4_3g03433 | gene30635 | 0.54 | GO:0004674,GO:0005524,GO:0006468,GO:0048544 | SD1-29 | S-domain-1 29 |
| gene39854 | Fvb3:1872986-1875693 | Fvb3 | 1872986 | 1875693 | FvH4_3g03434 | gene39854 | 0.48 | GO:0004672,GO:0005524,GO:0006468 | S-locus lectin protein kinase family protein | |
| gene39855 | Fvb3:1875960-1876356 | Fvb3 | 1875960 | 1876356 | FvH4_3g03435 | gene39855 | 0.5 | | SD1-29 | S-domain-1 29 |
| gene39856 | Fvb3:1879585-1883401 | Fvb3 | 1879585 | 1883401 | FvH4_3g03450 | gene39856 | 0.47 | GO:0004674,GO:0005524,GO:0006468,GO:0048544 | SD1-29 | S-domain-1 29 |
| gene30633 | Fvb3:1885337-1888514 | Fvb3 | 1885337 | 1888514 | FvH4_3g03451 | gene30633 | 0.47 | GO:0004672,GO:0005524,GO:0006468,GO:0048544 | S-locus lectin protein kinase family protein | |
| gene39857 | Fvb3:1890169-1891176 | Fvb3 | 1890169 | 1891176 | FvH4_3g03460 | gene39857 | 0.5 | GO:0004672,GO:0006468 | S-locus lectin protein kinase family protein | |
| gene39858 | Fvb3:1891752-1895554 | Fvb3 | 1891752 | 1895554 | FvH4_3g03461 | gene39858 | 0.47 | GO:0004674,GO:0005524,GO:0006468,GO:0048544 | SD1-29 | S-domain-1 29 |
| gene30630 | Fvb3:1900765-1902550 | Fvb3 | 1900765 | 1902550 | FvH4_3g03480 | gene30630 | 0.1 | GO:0003824,GO:0009058,GO:0030170 | ACS8 | 1-amino-cyclopropane-1-carboxylate synthase 8 |
| gene39859 | Fvb3:1903504-1909697 | Fvb3 | 1903504 | 1909697 | FvH4_3g03481 | gene39859 | 0.5 | GO:0004674,GO:0005524,GO:0006468,GO:0048544 | S-locus lectin protein kinase family protein | |
| gene30631 | Fvb3:1910643-1914082 | Fvb3 | 1910643 | 1914082 | FvH4_3g03482 | gene30631 | 0.46 | GO:0004674,GO:0005524,GO:000 | S-locus lectin | |

| | | | | | | | | | | |
|---|---|---|---|---|---|---|---|---|---|---|
| | | | | | | | | 6468,GO:0048544 | protein kinase family protein | |
| gene39860 | Fvb3:1925559-1926137 | Fvb3 | 1925559 | 1926137 | FvH4_3g03500 | gene39860 | 1 | | | |
| gene39861 | Fvb3:1926399-1930238 | Fvb3 | 1926399 | 1930238 | FvH4_3g03501 | gene39861 | 0.52 | GO:0004674,GO:0005524,GO:0006468,GO:0048544 | SD1-29 | S-domain-1 29 |
| gene39862 | Fvb3:1931602-1934547 | Fvb3 | 1931602 | 1934547 | FvH4_3g03502 | gene39862 | 0.44 | GO:0004672,GO:0005524,GO:0006468,GO:0048544 | S-locus lectin protein kinase family protein | |
| gene30752 | Fvb3:1937726-1941392 | Fvb3 | 1937726 | 1941392 | FvH4_3g03520 | gene30752 | 0.42 | GO:0004674,GO:0005524,GO:0006468,GO:0048544 | SD1-29 | S-domain-1 29 |
| gene30751 | Fvb3:1947703-1951778 | Fvb3 | 1947703 | 1951778 | FvH4_3g03521 | gene30751 | 0.49 | GO:0004674,GO:0005524,GO:0006468,GO:0048544 | SD1-29 | S-domain-1 29 |
| gene30750 | Fvb3:1953397-1955959 | Fvb3 | 1953397 | 1955959 | FvH4_3g03530 | gene30750 | 0.17 | GO:0003677,GO:0006355 | LFY,LFY3 | floral meristem identity control protein LEAFY (LFY) |
| gene30749 | Fvb3:1957303-1959663 | Fvb3 | 1957303 | 1959663 | FvH4_3g03540 | gene30749 | 0.51 | GO:0003677,GO:0006355 | ANAC098,ATCUC2,CUC2 | NAC (No Apical Meristem) domain transcriptional regulator superfamily protein |
| gene39863 | Fvb3:1968041-1969872 | Fvb3 | 1968041 | 1969872 | FvH4_3g03541 | gene39863 | 0.47 | GO:0004672,GO:0005524,GO:0006468 | S-locus protein kinase, putative | |
| gene39864 | Fvb3:1971043-1972422 | Fvb3 | 1971043 | 1972422 | FvH4_3g03542 | gene39864 | 1 | | | Gag-Pol-related retrotransposon family protein |
| gene39865 | Fvb3:1972940-1973908 | Fvb3 | 1972940 | 1973908 | FvH4_3g03543 | gene39865 | 1 | | | Reverse transcriptase (RNA-dependent DNA polymerase) |
| gene30748 | Fvb3:1977342-1981619 | Fvb3 | 1977342 | 1981619 | FvH4_3g03560 | gene30748 | 0.39 | GO:0004674,GO:0005524,GO:0006468,GO:0048544 | SD1-29 | S-domain-1 29 |
| gene36110 | Fvb3:1982582-1984962 | Fvb3 | 1982582 | 1984962 | FvH4_3g03561 | gene36110 | 1 | | CRK8 | cysteine-rich RLK (RECEPTOR-like protein kinase) 8 |
| gene30746 | Fvb3:1993305-1994378 | Fvb3 | 1993305 | 1994378 | FvH4_3g03580 | gene30746 | 1 | | ANAC100,ATNAC5,NAC100 | NAC domain containing protein 100 |
| gene30745 | Fvb3:2002057-2004128 | Fvb3 | 2002057 | 2004128 | FvH4_3g03581 | gene30745 | 0.44 | GO:0048544 | SD1-29 | S-domain-1 29 |
| gene30744 | Fvb3:2005205-2010120 | Fvb3 | 2005205 | 2010120 | FvH4_3g03590 | gene30744 | 0.25 | GO:0004674,GO:0005524,GO:0006468,GO:0048544 | B120 | S-locus lectin protein kinase family protein |
| gene30743 | Fvb3:2012052-2012596 | Fvb3 | 2012052 | 2012596 | FvH4_3g03600 | gene30743 | 0.15 | GO:0005509 | | Calcium-binding EF-hand family protein |
| gene39866 | Fvb3:2017693-2018750 | Fvb3 | 2017693 | 2018750 | FvH4_3g03601 | gene39866 | 1 | | | |
| gene36111 | Fvb3:2025184-2026088 | Fvb3 | 2025184 | 2026088 | FvH4_3g03610 | gene36111 | 0 | GO:0005509 | | Calcium-binding EF-hand family protein |

| | | | | | | | | | | |
|---|---|---|---|---|---|---|---|---|---|---|
| gene36112 | Fvb3:2032026-2041011 | Fvb3 | 2032026 | 2041011 | FvH4_3g03620 | gene36112 | 0.12 | GO:0000155,GO:0000160,GO:0016310 | AHK2,HK2 | histidine kinase 2 |
| gene30741 | Fvb3:2041916-2043479 | Fvb3 | 2041916 | 2043479 | FvH4_3g03630 | gene30741 | 0.37 | GO:0000977,GO:0003700,GO:0005634,GO:0045944,GO:0046983 | AGL24 | AGAMOUS-like 24 |
| gene30740 | Fvb3:2045154-2047400 | Fvb3 | 2045154 | 2047400 | FvH4_3g03640 | gene30740 | 0 | GO:0005622,GO:0008270 | B-box type zinc finger family protein | |
| gene30739 | Fvb3:2047356-2051306 | Fvb3 | 2047356 | 2051306 | FvH4_3g03650 | gene30739 | 0.16 | GO:0006950 | | Adenine nucleotide alpha hydrolases-like superfamily protein |
| gene36113 | Fvb3:2051965-2054538 | Fvb3 | 2051965 | 2054538 | FvH4_3g03660 | gene36113 | 0 | GO:0061630 | | phosphoinositide binding |
| gene30737 | Fvb3:2055536-2058092 | Fvb3 | 2055536 | 2058092 | FvH4_3g03670 | gene30737 | 0.15 | | | PLAC8 family protein |
| gene30736 | Fvb3:2059475-2061075 | Fvb3 | 2059475 | 2061075 | FvH4_3g03680 | gene30736 | 0.07 | GO:0003677 | ATMYB58,MYB58 | myb domain protein 58 |
| gene30735 | Fvb3:2063791-2067097 | Fvb3 | 2063791 | 2067097 | FvH4_3g03690 | gene30735 | 0.01 | | | Protein of unknown function (DUF594) |
| gene30734 | Fvb3:2067408-2069341 | Fvb3 | 2067408 | 2069341 | FvH4_3g03700 | gene30734 | 0.26 | GO:0005615 | | Serine protease inhibitor (SERPIN) family protein |
| gene30732 | Fvb3:2075516-2077533 | Fvb3 | 2075516 | 2077533 | FvH4_3g03710 | gene30732 | 0.25 | GO:0005615 | | Serine protease inhibitor (SERPIN) family protein |
| gene30731 | Fvb3:2079582-2081684 | Fvb3 | 2079582 | 2081684 | FvH4_3g03720 | gene30731 | 0.04 | GO:0005634,GO:0051726 | CYCD6;1 | Cyclin D6;1 |
| gene30730 | Fvb3:2087041-2089912 | Fvb3 | 2087041 | 2089912 | FvH4_3g03730 | gene30730 | 1 | GO:0005515 | | Octicosapeptide/Phox/Bem1p family protein |
| gene30729 | Fvb3:2093477-2096784 | Fvb3 | 2093477 | 2096784 | FvH4_3g03740 | gene30729 | 0.36 | GO:0005337,GO:0016021,GO:1901642 | ATENT3,ENT3,FUR1 | Major facilitator superfamily protein |
| gene30728 | Fvb3:2097824-2100941 | Fvb3 | 2097824 | 2100941 | FvH4_3g03750 | gene30728 | 0.34 | GO:0005337,GO:0016021,GO:1901642 | ATENT3,ENT3,FUR1 | Major facilitator superfamily protein |
| gene39867 | Fvb3:2101296-2102656 | Fvb3 | 2101296 | 2102656 | FvH4_3g03751 | gene39867 | 1 | GO:0003676,GO:0004523 | Polynucleotidyl transferase, ribonuclease H-like superfamily protein | |
| gene30727 | Fvb3:2108104-2114350 | Fvb3 | 2108104 | 2114350 | FvH4_3g03760 | gene30727 | 0.03 | | B160 | Zinc finger, RING-type;Transcription factor jumonji/aspartyl beta-hydroxylase |
| gene30726 | Fvb3:2114368-2118237 | Fvb3 | 2114368 | 2118237 | FvH4_3g03770 | gene30726 | 0.12 | | | Protein kinase superfamily protein |
| gene30725 | Fvb3:2123941-2125668 | Fvb3 | 2123941 | 2125668 | FvH4_3g03780 | gene30725 | 0 | GO:0003677 | ATM4,ATMYB102,MYB102 | MYB-like 102 |

| | | | | | | | | | | |
|---|---|---|---|---|---|---|---|---|---|---|
| gene30724 | Fvb3:2130409-2133928 | Fvb3 | 2130409 | 2133928 | FvH4_3g03790 | gene30724 | 0.14 | GO:0046854 | | Inositol monophosphatase family protein |
| gene36114 | Fvb3:2134381-2135089 | Fvb3 | 2134381 | 2135089 | FvH4_3g03800 | gene36114 | 0 | | | |
| gene30723 | Fvb3:2136348-2137154 | Fvb3 | 2136348 | 2137154 | FvH4_3g03810 | gene30723 | 1 | | | Wound-responsive family protein |
| gene39868 | Fvb3:2138534-2139174 | Fvb3 | 2138534 | 2139174 | FvH4_3g03820 | gene39868 | 1 | | | |
| gene30722 | Fvb3:2139906-2141834 | Fvb3 | 2139906 | 2141834 | FvH4_3g03821 | gene30722 | 1 | | | |
| gene30720 | Fvb3:2148137-2148516 | Fvb3 | 2148137 | 2148516 | FvH4_3g03840 | gene30720 | 1 | GO:0009116 | | uridine 5'-monophosphate synthase / UMP synthase (PYRE-F) (UMPS) |
| gene30719 | Fvb3:2155134-2156798 | Fvb3 | 2155134 | 2156798 | FvH4_3g03870 | gene30719 | 0 | | | |
| gene30718 | Fvb3:2158074-2161938 | Fvb3 | 2158074 | 2161938 | FvH4_3g03880 | gene30718 | 0.2 | GO:0005789 | | PapD-like superfamily protein |
| gene30717 | Fvb3:2164174-2164943 | Fvb3 | 2164174 | 2164943 | FvH4_3g03881 | gene30717 | 1 | | | |
| gene30716 | Fvb3:2166172-2166854 | Fvb3 | 2166172 | 2166854 | FvH4_3g03882 | gene30716 | 1 | | | |
| gene30715 | Fvb3:2172524-2175538 | Fvb3 | 2172524 | 2175538 | FvH4_3g03900 | gene30715 | 0.08 | GO:0016021,GO:0022857,GO:0055085 | ATSTP1, STP1 | sugar transporter 1 |
| gene30714 | Fvb3:2180550-2183354 | Fvb3 | 2180550 | 2183354 | FvH4_3g03910 | gene30714 | 1 | GO:0005515,GO:0006396 | Tetratricopeptide repeat (TPR)-like superfamily protein | |
| gene30713 | Fvb3:2183855-2184805 | Fvb3 | 2183855 | 2184805 | FvH4_3g03920 | gene30713 | 1 | | | |
| gene30712 | Fvb3:2185797-2187518 | Fvb3 | 2185797 | 2187518 | FvH4_3g03930 | gene30712 | 1 | GO:0005515 | | ARM repeat superfamily protein |
| gene30711 | Fvb3:2192859-2197338 | Fvb3 | 2192859 | 2197338 | FvH4_3g03931 | gene30711 | 1 | | | |
| gene30710 | Fvb3:2197836-2201324 | Fvb3 | 2197836 | 2201324 | FvH4_3g03932 | gene30710 | 1 | | | zinc knuckle (CCHC-type) family protein |
| gene30709 | Fvb3:2202893-2205067 | Fvb3 | 2202893 | 2205067 | FvH4_3g03940 | gene30709 | 0 | GO:0005515 | | F-box family protein |
| gene30708 | Fvb3:2207665-2208496 | Fvb3 | 2207665 | 2208496 | FvH4_3g03950 | gene30708 | 0.13 | GO:0007017,GO:0030286 | Dynein light chain type 1 family protein | |
| gene30707 | Fvb3:2209479-2214082 | Fvb3 | 2209479 | 2214082 | FvH4_3g03960 | gene30707 | 0.09 | GO:0005509,GO:0016491,GO:0055114 | NDB2 | NAD(P)H dehydrogenase B2 |
| gene30706 | Fvb3:2217498-2218328 | Fvb3 | 2217498 | 2218328 | FvH4_3g03970 | gene30706 | 1 | | | |
| gene30705 | Fvb3:2220247-2223648 | Fvb3 | 2220247 | 2223648 | FvH4_3g03980 | gene30705 | 0.02 | GO:0005515 | | |
| gene30704 | Fvb3:2224636-2225868 | Fvb3 | 2224636 | 2225868 | FvH4_3g03990 | gene30704 | 0.01 | GO:0005515 | UFO | F-box family protein |
| gene30703 | Fvb3:2228125-2230499 | Fvb3 | 2228125 | 2230499 | FvH4_3g04000 | gene30703 | 0.11 | | | SGNH hydrolase-type esterase superfamily protein |
| gene30702 | Fvb3:2233254-2235273 | Fvb3 | 2233254 | 2235273 | FvH4_3g04010 | gene30702 | 0.16 | | DFL1,GH3.6 | Auxin-responsive GH3 family protein |
| gene30701 | Fvb3:2238784-2240383 | Fvb3 | 2238784 | 2240383 | FvH4_3g04011 | gene30701 | 1 | | | |

| | | | | | | | | | | |
|---|---|---|---|---|---|---|---|---|---|---|
| gene30700 | Fvb3:2241443-2244542 | Fvb3 | 2241443 | 2244542 | FvH4_3g04012 | gene30700 | 1 | | ARP2,ATARP2,WRM | actin related protein 2 |
| gene30699 | Fvb3:2245250-2246484 | Fvb3 | 2245250 | 2246484 | FvH4_3g04020 | gene30699 | 0.02 | GO:0005515 | | F-box family protein |
| gene30698 | Fvb3:2248238-2251991 | Fvb3 | 2248238 | 2251991 | FvH4_3g04030 | gene30698 | 0.07 | GO:0005515 | | Transducin/WD40 repeat-like superfamily protein |
| gene30697 | Fvb3:2252416-2252727 | Fvb3 | 2252416 | 2252727 | FvH4_3g04031 | gene30697 | 1 | | | |
| gene30696 | Fvb3:2254908-2260020 | Fvb3 | 2254908 | 2260020 | FvH4_3g04040 | gene30696 | 0.22 | GO:0008531,GO:0009231,GO:0016787 | ATFMN/FHY,FMN/FHY | riboflavin kinase/FMN hydrolase |
| gene30695 | Fvb3:2260488-2269210 | Fvb3 | 2260488 | 2269210 | FvH4_3g04050 | gene30695 | 0.14 | GO:0005515,GO:0005680,GO:0030071,GO:0031145 | Transducin/WD40 repeat-like superfamily protein | |
| gene30694 | Fvb3:2267614-2272975 | Fvb3 | 2267614 | 2272975 | FvH4_3g04060 | gene30694 | 0.11 | GO:0005516,GO:0006950 | Calmodulin-binding protein | |
| gene30693 | Fvb3:2273592-2276556 | Fvb3 | 2273592 | 2276556 | FvH4_3g04070 | gene30693 | 0.21 | | | |
| gene30692 | Fvb3:2279559-2283961 | Fvb3 | 2279559 | 2283961 | FvH4_3g04090 | gene30692 | 0.15 | GO:0003951,GO:0008152 | SPHK1 | sphingosine kinase 1 |
| gene30691 | Fvb3:2288504-2292230 | Fvb3 | 2288504 | 2292230 | FvH4_3g04100 | gene30691 | 0.16 | GO:0003824,GO:0005992 | Haloacid dehalogenase-like hydrolase (HAD) superfamily protein | |
| gene36115 | Fvb3:2296892-2298138 | Fvb3 | 2296892 | 2298138 | FvH4_3g04110 | gene36115 | 0.13 | | | |
| gene36116 | Fvb3:2303454-2311128 | Fvb3 | 2303454 | 2311128 | FvH4_3g04120 | gene36116 | 0.08 | GO:0003677,GO:0008270 | HSI2-L1,HSL1,VAL2 | HSI2-like 1 |
| gene30689 | Fvb3:2311925-2313808 | Fvb3 | 2311925 | 2313808 | FvH4_3g04130 | gene30689 | 0.18 | GO:0005484,GO:0006886,GO:0016020,GO:0016192 | ATSYP124,SYP124 | syntaxin of plants 124 |
| gene30688 | Fvb3:2313986-2316242 | Fvb3 | 2313986 | 2316242 | FvH4_3g04140 | gene30688 | 0.03 | GO:0000813,GO:0032509 | VPS28-1,VPS28-2 | Vacuolar protein sorting-associated protein VPS28 family protein |
| gene30687 | Fvb3:2316784-2321613 | Fvb3 | 2316784 | 2321613 | FvH4_3g04150 | gene30687 | 0.04 | GO:0005515,GO:0006396 | Tetratricopeptide repeat (TPR)-like superfamily protein | |
| gene30685 | Fvb3:2322710-2324665 | Fvb3 | 2322710 | 2324665 | FvH4_3g04160 | gene30685 | 0.01 | | | Protein of unknown function (DUF761) |
| gene30684 | Fvb3:2330051-2330996 | Fvb3 | 2330051 | 2330996 | FvH4_3g04161 | gene30684 | 1 | | | |
| gene30683 | Fvb3:2331651-2335381 | Fvb3 | 2331651 | 2335381 | FvH4_3g04170 | gene30683 | 0.03 | GO:0003677,GO:0006355 | LFY,LFY3 | floral meristem identity control protein LEAFY (LFY) |

| | | | | | | | | | | |
|---|---|---|---|---|---|---|---|---|---|---|
| gene30682 | Fvb3:2337237-2339099 | Fvb3 | 2337237 | 2339099 | FvH4_3g04180 | gene30682 | 0.12 | GO:0003824,GO:0009058,GO:0030170 | ACS8 | 1-amino-cyclopropane-1-carboxylate synthase 8 |
| gene39869 | Fvb3:2339390-2340260 | Fvb3 | 2339390 | 2340260 | FvH4_3g04190 | gene39869 | 0.22 | GO:0003735,GO:0005840,GO:0006412 | Ribosomal protein L11 family protein | |
| gene36117 | Fvb3:2345654-2347251 | Fvb3 | 2345654 | 2347251 | FvH4_3g04191 | gene36117 | 1 | | | |
| gene30680 | Fvb3:2352139-2358276 | Fvb3 | 2352139 | 2358276 | FvH4_3g04200 | gene30680 | 0.24 | | ATSNF4, SNF4 | homolog of yeast sucrose nonfermenting 4 |
| gene39870 | Fvb3:2359667-2360037 | Fvb3 | 2359667 | 2360037 | FvH4_3g04210 | gene39870 | 1 | | DVL18,RTFL5 | ROTUNDIFOLIA like 5 |
| gene30679 | Fvb3:2361295-2365490 | Fvb3 | 2361295 | 2365490 | FvH4_3g04220 | gene30679 | 0.18 | GO:0005096 | AGD5,NEV | ARF-GAP domain 5 |
| gene39871 | Fvb3:2366518-2367865 | Fvb3 | 2366518 | 2367865 | FvH4_3g04230 | gene39871 | 0.44 | | | |
| gene39872 | Fvb3:2368453-2369188 | Fvb3 | 2368453 | 2369188 | FvH4_3g04231 | gene39872 | 0.44 | | | Protein of unknown function (DUF300) |
| gene30678 | Fvb3:2369280-2370216 | Fvb3 | 2369280 | 2370216 | FvH4_3g04232 | gene30678 | 0.37 | | | Protein of unknown function (DUF300) |
| gene30677 | Fvb3:2372687-2378024 | Fvb3 | 2372687 | 2378024 | FvH4_3g04233 | gene30677 | 1 | | | |
| gene30676 | Fvb3:2386233-2387546 | Fvb3 | 2386233 | 2387546 | FvH4_3g04240 | gene30676 | 0.44 | | | |
| gene30675 | Fvb3:2388143-2389916 | Fvb3 | 2388143 | 2389916 | FvH4_3g04241 | gene30675 | 0.29 | | | Protein of unknown function (DUF300) |
| gene30674 | Fvb3:2392811-2393323 | Fvb3 | 2392811 | 2393323 | FvH4_3g04242 | gene30674 | 0.8 | | | |
| gene30673 | Fvb3:2394024-2395850 | Fvb3 | 2394024 | 2395850 | FvH4_3g04250 | gene30673 | 0.13 | | | Putative lysine decarboxylase family protein |
| gene30672 | Fvb3:2402870-2403688 | Fvb3 | 2402870 | 2403688 | FvH4_3g04260 | gene30672 | 1 | | | Protein of unknown function (DUF1191) |
| gene30482 | Fvb3:2414813-2419554 | Fvb3 | 2414813 | 2419554 | FvH4_3g04270 | gene30482 | 0.03 | GO:0005634,GO:0006355,GO:0043565 | BUM,BUM1,SHL,STM,WAM,WAM1 | KNOX/ELK homeobox transcription factor |
| gene36118 | Fvb3:2434240-2445242 | Fvb3 | 2434240 | 2445242 | FvH4_3g04271 | gene36118 | 1 | GO:0046983 | | BED zinc finger ;hAT family dimerisation domain |
| gene39873 | Fvb3:2437887-2441102 | Fvb3 | 2437887 | 2441102 | FvH4_3g04272 | gene39873 | 1 | | | |
| gene39874 | Fvb3:2441838-2442306 | Fvb3 | 2441838 | 2442306 | FvH4_3g04273 | gene39874 | 1 | | | |
| gene34398 | Fvb3:2445503-2447544 | Fvb3 | 2445503 | 2447544 | FvH4_3g04274 | gene34398 | 1 | | | |
| gene39875 | Fvb3:2447567-2449053 | Fvb3 | 2447567 | 2449053 | FvH4_3g04275 | gene39875 | 1 | | | |
| gene34397 | Fvb3:2451076-2451596 | Fvb3 | 2451076 | 2451596 | FvH4_3g04276 | gene34397 | 1 | | | |
| gene39876 | Fvb3:2451879-2453196 | Fvb3 | 2451879 | 2453196 | FvH4_3g04277 | gene39876 | 1 | | | |
| gene30480 | Fvb3:2454125-2454500 | Fvb3 | 2454125 | 2454500 | FvH4_3g04278 | gene30480 | 1 | | | |
| gene39877 | Fvb3:2455276-2456618 | Fvb3 | 2455276 | 2456618 | FvH4_3g04279 | gene39877 | 1 | | | |
| gene30479 | Fvb3:2456669-2457157 | Fvb3 | 2456669 | 2457157 | FvH4_3g04280 | gene30479 | 1 | | | |

| | | | | | | | | | | |
|---|---|---|---|---|---|---|---|---|---|---|
| gene39878 | Fvb3:2457754-2458089 | Fvb3 | 2457754 | 2458089 | FvH4_3g04281 | gene39878 | 1 | | PRA1.B4 | prenylated RAB acceptor 1.B4 |
| gene30478 | Fvb3:2463114-2463608 | Fvb3 | 2463114 | 2463608 | FvH4_3g04290 | gene30478 | 0.35 | GO:0046983 | | basic helix-loop-helix (bHLH) DNA-binding family protein |
| gene30477 | Fvb3:2467937-2470515 | Fvb3 | 2467937 | 2470515 | FvH4_3g04300 | gene30477 | 0.27 | | | Putative lysine decarboxylase family protein |
| gene30475 | Fvb3:2478741-2480461 | Fvb3 | 2478741 | 2480461 | FvH4_3g04320 | gene30475 | 0.23 | GO:0016491,GO:0055114 | oxidoreductase, zinc-binding dehydrogenase family protein | |
| gene30474 | Fvb3:2481728-2482159 | Fvb3 | 2481728 | 2482159 | FvH4_3g04330 | gene30474 | 0.69 | | | RING/U-box superfamily protein |
| gene30473 | Fvb3:2483914-2485758 | Fvb3 | 2483914 | 2485758 | FvH4_3g04340 | gene30473 | 0.32 | GO:0016491,GO:0055114 | oxidoreductase, zinc-binding dehydrogenase family protein | |
| gene30472 | Fvb3:2486427-2490004 | Fvb3 | 2486427 | 2490004 | FvH4_3g04350 | gene30472 | 0.29 | GO:0003676,GO:0004519,GO:0006308 | ENDO4 | endonuclease 4 |
| gene30471 | Fvb3:2490330-2493176 | Fvb3 | 2490330 | 2493176 | FvH4_3g04360 | gene30471 | 0.43 | GO:0003676,GO:0004519,GO:0006308 | BFN1,ENDO1 | bifunctional nuclease i |
| gene30470 | Fvb3:2493649-2503227 | Fvb3 | 2493649 | 2503227 | FvH4_3g04370 | gene30470 | 0.21 | GO:0016491,GO:0055114 | ATNDI1,NDA1 | alternative NAD(P)H dehydrogenase 1 |
| gene30469 | Fvb3:2498348-2498629 | Fvb3 | 2498348 | 2498629 | FvH4_3g04371 | gene30469 | 0.28 | | | |
| gene30468 | Fvb3:2501261-2501731 | Fvb3 | 2501261 | 2501731 | FvH4_3g04372 | gene30468 | 1 | | | |
| gene30467 | Fvb3:2502379-2503545 | Fvb3 | 2502379 | 2503545 | FvH4_3g04380 | gene30467 | 0.84 | | | |
| gene30466 | Fvb3:2506864-2508628 | Fvb3 | 2506864 | 2508628 | FvH4_3g04381 | gene30466 | 1 | | | |
| gene30465 | Fvb3:2510283-2511340 | Fvb3 | 2510283 | 2511340 | FvH4_3g04390 | gene30465 | 1 | | | alpha/beta-Hydrolases superfamily protein |
| gene30464 | Fvb3:2516385-2518781 | Fvb3 | 2516385 | 2518781 | FvH4_3g04400 | gene30464 | 0 | GO:0003677 | PGA6,WUS,WUS1 | Homeodomain-like superfamily protein |
| gene30463 | Fvb3:2519250-2521670 | Fvb3 | 2519250 | 2521670 | FvH4_3g04401 | gene30463 | 1 | | | |
| gene30462 | Fvb3:2526433-2539920 | Fvb3 | 2526433 | 2539920 | FvH4_3g04410 | gene30462 | 0.25 | GO:0003824,GO:0006284,GO:0051539 | DME | HhH-GPD base excision DNA repair family protein |
| gene30460 | Fvb3:2542466-2545728 | Fvb3 | 2542466 | 2545728 | FvH4_3g04430 | gene30460 | 0.04 | GO:0006790,GO:0008441,GO:0046854 | AHL,ATAHL,HL | HAL2-like |
| gene30459 | Fvb3:2548136-2549101 | Fvb3 | 2548136 | 2549101 | FvH4_3g04450 | gene30459 | 1 | | | |
| gene30458 | Fvb3:2549192-2553125 | Fvb3 | 2549192 | 2553125 | FvH4_3g04440 | gene30458 | 0.12 | GO:0015031,GO:0016021 | SC3 | secretory carrier 3 |
| gene30457 | Fvb3:2553666-2558651 | Fvb3 | 2553666 | 2558651 | FvH4_3g04460 | gene30457 | 0.06 | | | Protein of unknown function (DUF707) |
| gene30456 | Fvb3:2561199-2564257 | Fvb3 | 2561199 | 2564257 | FvH4_3g04470 | gene30456 | 0.01 | GO:0005515 | BRD4 | bromodomain 4 |

| | | | | | | | | | | |
|---|---|---|---|---|---|---|---|---|---|---|
| gene36119 | Fvb3:2565092-2573380 | Fvb3 | 2565092 | 2573380 | FvH4_3g04490 | gene36119 | 0.06 | GO:0008017,GO:0008352,GO:0051013 | Transducin/WD40 repeat-like superfamily protein | |
| gene36120 | Fvb3:2575125-2581786 | Fvb3 | 2575125 | 2581786 | FvH4_3g04500 | gene36120 | 0.06 | GO:0004672,GO:0005515,GO:0005524,GO:0006468 | SRF3 | STRUBBELIG-receptor family 3 |
| gene30454 | Fvb3:2584220-2585013 | Fvb3 | 2584220 | 2585013 | FvH4_3g04510 | gene30454 | 0.27 | | | |
| gene30453 | Fvb3:2587117-2588921 | Fvb3 | 2587117 | 2588921 | FvH4_3g04520 | gene30453 | 1 | | | |
| gene30452 | Fvb3:2591487-2593892 | Fvb3 | 2591487 | 2593892 | FvH4_3g04521 | gene30452 | 1 | GO:0003677,GO:0046983 | BED zinc finger ;hAT family dimerisation domain | |
| gene30451 | Fvb3:2595061-2596221 | Fvb3 | 2595061 | 2596221 | FvH4_3g04530 | gene30451 | 0.01 | | | |
| gene30450 | Fvb3:2597499-2599924 | Fvb3 | 2597499 | 2599924 | FvH4_3g04540 | gene30450 | 0.29 | | | |
| gene30449 | Fvb3:2603071-2603504 | Fvb3 | 2603071 | 2603504 | FvH4_3g04541 | gene30449 | 1 | | | |
| gene30448 | Fvb3:2605809-2608029 | Fvb3 | 2605809 | 2608029 | FvH4_3g04550 | gene30448 | 0.19 | | | lysine decarboxylase family protein |
| gene30447 | Fvb3:2612473-2613372 | Fvb3 | 2612473 | 2613372 | FvH4_3g04560 | gene30447 | 1 | | | |
| gene30446 | Fvb3:2613547-2616827 | Fvb3 | 2613547 | 2616827 | FvH4_3g04570 | gene30446 | 0.09 | GO:0005515 | | LisH and RanBPM domains containing protein |
| gene30445 | Fvb3:2618718-2631807 | Fvb3 | 2618718 | 2631807 | FvH4_3g04580 | gene30445 | 0.04 | GO:0005524 | EDA16 | SNF2 domain-containing protein / helicase domain-containing protein / zinc finger protein-related |
| gene30444 | Fvb3:2627189-2628148 | Fvb3 | 2627189 | 2628148 | FvH4_3g04590 | gene30444 | 0.02 | | | alpha/beta-Hydrolases superfamily protein |
| gene30443 | Fvb3:2633585-2637467 | Fvb3 | 2633585 | 2637467 | FvH4_3g04600 | gene30443 | 0.18 | GO:0004185,GO:0006508 | scpl31 | serine carboxypeptidase-like 31 |
| gene30442 | Fvb3:2638003-2640481 | Fvb3 | 2638003 | 2640481 | FvH4_3g04610 | gene30442 | 0.05 | GO:0009690,GO:0019139,GO:0055114,GO:0071949 | ATCKX5, CKX7 | cytokinin oxidase 7 |
| gene30441 | Fvb3:2645673-2649644 | Fvb3 | 2645673 | 2649644 | FvH4_3g04611 | gene30441 | 1 | | | |
| gene30440 | Fvb3:2651305-2654139 | Fvb3 | 2651305 | 2654139 | FvH4_3g04620 | gene30440 | 0.18 | GO:0004185,GO:0006508 | scpl31 | serine carboxypeptidase-like 31 |
| gene30439 | Fvb3:2655643-2657423 | Fvb3 | 2655643 | 2657423 | FvH4_3g04630 | gene30439 | 0.13 | GO:0003677,GO:0006355 | anac025,NAC025 | NAC domain containing protein 25 |
| gene30438 | Fvb3:2662560-2663368 | Fvb3 | 2662560 | 2663368 | FvH4_3g04640 | gene30438 | 1 | GO:0005515,GO:0007165 | Toll-Interleukin-Resistance (TIR) domain | |

| | | | | | | | | | | |
|---|---|---|---|---|---|---|---|---|---|---|
| | | | | | | | | | family protein | |
| gene30437 | Fvb3:2667733-2668380 | Fvb3 | 2667733 | 2668380 | FvH4_3g04641 | gene30437 | 1 | | | |
| gene36121 | Fvb3:2668901-2673182 | Fvb3 | 2668901 | 2673182 | FvH4_3g04650 | gene36121 | 0.09 | | | disease resistance protein (TIR class), putative |
| gene36122 | Fvb3:2673192-2677479 | Fvb3 | 2673192 | 2677479 | FvH4_3g04651 | gene36122 | 0.09 | GO:0003676,GO:0006364,GO:0008408 | Exonuclease family protein | |

**Supplemental Table 8.** Phenotypic data of the two sets of individual with extremes phenotypes for pelargonidin-3-glucoside and studied using microarray. The choice of genotypes was based on the level of PgGs in 2010 and according to the genotyping with the marker located in the peak of QTL: AX-89826440-M3a.

| Genotype | year | Level | Ant | PgGs | PgGsM | PgRs | CyGs | AfPgGs | Fvo | KGs | KGn | KCoGs | QGn | F3ol | Cat | CatCat | AfCat | AfGs | UnK | UnK1 | UnK2 | ANTHc | FLAVc | PHENc | FRAP | TEAC | COLOUR |
|---|---|---|---|---|---|---|---|---|---|---|---|---|---|---|---|---|---|---|---|---|---|---|---|---|---|---|---|
| 6 | 2010 | high | 40.49 | 37.67 | 0.528 | 1.134 | 1.15 | 0.013 | 0.616 | 0.169 | 0.12 | 0.106 | 0.221 | 1.706 | 0.327 | 0.757 | 0.083 | 0.539 | 1.161 | 0.896 | 0.265 | 965.8 | 630.5 | 1.771 | 8.665 | 14.07 | |
| 7 | 2010 | high | 28.84 | 26.79 | 0.16 | 1.324 | 0.546 | 0.016 | 0.442 | 0.183 | 0.082 | 0.11 | 0.067 | 1.983 | 0.223 | 0.478 | 0.044 | 1.238 | 0.874 | 0.685 | 0.189 | 486 | 577.6 | 1.477 | 8.285 | 14.73 | |
| 34 | 2010 | high | 38.89 | 36.67 | 0 | 1.325 | 0.881 | 0.016 | 0.414 | 0.153 | 0.052 | 0.172 | 0.037 | 1.19 | 0.206 | 0.458 | 0.052 | 0.474 | 1.3 | 0.961 | 0.339 | 1103 | 585 | 1.576 | 8.152 | 14.46 | |
| 36 | 2010 | high | 28.61 | 25.8 | 0.608 | 1.359 | 0.829 | 0.011 | 0.301 | 0.137 | 0.078 | 0.055 | 0.031 | 1.067 | 0.239 | 0.582 | 0.051 | 0.194 | 0.837 | 0.64 | 0.197 | 757.2 | 527.1 | 1.1 | 6.786 | 12.51 | |
| 85 | 2010 | high | 33.67 | 31.4 | 0.174 | 1.275 | 0.809 | 0.011 | 0.313 | 0.095 | 0.074 | 0.099 | 0.045 | 1.098 | 0.243 | 0.578 | 0.068 | 0.208 | 1.025 | 0.767 | 0.258 | 761.1 | 659.7 | 1.64 | 8.964 | 16.19 | |
| 198 | 2010 | high | 36.28 | 33.06 | 0.24 | 2.314 | 0.642 | 0.021 | 0.325 | 0.109 | 0.064 | 0.071 | 0.082 | 2.637 | 0.233 | 0.56 | 0.066 | 1.778 | 0.982 | 0.725 | 0.257 | 1014 | 582.8 | 1.638 | 9.792 | 16.61 | |
| 216 | 2010 | high | 32.15 | 28.59 | 0.275 | 2.894 | 0.37 | 0.02 | 0.325 | 0.091 | 0.063 | 0.132 | 0.04 | 0.866 | 0.207 | 0.409 | 0.045 | 0.205 | 0.905 | 0.691 | 0.215 | 1048 | 405.8 | 1.627 | 7.756 | 14.21 | |
| 16 | 2010 | low | 27.74 | 26.61 | 0.133 | 0.814 | 0.163 | 0.019 | 0.308 | 0.108 | 0.085 | 0.077 | 0.038 | 1.211 | 0.235 | 0.565 | 0.061 | 0.351 | 0.849 | 0.663 | 0.186 | 661.1 | 261.6 | 1.124 | 7.423 | 12.45 | |
| 19 | 2010 | low | 20.97 | 19.69 | 0.091 | 0.957 | 0.232 | 0.006 | 0.17 | 0.037 | 0.055 | 0.045 | 0.033 | 1.093 | 0.216 | 0.458 | 0.041 | 0.378 | 0.6 | 0.477 | 0.123 | 661 | 294.3 | 1.354 | 7.623 | 13.78 | |
| 46 | 2010 | low | 14.97 | 14.08 | 0.015 | 0.734 | 0.13 | 0.01 | 0.168 | 0.065 | 0.029 | 0.051 | 0.023 | 1.178 | 0.246 | 0.476 | 0.048 | 0.408 | 0.465 | 0.356 | 0.109 | 352.5 | 347.7 | 0.892 | 6.751 | 12.4 | |
| 78 | 2010 | low | 18.25 | 16.8 | 0.023 | 1.167 | 0.253 | 0.009 | 0.253 | 0.073 | 0.047 | 0.075 | 0.057 | 1.469 | 0.22 | 0.435 | 0.069 | 0.745 | 0.472 | 0.374 | 0.098 | 515.7 | 482.8 | 1.249 | 6.832 | 13.23 | |
| 94 | 2010 | low | 19.07 | 18.11 | 0 | 0.828 | 0.132 | 0.007 | 0.177 | 0.037 | 0.062 | 0.038 | 0.041 | 0.665 | 0.171 | 0.315 | 0.029 | 0.15 | 0.561 | 0.409 | 0.152 | 476.1 | 386.8 | 1.047 | 6.221 | 12.31 | |
| 111 | 2010 | low | 18.01 | 16.66 | 0 | 1.176 | 0.165 | 0.003 | 0.301 | 0.096 | 0.076 | 0.087 | 0.043 | 1.155 | 0.245 | 0.507 | 0.045 | 0.358 | 0.5 | 0.396 | 0.104 | 672.8 | 456.9 | 1.175 | 6.907 | 13.18 | |
| 180 | 2010 | low | 15.85 | 14.63 | 0 | 1.091 | 0.125 | 0.004 | 0.182 | 0.062 | 0.042 | 0.058 | 0.021 | 0.972 | 0.199 | 0.38 | 0.039 | 0.354 | 0.541 | 0.421 | 0.12 | 484.1 | 296.5 | 1.05 | 4.652 | 9.84 | |
| 6 | 2011 | high | 38.87 | 33.59 | 0.356 | 3.969 | 0.488 | 0.47 | 0.354 | 0.155 | 0.099 | 0.065 | 0.035 | 1.713 | 0.316 | 0.708 | 0.092 | 0.596 | 1.214 | 0.932 | 0.282 | 1085 | 447.2 | 4.428 | | 35.48 | 5.75 |
| 7 | 2011 | high | 28.6 | 24.74 | 0.373 | 2.41 | 0.714 | 0.365 | 0.4 | 0.188 | 0.09 | 0.082 | 0.04 | 1.525 | 0.244 | 0.525 | 0.048 | 0.709 | 0.994 | 0.745 | 0.249 | 744.9 | 358.3 | 2.908 | | 29.09 | 4 |
| 34 | 2011 | high | 32.14 | 27.03 | 0 | 4.376 | 0.452 | 0.287 | 0.227 | 0.121 | 0.034 | 0.05 | 0.023 | 1.235 | 0.188 | 0.442 | 0.041 | 0.565 | 0.981 | 0.751 | 0.23 | 1200 | 399.5 | 2.398 | | 33.75 | 5.5 |
| 36 | 2011 | high | 34.04 | 28.49 | 0.963 | 3.687 | 0.52 | 0.385 | 0.282 | 0.155 | 0.071 | 0.036 | 0.02 | 0.915 | 0.195 | 0.461 | 0.052 | 0.207 | 1.034 | 0.81 | 0.224 | 1220 | 469.2 | 2.792 | | 39.98 | 6 |
| 85 | 2011 | high | 34.19 | 28.51 | 0 | 4.732 | 0.61 | 0.338 | 0.243 | 0.085 | 0.076 | 0.052 | 0.03 | 1.159 | 0.24 | 0.584 | 0.085 | 0.249 | 1.01 | 0.794 | 0.216 | 1052 | 384 | 2.259 | | 33.36 | 3.25 |
| 198 | 2011 | high | 34.17 | 29.74 | 0 | 3.443 | 0.497 | 0.49 | 0.189 | 0.104 | 0.036 | 0.024 | 0.026 | 2.567 | 0.268 | 0.579 | 0.09 | 1.631 | 0.927 | 0.732 | 0.195 | 1028 | 424.7 | 2.426 | | 33.53 | 5.25 |
| 216 | 2011 | high | 36.15 | 28.02 | 1.242 | 6.243 | 0.353 | 0.289 | 0.268 | 0.127 | 0.056 | 0.058 | 0.028 | 1.209 | 0.304 | 0.713 | 0.077 | 0.116 | 0.998 | 0.758 | 0.239 | 1102 | 591.6 | 2.84 | | 39.53 | 6 |
| 16 | 2011 | low | 24.99 | 20.75 | 0.071 | 3.656 | 0.271 | 0.246 | 0.373 | 0.145 | 0.096 | 0.078 | 0.053 | 1.467 | 0.354 | 0.787 | 0.069 | 0.257 | 0.771 | 0.617 | 0.154 | 726.6 | 417.7 | 2.487 | | 36.62 | 2 |
| 19 | 2011 | low | 21.82 | 19.37 | 0 | 2.029 | 0.174 | 0.248 | 0.167 | 0.064 | 0.062 | 0.019 | 0.022 | 1.084 | 0.189 | 0.371 | 0.051 | 0.472 | 0.75 | 0.595 | 0.155 | 765.3 | 390.9 | 2.457 | | 34.48 | 1.75 |
| 46 | 2011 | low | 19.57 | 17.46 | 0.101 | 1.547 | 0.306 | 0.152 | 0.271 | 0.093 | 0.058 | 0.064 | 0.056 | 1.576 | 0.284 | 0.704 | 0.057 | 0.531 | 0.483 | 0.37 | 0.114 | 438.5 | 426 | 2.414 | | 33.44 | 0.75 |

| | | | | | | | | | | | | | | | | | | | | | | | | | | |
|---|---|---|---|---|---|---|---|---|---|---|---|---|---|---|---|---|---|---|---|---|---|---|---|---|---|---|---|
| 78 | 2011 | low | 23.45 | 19.63 | 0.074 | 3.185 | 0.324 | 0.237 | 0.259 | 0.116 | 0.047 | 0.05 | 0.045 | 1.538 | 0.284 | 0.619 | 0.098 | 0.536 | 0.603 | 0.489 | 0.115 | 758.3 | 433.6 | 2.366 | | 36.74 | 2.25 |
| 94 | 2011 | low | 22.09 | 19.6 | 0.078 | 2.093 | 0.185 | 0.128 | 0.273 | 0.102 | 0.084 | 0.051 | 0.036 | 0.953 | 0.283 | 0.523 | 0.06 | 0.088 | 0.723 | 0.545 | 0.178 | 607.1 | 366.7 | 2.262 | | 31.06 | 1 |
| 111 | 2011 | low | 27.46 | 22.98 | 0 | 3.414 | 0.858 | 0.207 | 0.297 | 0.148 | 0.054 | 0.06 | 0.035 | 0.837 | 0.18 | 0.401 | 0.045 | 0.211 | 0.8 | 0.625 | 0.174 | 979.7 | 358.2 | 2.203 | | 31.83 | 5.25 |
| 180 | 2011 | low | 24.03 | 19.48 | 0 | 4.115 | 0.228 | 0.204 | 0.237 | 0.103 | 0.072 | 0.039 | 0.023 | 1.126 | 0.229 | 0.481 | 0.067 | 0.349 | 0.663 | 0.525 | 0.137 | 497.2 | 300.7 | 2.047 | | 25.28 | 1.75 |

| year | | Ant.mean | PgGs.mean | PgGsM.mean | PgRs.mean | CyGs.mean | AfPgGs.mean | Fvo.mean | KGs.mean | KGn.mean | KCoGs.mean | QGn.mean | F3ol.mean | Cat.mean | CatCat.mean | AfCat.mean | AfGs.mean | UnK.mean | UnK1.mean | UnK2.mean | ANTHc.mean | FLAVc.mean | PHENc.mean | FRAP.mean | TEAC.mean | |
|---|---|---|---|---|---|---|---|---|---|---|---|---|---|---|---|---|---|---|---|---|---|---|---|---|---|---|
| 2010 | | 4E-05 | 1E-04 | 0.022 | 0.034 | 8E-04 | 0.015 | 0.007 | 0.003 | 0.101 | 0.023 | 0.186 | 0.159 | 0.289 | 0.092 | 0.164 | 0.3 | 2E-04 | 2E-04 | 4E-04 | 0.006 | 6E-04 | 0.002 | 0.006 | 0.009 | |
| 2011 | | 2E-05 | 3E-05 | 0.097 | 0.052 | 0.1 | 7E-04 | 0.739 | 0.209 | 0.898 | 0.941 | 0.119 | 0.317 | 0.82 | 0.807 | 0.624 | 0.292 | 5E-05 | 1E-04 | 5E-05 | 0.001 | 0.142 | 0.099 | | 0.314 | 0.00<br>1 |

**Supplemental Table 9.** List of genes included in the support interval of M3a colour-related QTLs that were tested by microarray. The support interval was on Fvb3 between 1.213489 Mb to 2.673762 Mb. In this region, a total of 232 genes were tested on microarray. More particular attention were provided to three genes, underlined in yellow in this table. These three genes were considered as candidate genes for controlling the M3a colour-related QTLs.

| | | | | | | | | | | | | | | | | | | | | | | |
|---|---|---|---|---|---|---|---|---|---|---|---|---|---|---|---|---|---|---|---|---|---|---|
| | | 19 | 21 | | | | | | 18 | | | | | | 11 | | | | | | | |
| | Genotype | 8 | 6 | 6 | 34 | 36 | 85 | 7 | 0 | 19 | 46 | 78 | 94 | 16 | 1 | | | | | | | |
| | Choice | | | | | | | | | | | | | | | | | | | | | |
| | PgGs+genotyp | hig | hig | hig | hig | hig | hig | hig | lo | lo | lo | lo | lo | lo | lo | red=<0. | FvH4_name_ | LG_v4. | start_v4 | end_v4. | | |
| | e | h | h | h | h | h | h | h | w | w | w | w | w | w | w | 05 | v4.0.a2 | 0.a2 | .0.a2 | 0.a2 | annot_v4.0.a2 | |
| 196 | | 18. | 20. | 24. | 22. | 23. | 18. | 21. | 18. | 22. | 22. | 25. | 21. | 24. | 27. | 0.2169 | FvH4_3g028 | | 151263 | 151388 | | |
| 25 | GENE19521 | 6 | 8 | 8 | 1 | 1 | 8 | 5 | 4 | 7 | 6 | 5 | 3 | 5 | 6 | 2079 | 51 | Fvb3 | 3 | 0 | | 0 |
| 196 | | 42. | | 71. | 55. | 61. | 65. | 55. | 65. | 50. | 55. | 62. | 61. | 53. | 68. | 0.6928 | FvH4_3g028 | | 148284 | 148484 | | |
| 27 | GENE19523 | 2 | 53 | 3 | 9 | 2 | 1 | 8 | 5 | 3 | 9 | 2 | 1 | 5 | 4 | 3635 | 31 | Fvb3 | 9 | 3 | MATE efflux family protein | |
| 196 | | 14. | 17. | 13. | 12. | 17. | 14. | 15. | 13. | 12. | 12. | 12. | 15. | 13. | | 0.1124 | FvH4_3g028 | | 147997 | 148212 | | |
| 28 | GENE19524 | 5 | 8 | 4 | 2 | 3 | 5 | 3 | 4 | 1 | 8 | 8 | 5 | 9 | 14 | 1624 | 30 | Fvb3 | 0 | 7 | cysteine-rich RLK (RECEPTOR-like protein kinase) 25 | |
| 196 | | 21 | 39 | 28 | 56 | 29 | 45 | 38 | 64 | 33 | 72 | 55 | 60 | 42 | 36 | 0.0562 | FvH4_3g028 | | 147594 | 147900 | | |
| 29 | GENE19525 | 2 | 4 | 9 | 4 | 2 | 1 | 2 | 2 | 3 | 2 | 0 | 5 | 3 | 9 | 2445 | 20 | Fvb3 | 0 | 8 | cysteine-rich RLK (RECEPTOR-like protein kinase) 25 | |
| 196 | | 19. | 19. | 26. | 21. | 23. | 18. | 24. | 20. | | 21. | 23. | 24. | 23. | 20. | 0.8373 | FvH4_3g028 | | 147338 | 147394 | | |
| 30 | GENE19526 | 4 | 7 | 2 | 8 | 6 | 6 | 2 | 4 | 17 | 3 | 9 | 9 | 4 | 6 | 8615 | 01 | Fvb3 | 5 | 1 | cysteine-rich RLK (RECEPTOR-like protein kinase) 33 | |
| 196 | | | 21. | 26. | 20. | 20. | 24. | 23. | 23. | 18. | 20. | 23. | 26. | 25. | 30. | 0.3657 | FvH4_3g027 | | 146564 | 146610 | | |
| 31 | GENE19527 | 19 | 7 | 1 | 9 | 7 | 2 | 9 | 4 | 7 | 3 | 8 | 4 | 1 | 6 | 9143 | 90 | Fvb3 | 3 | 1 | MATE efflux family protein | |
| 196 | | 24. | 42. | 42. | 39. | 37. | 31. | 50. | 36. | 27. | 40. | 49. | 56. | | 33. | 0.4632 | FvH4_3g027 | | 145607 | 145866 | | |
| 32 | GENE19528 | 3 | 2 | 6 | 2 | 8 | 1 | 9 | 8 | 9 | 8 | 6 | 5 | 50 | 4 | 2563 | 74 | Fvb3 | 1 | 1 | cysteine-rich RLK (RECEPTOR-like protein kinase) 25 | |
| 196 | | 26. | 54. | 45. | 76. | 39. | | | 82. | 50. | 92. | 75. | 80. | 63. | | 0.0269 | FvH4_3g027 | | 144995 | 145357 | | |
| 34 | GENE19530 | 7 | 8 | 4 | 1 | 3 | 53 | 52 | 7 | 1 | 7 | 9 | 5 | 7 | 51 | 4328 | 73 | Fvb3 | 6 | 1 | cysteine-rich RLK (RECEPTOR-like protein kinase) 25 | |
| 196 | | 15 | 36 | 32 | 53. | 57 | 52. | 25 | 53. | 51. | 48. | 36. | 29. | 45. | 58. | 0.0261 | FvH4_3g027 | | 143779 | 144355 | | |
| 36 | GENE19532 | 6 | 7 | 5 | 5 | 9 | 7 | 4 | 7 | 1 | 8 | 1 | 1 | 7 | 4 | 444 | 72 | Fvb3 | 8 | 4 | cysteine-rich RLK (RECEPTOR-like protein kinase) 10 | |
| 196 | | 28. | 32. | 36. | | 28. | 25. | 33. | 28. | | 31. | 30. | 27. | 35. | 27. | 0.6287 | FvH4_3g027 | | 143111 | 143544 | | |
| 37 | GENE19533 | 1 | 6 | 6 | 33 | 4 | 6 | 5 | 1 | 31 | 1 | 2 | 6 | 7 | 9 | 6617 | 71 | Fvb3 | 0 | 6 | cysteine-rich RLK (RECEPTOR-like protein kinase) 10 | |
| 196 | | 19. | 24. | 21. | 28. | 22. | 27. | 26. | | 25. | 16. | 18. | 14. | 18. | 15. | 0.0248 | FvH4_3g027 | | 142637 | 143030 | | |
| 38 | GENE19534 | 7 | 1 | 9 | 2 | 2 | 8 | 3 | 24 | 8 | 3 | 6 | 9 | 4 | 3 | 5352 | 70 | Fvb3 | 5 | 7 | cysteine-rich RLK (RECEPTOR-like protein kinase) 25 | |
| 196 | | 30. | 56. | 13 | 11 | 25. | 12 | 37. | 53. | 13 | 13 | 78. | 70. | 79. | 14 | 0.2599 | FvH4_3g027 | | 141829 | 142530 | | |
| 41 | GENE19537 | 2 | 2 | 0 | 1 | 5 | 3 | 7 | 5 | 9 | 5 | 8 | 7 | 4 | 5 | 5151 | 54 | Fvb3 | 8 | 1 | cysteine-rich RLK (RECEPTOR-like protein kinase) 25 | |
| 196 | | 8.9 | 11. | | 8.9 | 10. | 11. | 9.4 | 11. | 8.7 | 9.7 | 9.9 | 9.0 | 9.7 | 13. | 0.6492 | FvH4_3g027 | | 140672 | 140799 | | |
| 42 | GENE19538 | 9 | 5 | 9.3 | 1 | 5 | 5 | 8 | 8 | 1 | 4 | 7 | 3 | 3 | 7 | 2463 | 53 | Fvb3 | 4 | 4 | | 0 |
| 196 | | 22. | 28. | 23. | | 23. | 29. | | 25. | 29. | 22. | 24. | 23. | 21. | 28. | 0.9993 | FvH4_3g027 | | 139882 | 140146 | | |
| 43 | GENE19539 | 2 | 3 | 5 | 24 | 7 | 1 | 24 | 1 | 8 | 1 | 8 | 2 | 3 | 5 | 1019 | 52 | Fvb3 | 3 | 4 | cysteine-rich RLK (RECEPTOR-like protein kinase) 25 | |
| 196 | | 12 | 32 | 10 | 15 | 86. | 10 | 89. | | 70. | 85. | 46. | 72. | 89. | 60. | 0.0785 | FvH4_3g027 | | 138676 | 139010 | | |
| 45 | GENE19541 | 0 | 5 | 3 | 3 | 7 | 4 | 7 | 80 | 9 | 8 | 9 | 5 | 6 | 2 | 5174 | 31 | Fvb3 | 2 | 2 | cysteine-rich RLK (RECEPTOR-like protein kinase) 25 | |
| 196 | | 34 | 21 | 34 | 29 | 24 | 22 | 26 | 18 | 27 | 18 | 16 | 18 | 13 | 18 | 0.0060 | FvH4_3g027 | | 138404 | 138684 | | |
| 46 | GENE19542 | 9 | 8 | 2 | 5 | 0 | 7 | 1 | 0 | 7 | 8 | 0 | 5 | 2 | 9 | 2831 | 30 | Fvb3 | 3 | 9 | Trimeric LpxA-like enzymes superfamily protein | |
| 196 | | 27. | 33. | 31. | | 29. | 39. | 38. | 90. | 33. | 29. | 33. | 42. | 29. | 25. | 0.4648 | FvH4_3g027 | | 138348 | 138406 | Late embryogenesis abundant (LEA) hydroxyproline-rich | |
| 47 | GENE19543 | 9 | 8 | 6 | 36 | 2 | 9 | 6 | 3 | 5 | 8 | 8 | 3 | 4 | 1 | 4317 | 20 | Fvb3 | 0 | 4 | glycoprotein family | |

| | | | | | | | | | | | | | | | | | | | | | | |
|---|---|---|---|---|---|---|---|---|---|---|---|---|---|---|---|---|---|---|---|---|---|---|
| 196 | | 54. | 35. | 45. | 52. | 34. | 62. | 58. | 74. | 45. | 43. | 54. | 55. | 32. | 43. | 0.9057 | FvH4_3g027 | | 137821 | 138013 | | |
| 48 | GENE19544 | 8 | 1 | 7 | 2 | 1 | 8 | 3 | 3 | 4 | 2 | 2 | 3 | 9 | 3 | 6558 | 00 | Fvb3 | 1 | 0 | Peroxidase superfamily protein | |
| 196 | | ## | 84 | 51 | 42 | 64 | 51 | 44 | 71 | 44 | 54 | 54 | 49 | 43 | 53 | 0.2966 | FvH4_3g026 | | 136855 | 137155 | | |
| 49 | GENE19545 | ## | 01 | 74 | 05 | 29 | 47 | 94 | 54 | 73 | 46 | 67 | 45 | 04 | 28 | 7534 | 90 | Fvb3 | 2 | 2 | | 0 |
| 196 | | 19 | 31 | 24 | 25 | 27 | 30 | 27 | 36 | 32 | 33 | 26 | 38 | 25 | 30 | 0.0409 | FvH4_3g026 | | 136562 | 136745 | | |
| 50 | GENE19546 | 2 | 2 | 5 | 5 | 0 | 3 | 7 | 9 | 0 | 8 | 1 | 8 | 9 | 7 | 2827 | 80 | Fvb3 | 1 | 9 | PPDK regulatory protein | |
| 196 | | 32. | 31. | 28. | 27. | | | 28. | 30. | 28. | 29. | 29. | | 27. | | 0.9579 | FvH4_3g026 | | 136038 | 136075 | | |
| 51 | GENE19547 | 5 | 7 | 9 | 7 | 27 | 28 | 6 | 1 | 8 | 7 | 6 | 29 | 8 | 29 | 7146 | 71 | Fvb3 | 8 | 7 | | 0 |
| 196 | | 25 | 28 | 31 | 35 | 25 | 42 | 38 | 34 | 52 | 42 | 35 | 36 | 34 | 50 | 0.0525 | FvH4_3g026 | | 134095 | 134586 | | |
| 54 | GENE19550 | 9 | 5 | 9 | 6 | 9 | 2 | 3 | 2 | 1 | 2 | 6 | 2 | 4 | 8 | 5348 | 50 | Fvb3 | 8 | 8 | tyrosyl-DNA phosphodiesterase-related | |
| 196 | | 11 | 97 | 10 | 96 | 90 | 11 | 94 | 11 | 10 | 86 | 10 | 87 | 85 | 10 | 0.5019 | FvH4_3g026 | | 133701 | 134076 | | |
| 55 | GENE19551 | 41 | 0 | 17 | 1 | 4 | 24 | 7 | 09 | 70 | 7 | 17 | 9 | 0 | 17 | 4054 | 41 | Fvb3 | 9 | 0 | polyubiquitin 10 | |
| 196 | | 75 | 28 | 31 | 38 | 30 | 30 | 23 | 22 | 26 | 22 | 19 | 20 | 33 | 16 | 0.0847 | FvH4_3g026 | | 133477 | 133632 | | |
| 56 | GENE19552 | 0 | 6 | 9 | 3 | 7 | 0 | 4 | 1 | 9 | 2 | 2 | 8 | 6 | 8 | 9406 | 40 | Fvb3 | 1 | 9 | postsynaptic protein-related | |
| 196 | | 18 | 15 | 23 | 16 | 88 | 44 | 12 | 39 | 69. | 14 | 17 | 20 | 17 | 10 | 0.2705 | FvH4_3g026 | | 133173 | 133385 | | |
| 57 | GENE19553 | 2 | 0 | 8 | 7 | 0 | 6 | 9 | 8 | 5 | 9 | 8 | 7 | 0 | 2 | 672 | 30 | Fvb3 | 2 | 4 | xyloglucan endotransglucosylase/hydrolase 8 | |
| 196 | | 63. | 26. | 27. | 28. | 44. | | 48. | 36. | 46. | | 29. | 27. | 34. | 19. | 0.2923 | FvH4_3g026 | | 132663 | 132795 | | |
| 59 | GENE19555 | 7 | 2 | 8 | 4 | 7 | 49 | 7 | 1 | 6 | 45 | 5 | 1 | 3 | 8 | 8918 | 20 | Fvb3 | 8 | 7 | Tetratricopeptide repeat (TPR)-like superfamily protein | |
| 196 | | ## | ## | ## | ## | ## | ## | ## | ## | ## | ## | ## | ## | ## | ## | 0.0077 | FvH4_3g026 | | 132497 | 132599 | Oligosaccharyltransferase complex/magnesium transporter | |
| 60 | GENE19556 | ## | ## | ## | ## | ## | ## | ## | ## | ## | ## | ## | ## | ## | ## | 0245 | 00 | Fvb3 | 9 | 8 | family protein | |
| 196 | | 11 | 11 | 13 | 12 | 12 | 11 | 13 | 10 | 12 | 13 | 12 | 12 | 12 | 96. | 0.4555 | FvH4_3g025 | | 132266 | 132385 | | |
| 61 | GENE19557 | 7 | 4 | 3 | 8 | 2 | 1 | 9 | 9 | 0 | 2 | 3 | 7 | 4 | 2 | 7236 | 94 | Fvb3 | 3 | 3 | F-box family protein | |
| 196 | | 25. | 21. | 22. | 29. | 28. | 27. | 24. | 36. | 30. | 26. | 28. | 28. | 25. | 27. | 0.0727 | FvH4_3g025 | | 131822 | 131921 | | |
| 62 | GENE19558 | 7 | 9 | 9 | 1 | 6 | 2 | 3 | 4 | 1 | 7 | 3 | 8 | 6 | 4 | 4115 | 93 | Fvb3 | 9 | 3 | | 0 |
| 196 | | 10. | 8.8 | 9.8 | 8.3 | 9.8 | 8.9 | 8.7 | 9.7 | 10. | 8.9 | 9.4 | 10. | 8.1 | 9.5 | 0.6019 | FvH4_3g025 | | 131299 | 131539 | | |
| 63 | GENE19559 | 1 | 9 | 7 | 3 | 5 | 6 | 3 | 9 | 3 | 2 | 1 | 1 | 3 | 4 | 1827 | 91 | Fvb3 | 3 | 4 | F-box/RNI-like superfamily protein | |
| 196 | | 41. | 36. | 37. | 51. | 50. | 42. | 40. | 45. | 45. | | 42. | 49. | 42. | 44. | 0.5327 | FvH4_3g025 | | 130782 | 130917 | | |
| 64 | GENE19560 | 4 | 5 | 4 | 5 | 4 | 9 | 5 | 7 | 3 | 42 | 8 | 5 | 4 | 1 | 8911 | 90 | Fvb3 | 1 | 4 | | 0 |
| 196 | | 11 | 17 | 20 | 12 | 19 | 12 | 20 | 13 | 22 | 18 | 26 | 16 | 21 | 26 | 0.0914 | FvH4_3g025 | | 130490 | 130537 | | |
| 65 | GENE19561 | 4 | 2 | 2 | 9 | 7 | 9 | 7 | 5 | 5 | 4 | 7 | 5 | 7 | 6 | 1769 | 81 | Fvb3 | 3 | 0 | 1-amino-cyclopropane-1-carboxylate synthase 12 | |
| 196 | | 15. | 11. | | 12. | 13. | 11. | 11. | 11. | | 13. | 12. | 13. | 13. | 13. | 0.5232 | FvH4_3g025 | | 130141 | 130351 | | |
| 66 | GENE19562 | 6 | 5 | 11 | 6 | 4 | 5 | 5 | 1 | 14 | 1 | 3 | 1 | 6 | 1 | 7301 | 80 | Fvb3 | 8 | 4 | F-box/RNI-like superfamily protein | |
| 196 | | 15 | 18 | 23 | 22 | 16 | 17 | 27 | 32 | 31 | 23 | 38 | 30 | 25 | 46 | 0.0046 | FvH4_3g025 | | 129813 | 129844 | | |
| 67 | GENE19563 | 1 | 0 | 6 | 9 | 1 | 9 | 3 | 6 | 1 | 9 | 7 | 6 | 0 | 2 | 0349 | 72 | Fvb3 | 2 | 6 | ferrochelatase 2 | |
| 196 | | 21. | 24. | 23. | 30. | 27. | 27. | 21. | 25. | 27. | 24. | 28. | 26. | 23. | 23. | 0.7607 | FvH4_3g025 | | 129344 | 129492 | | |
| 68 | GENE19564 | 5 | 7 | 2 | 5 | 2 | 1 | 6 | 9 | 1 | 5 | 1 | 4 | 5 | 5 | 9662 | 71 | Fvb3 | 4 | 6 | | 0 |
| 196 | | 17 | 91 | 61 | 61 | 63 | 10 | 80 | 53 | 46 | 77 | 40 | 46 | 78 | 51 | 0.0653 | FvH4_3g025 | | 128920 | 129263 | | |
| 70 | GENE19566 | 34 | 1 | 8 | 2 | 3 | 51 | 7 | 1 | 5 | 0 | 9 | 1 | 2 | 5 | 0566 | 70 | Fvb3 | 1 | 3 | PAK-box/P21-Rho-binding family protein | |
| 196 | | 13 | 93. | 63. | 31. | 11 | 16 | 18 | 10 | 41. | 63. | 54. | 82. | 86. | 37. | 0.0803 | FvH4_3g025 | | 128435 | 128659 | | |
| 71 | GENE19567 | 4 | 8 | 4 | 5 | 1 | 4 | 5 | 3 | 1 | 8 | 9 | 2 | 1 | 9 | 6886 | 60 | Fvb3 | 7 | 7 | methylesterase PCR A | |
| 196 | | 13 | 27 | 27 | 21 | 35 | 28 | 12 | 54 | 86. | 13 | 18 | 28 | 17 | 65. | 0.6907 | FvH4_3g025 | | 128288 | 128382 | | |
| 72 | GENE19568 | 7 | 6 | 4 | 8 | 3 | 9 | 3 | 5 | 3 | 9 | 1 | 1 | 1 | 4 | 177 | 50 | Fvb3 | 2 | 7 | NTF2-like | |
| 196 | | 18. | 20. | 17. | 18. | 17. | | 19. | 17. | 16. | 19. | 24. | 18. | 19. | 22. | 0.2907 | FvH4_3g025 | | 127970 | 128217 | | |
| 74 | GENE19570 | 5 | 4 | 4 | 8 | 4 | 17 | 4 | 5 | 5 | 3 | 1 | 2 | 8 | 5 | 7121 | 30 | Fvb3 | 1 | 0 | COBRA-like protein 11 precursor | |
| 196 | | 59 | 87 | 90 | 98 | 62 | 11 | 55 | 96 | 89 | 93 | 85 | 57 | 71 | 10 | 0.7504 | FvH4_3g025 | | 126796 | 127344 | | |
| 76 | GENE19572 | 2 | 4 | 8 | 8 | 9 | 50 | 7 | 7 | 5 | 3 | 5 | 0 | 4 | 01 | 3903 | 20 | Fvb3 | 0 | 3 | protodermal factor 2 | |

| | | | | | | | | | | | | | | | | | | | | | | |
|---|---|---|---|---|---|---|---|---|---|---|---|---|---|---|---|---|---|---|---|---|---|---|
| 196 | | 20 | 29 | 23 | 15 | 30 | 31 | 33 | 33 | 32 | 31 | 26 | 27 | 32 | 44 | 0.0825 | FvH4_3g025 | | 125597 | 126207 | | |
| 77 | GENE19573 | 06 | 41 | 29 | 16 | 94 | 04 | 36 | 33 | 32 | 56 | 76 | 35 | 39 | 66 | 2145 | 10 | Fvb3 | 1 | 6 | PCF11P-similar protein 4 | |
| 196 | | 13 | 13 | 15 | 10 | 13 | 98. | 16 | 62. | | 64. | 91. | 60. | 58. | 56. | 0.0001 | FvH4_3g024 | | 123767 | 124264 | | |
| 81 | GENE19577 | 2 | 4 | 9 | 6 | 5 | 6 | 7 | 7 | 93 | 3 | 2 | 5 | 2 | 5 | 9409 | 80 | Fvb3 | 4 | 2 | Pseudouridine synthase family protein | |
| 196 | | 31 | 27 | 42 | 48 | 42 | 32 | 33 | 31 | 30 | 33 | 41 | 49 | 26 | 31 | 0.6141 | FvH4_3g024 | | 123509 | 123813 | | |
| 82 | GENE19578 | 6 | 4 | 3 | 3 | 4 | 9 | 6 | 1 | 4 | 8 | 4 | 5 | 1 | 0 | 7767 | 90 | Fvb3 | 8 | 0 | adenosine/AMP deaminase family protein | |
| 196 | | 16. | 17. | | | 18. | 18. | 18. | 21. | 19. | 16. | 19. | | 19. | 22. | 0.1051 | FvH4_3g024 | | 123380 | 123442 | | |
| 83 | GENE19579 | 8 | 1 | 18 | 17 | 7 | 5 | 7 | 3 | 4 | 7 | 4 | 17 | 6 | 9 | 343 | 70 | Fvb3 | 7 | 4 | LA RNA-binding protein | |
| 196 | | 13 | 12 | 81 | 69 | 11 | 16 | 12 | 10 | 15 | 13 | 12 | 11 | 11 | 10 | 0.7364 | FvH4_3g024 | | 122536 | 123245 | | |
| 84 | GENE19580 | 61 | 02 | 6 | 8 | 69 | 51 | 77 | 95 | 32 | 40 | 05 | 45 | 62 | 30 | 2559 | 60 | Fvb3 | 0 | 7 | phospholipases;galactolipases | |
| 196 | | 18. | 22. | 22. | 24. | 27. | 28. | 22. | 24. | 22. | | 30. | 50. | 18. | 17. | 0.5127 | FvH4_3g024 | | 122386 | 122426 | | |
| 85 | GENE19581 | 6 | 7 | 6 | 4 | 3 | 9 | 7 | 9 | 3 | 24 | 9 | 5 | 5 | 5 | 9267 | 50 | Fvb3 | 8 | 3 | | 0 |
| 196 | | 15. | 12. | 15. | 15. | 13. | 16. | 13. | 14. | | 15. | 13. | 14. | 13. | 15. | 0.7892 | FvH4_3g024 | | 122171 | 122425 | | |
| 86 | GENE19582 | 3 | 1 | 6 | 1 | 9 | 1 | 8 | 8 | 13 | 1 | 9 | 9 | 8 | 2 | 4359 | 40 | Fvb3 | 9 | 7 | Protein kinase superfamily protein | |
| 196 | | 16 | 18 | 14 | 11 | 16 | 13 | 19 | 15 | 20 | 11 | 16 | 17 | 10 | 19 | 0.9222 | FvH4_3g024 | | 121867 | 122095 | | |
| 87 | GENE19583 | 00 | 06 | 79 | 58 | 33 | 55 | 47 | 38 | 60 | 41 | 32 | 80 | 17 | 32 | 553 | 30 | Fvb3 | 6 | 5 | cardiolipin synthase | |
| 196 | | 64. | 41. | 80. | 62. | 81. | 77. | | 60. | 77. | 78. | 84. | 61. | 81. | | 0.2098 | FvH4_3g024 | | 121795 | 121859 | | |
| 88 | GENE19584 | 2 | 9 | 4 | 9 | 6 | 3 | 66 | 8 | 5 | 3 | 1 | 2 | 1 | 98 | 9138 | 21 | Fvb3 | 9 | 6 | | 0 |
| 247 | | 12 | 73. | 89. | 82. | 14 | 92. | 15 | 95. | 93. | 16 | 89. | 80. | 87. | 12 | 0.8846 | FvH4_3g028 | | 152811 | 152869 | | |
| 27 | GENE24648 | 1 | 8 | 4 | 1 | 1 | 2 | 0 | 5 | 9 | 1 | 2 | 4 | 5 | 7 | 5866 | 52 | Fvb3 | 5 | 2 | | 0 |
| 247 | | 14 | 56 | 79 | 18 | 41 | 21 | 11 | 21 | 56 | 72 | 47 | 29 | 91 | 32 | 0.0297 | FvH4_3g028 | | 153198 | 153489 | | |
| 28 | GENE24649 | 78 | 2 | 4 | 46 | 8 | 28 | 90 | 9 | 9 | 2 | 2 | 6 | 0 | 8 | 9006 | 70 | Fvb3 | 0 | 7 | cysteine-rich RLK (RECEPTOR-like protein kinase) 29 | |
| 247 | | 15. | 11. | 15. | 12. | 14. | 14. | 12. | 16. | 13. | 16. | 12. | 13. | 12. | 13. | 0.8511 | FvH4_3g028 | | 153697 | 153761 | | |
| 29 | GENE24650 | 5 | 8 | 7 | 8 | 4 | 2 | 6 | 4 | 3 | 4 | 6 | 6 | 6 | 2 | 6486 | 80 | Fvb3 | 1 | 0 | AGAMOUS-like 29 | |
| 247 | | 13. | 19. | 11. | | 19. | | 14. | 19. | 25. | 14. | 16. | 14. | 11. | 18. | 0.4058 | FvH4_3g028 | | 153770 | 153981 | | |
| 30 | GENE24651 | 7 | 7 | 6 | 11 | 1 | 17 | 9 | 4 | 7 | 1 | 6 | 7 | 2 | 7 | 97 | 81 | Fvb3 | 9 | 5 | | 0 |
| 247 | | 54. | 43. | 29. | 40. | 41. | | 35. | 63. | 43. | 39. | 41. | 51. | 36. | 39. | 0.8563 | FvH4_3g028 | | 154297 | 154524 | | |
| 31 | GENE24652 | 1 | 7 | 5 | 9 | 4 | 63 | 2 | 5 | 5 | 5 | 1 | 4 | 9 | 2 | 0989 | 90 | Fvb3 | 8 | 7 | RNI-like superfamily protein | |
| 247 | | 28 | 77 | 53 | 14 | 23 | 49 | 13 | 51 | 10 | 17 | 76 | 98 | 71 | 58 | 0.1616 | FvH4_3g029 | | 154673 | 154967 | | |
| 32 | GENE24653 | 9 | 0 | 6 | 75 | 2 | 3 | 8 | 8 | 79 | 78 | 1 | 0 | 0 | 0 | 0935 | 00 | Fvb3 | 1 | 8 | Sulfite exporter TauE/SafE family protein | |
| 247 | | 77 | 45 | 54 | 48 | 46 | 52 | 50 | 40 | 45 | 46 | 51 | 49 | 39 | 47 | 0.1154 | FvH4_3g029 | | 154970 | 155247 | | |
| 33 | GENE24654 | 02 | 53 | 96 | 95 | 30 | 14 | 09 | 15 | 71 | 23 | 55 | 99 | 28 | 15 | 2423 | 10 | Fvb3 | 1 | 7 | basic helix-loop-helix (bHLH) DNA-binding superfamily protein | |
| 247 | | 15 | 11 | 22 | 13 | 23 | 10 | 13 | 14 | 46 | 62 | 16 | 46 | 35 | 23 | 0.5607 | FvH4_3g029 | | 155328 | 155466 | | |
| 34 | GENE24655 | 81 | 12 | 20 | 06 | 2 | 46 | 82 | 03 | 9 | 0 | 13 | 8 | 7 | 96 | 2576 | 11 | Fvb3 | 1 | 8 | C-terminal cysteine residue is changed to a serine 1 | |
| 247 | | 32 | 46 | 22 | 47 | 45 | 40 | 57 | 31 | 29 | 18 | 20 | 20 | 18 | 17 | 0.0030 | FvH4_3g029 | | 155483 | 155982 | | |
| 35 | GENE24656 | 9 | 9 | 2 | 7 | 3 | 4 | 7 | 4 | 4 | 2 | 5 | 6 | 7 | 1 | 5414 | 12 | Fvb3 | 3 | 4 | kinesin 1 | |
| 247 | | 44. | 15 | 20 | 81. | 53. | 14 | 71. | 32 | 69. | 31 | 14 | 39 | 66. | 60. | 0.1658 | FvH4_3g029 | | 156143 | 156290 | | |
| 36 | GENE24657 | 8 | 5 | 5 | 5 | 4 | 0 | 3 | 7 | 5 | 8 | 5 | 9 | 4 | 9 | 1222 | 20 | Fvb3 | 8 | 4 | photosystem II subunit Q-2 | |
| 247 | | ## | ## | ## | ## | ## | ## | ## | ## | ## | ## | ## | ## | ## | ## | 0.2487 | FvH4_3g029 | | 156414 | 156705 | DNA-binding storekeeper protein-related transcriptional | |
| 37 | GENE24658 | ## | ## | ## | ## | ## | ## | ## | ## | ## | ## | ## | ## | ## | ## | 9509 | 30 | Fvb3 | 6 | 6 | regulator | |
| 247 | | 65 | 51 | 99 | 96 | 53 | 58 | 48 | 32 | 34 | 26 | 43 | 48 | 60 | 30 | 0.0141 | FvH4_3g029 | | 156912 | 157013 | | |
| 38 | GENE24659 | 0 | 1 | 3 | 3 | 2 | 4 | 5 | 2 | 3 | 5 | 7 | 7 | 0 | 8 | 0432 | 40 | Fvb3 | 1 | 0 | histone H2A 2 | |
| 247 | | 38. | 99. | 75. | 63. | 52. | 62. | 43. | 16 | 50. | 58. | 71. | 18 | 61. | 44. | 0.2547 | FvH4_3g029 | | 156987 | 157081 | | |
| 39 | GENE24660 | 4 | 3 | 1 | 2 | 8 | 2 | 9 | 8 | 4 | 3 | 3 | 2 | 4 | 8 | 8355 | 50 | Fvb3 | 8 | 6 | Protein of unknown function (DUF1218) | |
| 247 | | ## | ## | ## | ## | ## | ## | ## | ## | ## | ## | ## | ## | ## | ## | 0.1197 | FvH4_3g029 | | 157114 | 157391 | | |
| 40 | GENE24661 | ## | ## | ## | ## | ## | ## | ## | ## | ## | ## | ## | ## | ## | ## | 1601 | 60 | Fvb3 | 6 | 2 | Aldolase-type TIM barrel family protein | |

| | | | | | | | | | | | | | | | | | | | | | | |
|---|---|---|---|---|---|---|---|---|---|---|---|---|---|---|---|---|---|---|---|---|---|---|
| 247 | | 18 | 16 | 13 | 15 | 13 | 16 | 16 | 17 | 16 | 15 | 19 | 14 | 18 | 19 | 0.1410 | FvH4_3g029 | | 157600 | 157824 | | |
| 42 | GENE24663 | 56 | 17 | 43 | 88 | 71 | 99 | 94 | 98 | 56 | 84 | 38 | 78 | 22 | 84 | 4981 | 70 | Fvb3 | 6 | 8 | | 0 |
| 247 | | 27. | 35. | 27. | 29. | 29. | 42. | 48. | 25. | 33. | 34. | 36. | 27. | 28. | 60. | 0.8905 | FvH4_3g029 | | 157884 | 158226 | | |
| 43 | GENE24664 | 3 | 4 | 9 | 4 | 6 | 2 | 3 | 6 | 2 | 3 | 1 | 2 | 2 | 8 | 3707 | 71 | Fvb3 | 6 | 5 | | 0 |
| 247 | | 96 | 16 | 21 | 19 | 98 | 16 | 12 | 19 | 43 | 39 | 28 | 27 | 27 | 20 | 0.0048 | FvH4_3g029 | | 158375 | 158681 | | |
| 44 | GENE24665 | 6 | 07 | 68 | 68 | 8 | 93 | 45 | 04 | 07 | 52 | 26 | 79 | 41 | 48 | 7185 | 80 | Fvb3 | 5 | 9 | NAD(P)-binding Rossmann-fold superfamily protein | |
| 247 | | 26. | 23. | 24. | 22. | | 26. | 27. | 25. | | 24. | 23. | 21. | | | 0.8550 | FvH4_3g029 | | 158547 | 158882 | | |
| 45 | GENE24666 | 5 | 3 | 9 | 7 | 24 | 8 | 2 | 5 | 32 | 4 | 9 | 6 | 27 | 23 | 3542 | 90 | Fvb3 | 4 | 1 | Protein kinase superfamily protein | |
| 247 | | 48 | 94 | 23 | 33 | 18 | 17 | 18 | 71 | 20 | 56 | 23 | 36 | 60 | 24 | 0.7931 | FvH4_3g030 | | 159074 | 159354 | | |
| 46 | GENE24667 | 42 | 1 | 08 | 64 | 41 | 00 | 42 | 2 | 4 | 35 | 77 | 50 | 14 | 5 | 2829 | 00 | Fvb3 | 7 | 2 | terpene synthase 03 | |
| 247 | | 72 | 10 | 11 | 82 | 10 | 91 | 14 | 14 | 94 | 10 | 12 | 12 | 10 | 15 | 0.1476 | FvH4_3g030 | | 159460 | 159570 | | |
| 47 | GENE24668 | 8 | 93 | 37 | 4 | 20 | 2 | 58 | 05 | 6 | 59 | 46 | 52 | 42 | 27 | 5084 | 10 | Fvb3 | 3 | 9 | RING/U-box superfamily protein | |
| 247 | | 49 | 61 | 50 | 41 | 55 | 60 | 67 | 72 | 67 | 61 | 71 | 56 | 63 | 79 | 0.0179 | FvH4_3g030 | | 159966 | 160424 | | |
| 48 | GENE24669 | 51 | 41 | 86 | 19 | 11 | 47 | 83 | 91 | 63 | 94 | 10 | 10 | 12 | 58 | 9402 | 30 | Fvb3 | 6 | 4 | eukaryotic translation initiation factor-related | |
| 247 | | 25. | 20. | 21. | 23. | 18. | 24. | 21. | | 26. | 24. | 25. | 22. | 18. | 19. | 0.5898 | FvH4_3g030 | | 160451 | 160734 | | |
| 49 | GENE24670 | 1 | 3 | 3 | 1 | 7 | 8 | 1 | 23 | 5 | 6 | 5 | 2 | 8 | 3 | 4169 | 40 | Fvb3 | 7 | 9 | Phototropic-responsive NPH3 family protein | |
| 247 | | 14 | 25 | 17 | 32 | 23 | 30 | 24 | 25 | 28 | 37 | 20 | 22 | 26 | 19 | 0.5904 | FvH4_3g030 | | 160788 | 161277 | | |
| 50 | GENE24671 | 05 | 44 | 99 | 75 | 96 | 10 | 15 | 63 | 64 | 64 | 93 | 37 | 43 | 78 | 729 | 41 | Fvb3 | 1 | 1 | phosphoinositide binding | |
| 247 | | 57. | 35. | 40. | 41. | 35. | 36. | 29. | 29. | 29. | 44. | 29. | 40. | 44. | 37. | 0.5057 | FvH4_3g030 | | 161329 | 161571 | | |
| 51 | GENE24672 | 8 | 4 | 1 | 3 | 3 | 1 | 8 | 1 | 2 | 4 | 9 | 4 | 2 | 9 | 6999 | 42 | Fvb3 | 0 | 0 | terpene synthase 14 | |
| 247 | | 13. | 13. | 15. | | 16. | 11. | 13. | 12. | 15. | 15. | 16. | 13. | | 17. | 0.5814 | FvH4_3g030 | | 164511 | 164752 | | |
| 52 | GENE24673 | 4 | 4 | 8 | 14 | 3 | 8 | 7 | 2 | 1 | 3 | 4 | 8 | 12 | 5 | 2249 | 80 | Fvb3 | 9 | 7 | Tetratricopeptide repeat (TPR)-like superfamily protein | |
| 247 | | 22 | 24 | 33 | 36 | 32 | 32 | 36 | 27 | 51 | 32 | 48 | 38 | 33 | 50 | 0.0557 | FvH4_3g030 | | 162927 | 162992 | | |
| 54 | GENE24675 | 89 | 62 | 47 | 21 | 60 | 46 | 75 | 44 | 09 | 97 | 46 | 97 | 33 | 08 | 4981 | 60 | Fvb3 | 0 | 4 | Tetratricopeptide repeat (TPR)-like superfamily protein | |
| 247 | | ## | ## | ## | ## | ## | 98 | ## | ## | ## | ## | ## | ## | ## | ## | 0.6881 | FvH4_3g030 | | 163320 | 163536 | | |
| 55 | GENE24676 | ## | ## | ## | ## | ## | 93 | ## | ## | ## | ## | ## | ## | ## | ## | 6534 | 61 | Fvb3 | 6 | 2 | terpene synthase 14 | |
| 247 | | 33 | 33 | 34 | 29 | 28 | 22 | 34 | 27 | 37 | 24 | 43 | 30 | 28 | 48 | 0.3827 | FvH4_3g030 | | 163567 | 163631 | | |
| 56 | GENE24677 | 48 | 58 | 49 | 91 | 03 | 92 | 55 | 62 | 84 | 66 | 77 | 76 | 29 | 29 | 1562 | 70 | Fvb3 | 0 | 6 | Tetratricopeptide repeat (TPR)-like superfamily protein | |
| 247 | | 33. | 28. | 82. | 98. | | | 10 | 21. | 10 | 10 | | 66. | 80. | 84. | 0.9505 | FvH4_3g030 | | 163670 | 163789 | | |
| 57 | GENE24678 | 1 | 4 | 2 | 6 | 97 | 94 | 5 | 2 | 1 | 2 | 77 | 2 | 8 | 3 | 3718 | 71 | Fvb3 | 6 | 2 | phosphoinositide binding | |
| 304 | | | 18. | | 20. | | | 22. | 21. | 22. | | 19. | 18. | 20. | 21. | 0.1997 | FvH4_3g046 | | 266773 | 266838 | | |
| 91 | GENE30437 | 25 | 7 | 23 | 2 | 24 | 21 | 1 | 8 | 6 | 20 | 5 | 2 | 7 | 6 | 2271 | 41 | Fvb3 | 3 | 0 | | 0 |
| 304 | | 24. | 22. | 28. | 26. | 25. | 30. | 24. | 30. | 28. | 24. | | 22. | 20. | 27. | 0.8388 | FvH4_3g046 | | 266256 | 266336 | | |
| 92 | GENE30438 | 5 | 9 | 6 | 2 | 9 | 4 | 3 | 1 | 8 | 9 | 25 | 8 | 8 | 9 | 4801 | 40 | Fvb3 | 0 | 8 | Toll-Interleukin-Resistance (TIR) domain family protein | |
| 304 | | ## | 93 | ## | ## | 93 | 89 | 96 | 84 | 85 | 99 | 89 | 82 | ## | ## | 0.2419 | FvH4_3g046 | | 265564 | 265742 | | |
| 93 | GENE30439 | ## | 71 | ## | ## | 58 | 08 | 24 | 11 | 66 | 08 | 13 | 02 | ## | ## | 7049 | 30 | Fvb3 | 3 | 3 | NAC domain containing protein 25 | |
| 304 | | 10. | | 15. | 14. | 13. | 14. | 13. | 14. | 16. | 16. | 17. | 14. | 14. | 12. | 0.2013 | FvH4_3g046 | | 265130 | 265413 | | |
| 94 | GENE30440 | 8 | 14 | 8 | 7 | 6 | 5 | 7 | 1 | 2 | 6 | 7 | 3 | 5 | 4 | 9644 | 20 | Fvb3 | 5 | 9 | serine carboxypeptidase-like 31 | |
| 304 | | | 32. | 33. | 26. | 34. | 32. | 38. | 24. | 39. | 40. | 34. | 62. | 53. | 37. | 0.0922 | FvH4_3g046 | | 264567 | 264964 | | |
| 95 | GENE30441 | 28 | 6 | 2 | 5 | 3 | 4 | 6 | 8 | 3 | 6 | 7 | 8 | 4 | 6 | 5467 | 11 | Fvb3 | 3 | 4 | | 0 |
| 304 | | 18. | | 18. | 18. | 19. | 20. | 20. | 19. | 16. | 17. | 19. | 24. | 18. | 19. | 0.9176 | FvH4_3g046 | | 263800 | 264048 | | |
| 96 | GENE30442 | 2 | 21 | 9 | 2 | 5 | 9 | 2 | 9 | 6 | 6 | 4 | 3 | 9 | 3 | 1351 | 10 | Fvb3 | 3 | 1 | cytokinin oxidase 7 | |
| 304 | | 30. | 54. | 28. | 29. | 28. | 35. | 55. | 32. | 38. | 30. | 45. | 61. | 27. | 32. | 0.8967 | FvH4_3g046 | | 263358 | 263746 | | |
| 97 | GENE30443 | 3 | 8 | 8 | 3 | 6 | 6 | 5 | 5 | 5 | 1 | 9 | 3 | 9 | 6 | 993 | 00 | Fvb3 | 5 | 7 | serine carboxypeptidase-like 31 | |
| 304 | | 12 | 13 | 14 | 10 | 16 | 16 | 19 | 13 | 15 | 15 | 15 | 16 | 17 | 12 | 0.8464 | FvH4_3g045 | | 262718 | 262814 | | |
| 98 | GENE30444 | 52 | 18 | 95 | 34 | 20 | 92 | 68 | 79 | 01 | 22 | 08 | 81 | 10 | 60 | 4335 | 90 | Fvb3 | 9 | 8 | alpha/beta-Hydrolases superfamily protein | |

| | | | | | | | | | | | | | | | | | | | | | | |
|---|---|---|---|---|---|---|---|---|---|---|---|---|---|---|---|---|---|---|---|---|---|---|
| 304 | | 13 | 30 | 20 | 22 | 28 | 32 | 29 | 45 | 35 | 34 | 25 | 20 | 33 | 40 | 0.0676 | FvH4_3g045 | | 261871 | 263180 | SNF2 domain-containing protein / helicase domain-containing | |
| 99 | GENE30445 | 83 | 07 | 20 | 11 | 09 | 84 | 84 | 78 | 56 | 06 | 87 | 20 | 44 | 28 | 1169 | 80 | Fvb3 | 8 | 7 | protein / zinc finger protein-related | |
| 305 | | 73 | 56 | 65 | 64 | 60 | 52 | 56 | 59 | 50 | 52 | 57 | 57 | 49 | 53 | 0.0525 | FvH4_3g045 | | 261354 | 261682 | | |
| 00 | GENE30446 | 57 | 00 | 36 | 17 | 32 | 96 | 13 | 92 | 01 | 34 | 80 | 57 | 76 | 36 | 2716 | 70 | Fvb3 | 7 | 7 | LisH and RanBPM domains containing protein | |
| 305 | | 24. | 22. | 33. | 87. | 32. | | 19. | 72. | 18. | 23. | 58. | 43. | 23. | 28. | 0.6579 | FvH4_3g045 | | 261247 | 261337 | | |
| 01 | GENE30447 | 4 | 5 | 1 | 5 | 8 | 93 | 3 | 2 | 8 | 2 | 5 | 3 | 3 | 1 | 3814 | 60 | Fvb3 | 3 | 2 | | 0 |
| 305 | | 18 | 15 | 41 | 22 | 31 | 32 | 44 | 39 | 24 | 47 | 50 | 29 | 46 | 25 | 0.1888 | FvH4_3g045 | | 260580 | 260802 | | |
| 02 | GENE30448 | 2 | 3 | 3 | 6 | 7 | 1 | 0 | 7 | 6 | 8 | 3 | 3 | 4 | 0 | 4472 | 50 | Fvb3 | 9 | 9 | lysine decarboxylase family protein | |
| 305 | | 39. | 45. | 84. | 61. | 67. | 58. | 62. | 43. | 76. | 73. | 67. | 77. | | 89. | 0.1347 | FvH4_3g045 | | 260307 | 260350 | | |
| 03 | GENE30449 | 3 | 3 | 5 | 1 | 3 | 2 | 6 | 2 | 8 | 5 | 7 | 8 | 77 | 6 | 4892 | 41 | Fvb3 | 1 | 4 | | 0 |
| 305 | | 61. | 51. | 79. | 69. | | 86. | 52. | 70. | | 54. | 93. | 15 | 53. | 47. | 0.8156 | FvH4_3g045 | | 259749 | 259992 | | |
| 04 | GENE30450 | 7 | 7 | 1 | 9 | 81 | 9 | 7 | 6 | 39 | 1 | 7 | 1 | 4 | 9 | 3746 | 40 | Fvb3 | 9 | 4 | | 0 |
| 305 | | 10 | 90. | | 62. | 66. | 82. | 90. | | 23. | 20. | 24. | 24. | | 31. | 6.5343 | FvH4_3g045 | | 259506 | 259622 | | |
| 05 | GENE30451 | 8 | 2 | 64 | 1 | 8 | 6 | 3 | 40 | 4 | 6 | 3 | 1 | 41 | 1 | E-05 | 30 | Fvb3 | 1 | 1 | | 0 |
| 305 | | 38. | 30. | 35. | | 39. | 50. | 35. | 33. | 30. | 29. | 25. | | 40. | 27. | 0.0161 | FvH4_3g045 | | 259148 | 259389 | | |
| 06 | GENE30452 | 7 | 2 | 5 | 47 | 6 | 6 | 8 | 7 | 5 | 7 | 2 | 24 | 2 | 2 | 2379 | 21 | Fvb3 | 7 | 2 | BED zinc finger ;hAT family dimerisation domain | |
| 305 | | 71. | 63. | 65. | 56. | 44. | 56. | 50. | 13 | 46. | 38. | 55. | 41. | 43. | 62. | 0.8536 | FvH4_3g045 | | 258711 | 258892 | | |
| 07 | GENE30453 | 4 | 9 | 9 | 7 | 1 | 6 | 2 | 9 | 1 | 9 | 2 | 7 | 3 | 8 | 4529 | 20 | Fvb3 | 7 | 1 | | 0 |
| 305 | | 49 | 23 | 34 | 27 | 59 | 48 | 69 | 31 | 60 | 55 | 72 | 30 | 24 | 49 | 0.8620 | FvH4_3g045 | | 258422 | 258501 | | |
| 08 | GENE30454 | 0 | 1 | 4 | 0 | 1 | 5 | 8 | 1 | 2 | 2 | 6 | 0 | 3 | 3 | 1113 | 10 | Fvb3 | 0 | 3 | | 0 |
| 305 | | 10 | 16 | 15 | 18 | 21 | 15 | 11 | 24 | 26 | 21 | 23 | 19 | 11 | 42 | 0.0515 | FvH4_3g044 | | 256119 | 256425 | | |
| 10 | GENE30456 | 1 | 7 | 9 | 8 | 1 | 2 | 2 | 8 | 4 | 9 | 3 | 4 | 6 | 0 | 9352 | 70 | Fvb3 | 9 | 7 | bromodomain 4 | |
| 305 | | 49 | 41 | 51 | 52 | 41 | 57 | 43 | 17 | 44 | 40 | 17 | 27 | 56 | 20 | 0.0347 | FvH4_3g044 | | 255366 | 255865 | | |
| 11 | GENE30457 | 05 | 18 | 82 | 28 | 91 | 17 | 57 | 74 | 84 | 66 | 91 | 60 | 73 | 01 | 0987 | 60 | Fvb3 | 6 | 1 | Protein of unknown function (DUF707) | |
| 305 | | ## | ## | ## | ## | ## | ## | ## | ## | ## | ## | ## | ## | ## | ## | 0.5327 | FvH4_3g044 | | 254919 | 255312 | | |
| 12 | GENE30458 | ## | ## | ## | ## | ## | ## | ## | ## | ## | ## | ## | ## | ## | ## | 1328 | 40 | Fvb3 | 2 | 5 | secretory carrier 3 | |
| 305 | | 20 | 27 | 32 | 26 | 41 | 17 | 23 | 21 | 20 | 21 | 16 | 28 | 26 | 18 | 0.1642 | FvH4_3g044 | | 254813 | 254910 | | |
| 13 | GENE30459 | 01 | 82 | 50 | 18 | 16 | 94 | 36 | 80 | 92 | 67 | 04 | 14 | 06 | 57 | 9412 | 50 | Fvb3 | 6 | 1 | | 0 |
| 305 | | 79 | 57 | 66 | 58 | 60 | 62 | 57 | 63 | 47 | 54 | 48 | 45 | 53 | 43 | 0.0103 | FvH4_3g044 | | 254246 | 254572 | | |
| 14 | GENE30460 | 9 | 3 | 0 | 8 | 4 | 7 | 0 | 4 | 6 | 8 | 1 | 8 | 8 | 3 | 0202 | 30 | Fvb3 | 6 | 8 | HAL2-like | |
| 305 | | 60 | 12 | 60 | 10 | 16 | 13 | 15 | 12 | 18 | 15 | 85 | 96 | 21 | 13 | 0.2774 | FvH4_3g044 | | 252643 | 253992 | | |
| 16 | GENE30462 | 2 | 85 | 5 | 92 | 06 | 98 | 59 | 84 | 60 | 82 | 5 | 2 | 99 | 24 | 4171 | 10 | Fvb3 | 3 | 0 | HhH-GPD base excision DNA repair family protein | |
| 305 | | 8.9 | 8.1 | 6.4 | 6.2 | 5.9 | 6.7 | | 6.4 | 8.5 | 6.9 | | 8.2 | 7.4 | 7.8 | 0.4036 | FvH4_3g044 | | 251925 | 252167 | | |
| 17 | GENE30463 | 1 | 4 | 9 | 3 | 9 | 5 | 7.5 | 5 | 2 | 5 | 7.5 | 4 | 5 | 9 | 3095 | 01 | Fvb3 | 0 | 0 | | 0 |
| 305 | | 32. | 54. | 25. | 38. | 31. | 21. | 25. | 43. | | 20. | | 21. | 21. | 28. | 0.2935 | FvH4_3g044 | | 251638 | 251878 | | |
| 18 | GENE30464 | 1 | 3 | 3 | 2 | 4 | 5 | 1 | 8 | 21 | 2 | 31 | 4 | 4 | 1 | 5578 | 00 | Fvb3 | 5 | 1 | Homeodomain-like superfamily protein | |
| 305 | | 16 | 15 | 23 | 19 | 16 | 12 | 16 | 12 | 16 | 13 | 19 | 18 | 20 | 21 | 0.8442 | FvH4_3g043 | | 251028 | 251134 | | |
| 19 | GENE30465 | 5 | 6 | 9 | 3 | 3 | 3 | 2 | 5 | 6 | 9 | 8 | 0 | 3 | 7 | 4884 | 90 | Fvb3 | 3 | 0 | alpha/beta-Hydrolases superfamily protein | |
| 305 | | 56 | 48 | 68 | 48 | 75 | 41 | 26 | 46 | 88 | 92 | 10 | 89 | 43 | 78 | 0.0381 | FvH4_3g043 | | 250686 | 250862 | | |
| 20 | GENE30466 | 5 | 0 | 2 | 2 | 1 | 9 | 9 | 5 | 6 | 2 | 21 | 5 | 8 | 2 | 8937 | 81 | Fvb3 | 4 | 8 | | 0 |
| 305 | | 86 | 17 | 22 | 25 | 22 | 12 | 97 | 10 | 32 | 30 | 38 | 38 | 13 | 28 | 0.0587 | FvH4_3g043 | | 250237 | 250354 | | |
| 21 | GENE30467 | 5 | 43 | 07 | 53 | 60 | 45 | 5 | 99 | 58 | 71 | 38 | 55 | 16 | 14 | 314 | 80 | Fvb3 | 9 | 5 | | 0 |
| 305 | | | 27. | 32. | 36. | 28. | 29. | 34. | 22. | | 30. | 39. | | 35. | 37. | 0.5509 | FvH4_3g043 | | 250126 | 250173 | | |
| 22 | GENE30468 | 26 | 8 | 8 | 5 | 5 | 3 | 5 | 7 | 28 | 7 | 7 | 33 | 3 | 6 | 0544 | 72 | Fvb3 | 1 | 1 | | 0 |
| 305 | | | | 25. | 25. | 22. | 21. | | 27. | 27. | 27. | 26. | 26. | 23. | 25. | 0.0080 | FvH4_3g043 | | 249834 | 249862 | | |
| 23 | GENE30469 | 20 | 25 | 1 | 6 | 7 | 7 | 22 | 7 | 6 | 6 | 5 | 1 | 5 | 6 | 9538 | 71 | Fvb3 | 8 | 9 | | 0 |

| | | | | | | | | | | | | | | | | | | | | | | |
|---|---|---|---|---|---|---|---|---|---|---|---|---|---|---|---|---|---|---|---|---|---|---|
| 305 | | 28. | 20 | 47. | 26 | 31. | 83. | 26. | 50. | 71. | 50. | 62. | 37. | 23. | 26. | 0.2069 | FvH4_3g043 | | 249364 | 250322 | | |
| 24 | GENE30470 | 2 | 8 | 8 | 4 | 9 | 4 | 9 | 5 | 8 | 4 | 7 | 4 | 8 | 3 | 2062 | 70 | Fvb3 | 9 | 7 | alternative NAD(P)H dehydrogenase 1 | |
| 305 | | 11 | 13 | 12 | 11 | 11 | 19 | 83. | 18 | 16 | 18 | 11 | 11 | 12 | 13 | 0.2606 | FvH4_3g043 | | 249033 | 249317 | | |
| 25 | GENE30471 | 4 | 0 | 6 | 1 | 6 | 5 | 6 | 3 | 8 | 2 | 1 | 2 | 8 | 7 | 6534 | 60 | Fvb3 | 0 | 6 | bifunctional nuclease i | |
| 305 | | 12 | 25 | 41 | 41 | 51 | 36 | 27 | 39 | 45 | 59 | 23 | 69 | 39 | 20 | 0.3144 | FvH4_3g043 | | 248642 | 249000 | | |
| 26 | GENE30472 | 1 | 7 | 2 | 7 | 0 | 6 | 9 | 4 | 8 | 0 | 7 | 4 | 1 | 5 | 8628 | 50 | Fvb3 | 7 | 4 | endonuclease 4 | |
| 305 | | ## | ## | ## | ## | ## | 97 | ## | ## | ## | ## | ## | ## | 93 | ## | 0.1145 | FvH4_3g043 | | 248391 | 248575 | | |
| 27 | GENE30473 | ## | ## | ## | ## | ## | 64 | ## | ## | ## | ## | ## | ## | 31 | ## | 1197 | 40 | Fvb3 | 4 | 8 | oxidoreductase, zinc-binding dehydrogenase family protein | |
| 305 | | 70. | 48. | 55. | 50. | 38. | 10 | 44. | 55. | 63. | 51. | 14 | 81. | 29. | 84. | 0.3919 | FvH4_3g043 | | 248172 | 248215 | | |
| 28 | GENE30474 | 9 | 2 | 3 | 9 | 9 | 1 | 2 | 5 | 8 | 5 | 1 | 4 | 7 | 9 | 4898 | 30 | Fvb3 | 8 | 9 | RING/U-box superfamily protein | |
| 305 | | 11 | 14 | 20 | 17 | 13 | 14 | 16 | 13 | 13 | 11 | 10 | 15 | 17 | 14 | 0.1986 | FvH4_3g043 | | 247874 | 248046 | | |
| 29 | GENE30475 | 93 | 56 | 24 | 01 | 50 | 45 | 81 | 52 | 88 | 42 | 30 | 25 | 11 | 16 | 2954 | 20 | Fvb3 | 1 | 1 | oxidoreductase, zinc-binding dehydrogenase family protein | |
| 305 | | ## | ## | ## | ## | ## | 93 | 75 | ## | 85 | ## | 83 | ## | ## | 66 | 0.7481 | FvH4_3g043 | | 246793 | 247051 | | |
| 31 | GENE30477 | ## | ## | ## | ## | ## | 61 | 91 | ## | 84 | ## | 55 | ## | ## | 33 | 0299 | 00 | Fvb3 | 7 | 5 | Putative lysine decarboxylase family protein | |
| 305 | | 49. | 62. | | 49. | 44. | 34. | 41. | 41. | 42. | 43. | | 47. | 45. | 43. | 0.3408 | FvH4_3g042 | | 246311 | 246360 | | |
| 32 | GENE30478 | 6 | 9 | 46 | 4 | 9 | 2 | 5 | 7 | 6 | 6 | 39 | 2 | 7 | 9 | 8778 | 90 | Fvb3 | 4 | 8 | basic helix-loop-helix (bHLH) DNA-binding family protein | |
| 305 | | 7.9 | 10. | 12. | | | 9.5 | 11. | 6.8 | 10. | 9.9 | 9.7 | 10. | 8.6 | 11. | 0.7222 | FvH4_3g042 | | 245666 | 245715 | | |
| 33 | GENE30479 | 1 | 2 | 2 | 12 | 7.6 | 2 | 3 | 2 | 5 | 9 | 7 | 9 | 5 | 6 | 7987 | 80 | Fvb3 | 9 | 7 | | 0 |
| 305 | | 45 | 61 | 96 | 11 | 43 | 42 | 34 | 54 | 10 | 87 | 16 | 99 | 69 | 16 | 0.0499 | FvH4_3g042 | | 241481 | 241955 | | |
| 35 | GENE30482 | 6 | 2 | 7 | 23 | 6 | 4 | 6 | 9 | 37 | 9 | 78 | 4 | 8 | 71 | 2258 | 70 | Fvb3 | 3 | 4 | KNOX/ELK homeobox transcription factor | |
| 306 | | 28. | 26. | 25. | 21. | 23. | 25. | 22. | 26. | 24. | 20. | 28. | 23. | 20. | 27. | 0.8406 | FvH4_3g034 | | 190076 | 190255 | | |
| 82 | GENE30630 | 9 | 6 | 3 | 5 | 5 | 7 | 9 | 8 | 5 | 5 | 4 | 7 | 9 | 4 | 3135 | 80 | Fvb3 | 5 | 0 | 1-amino-cyclopropane-1-carboxylate synthase 8 | |
| 306 | | 49. | 55. | 11 | 11 | 11 | 84. | 13 | 30. | | 35. | 44. | 38. | 38. | 33. | 0.0026 | FvH4_3g034 | | 191064 | 191408 | | |
| 83 | GENE30631 | 4 | 2 | 6 | 1 | 7 | 3 | 0 | 9 | 31 | 2 | 2 | 2 | 8 | 1 | 6259 | 82 | Fvb3 | 3 | 2 | S-locus lectin protein kinase family protein | |
| 306 | | 55. | 42. | 48. | | 53. | 61. | 49. | 42. | | 50. | 47. | 46. | 38. | 47. | 0.0574 | FvH4_3g034 | | 188533 | 188851 | | |
| 85 | GENE30633 | 5 | 7 | 8 | 66 | 3 | 5 | 4 | 9 | 52 | 4 | 3 | 4 | 5 | 4 | 3542 | 51 | Fvb3 | 7 | 4 | S-locus lectin protein kinase family protein | |
| 306 | | 86. | 86. | 95. | 92. | 70. | 90. | 12 | 84. | 86. | 69. | 10 | 71. | 67. | | 0.1669 | FvH4_3g034 | | 186848 | 187225 | | |
| 87 | GENE30635 | 5 | 5 | 4 | 4 | 9 | 7 | 0 | 4 | 1 | 2 | 9 | 6 | 4 | 73 | 6653 | 33 | Fvb3 | 6 | 0 | S-domain-1 29 | |
| 306 | | 17. | 14. | 17. | 15. | 14. | | 13. | 15. | 17. | 14. | 13. | 14. | 13. | 13. | 0.1721 | FvH4_3g034 | | 186404 | 186779 | | |
| 88 | GENE30636 | 7 | 9 | 1 | 7 | 8 | 16 | 6 | 4 | 3 | 1 | 7 | 6 | 7 | 4 | 5298 | 32 | Fvb3 | 2 | 2 | S-locus lectin protein kinase family protein | |
| 306 | | 41 | 50 | 42 | 78 | 35 | 54 | 38 | 44 | 47 | 41 | 48 | 20 | 53 | 51 | 0.4952 | FvH4_3g034 | | 185737 | 186256 | | |
| 89 | GENE30637 | 5 | 9 | 4 | 7 | 3 | 6 | 5 | 8 | 6 | 1 | 2 | 7 | 4 | 7 | 2371 | 31 | Fvb3 | 5 | 8 | S-locus lectin protein kinase family protein | |
| 306 | | 11 | 17 | 17 | 25 | 28 | 28 | 18 | 22 | 25 | 26 | 19 | 17 | 21 | 16 | 0.9241 | FvH4_3g034 | | 185040 | 185553 | | |
| 90 | GENE30638 | 4 | 0 | 8 | 3 | 8 | 0 | 8 | 3 | 8 | 1 | 7 | 3 | 1 | 9 | 9993 | 30 | Fvb3 | 6 | 3 | receptor kinase 3 | |
| 306 | | 46. | 32. | 38. | 37. | 31. | 27. | 47. | 49. | 66. | 56. | 25. | | | 33. | 0.4670 | FvH4_3g034 | | 184671 | 185004 | | |
| 91 | GENE30639 | 8 | 6 | 9 | 5 | 3 | 1 | 8 | 3 | 8 | 5 | 7 | 34 | 31 | 2 | 7595 | 20 | Fvb3 | 5 | 0 | receptor kinase 3 | |
| 306 | | 14 | 26 | 23 | 51 | 11 | 28 | 23 | 86. | 14 | 14 | 95. | 87. | 32 | 59. | 0.0667 | FvH4_3g034 | | 184252 | 184709 | | |
| 92 | GENE30640 | 9 | 2 | 0 | 9 | 6 | 7 | 0 | 5 | 2 | 3 | 4 | 7 | 4 | 5 | 7777 | 10 | Fvb3 | 6 | 5 | S-locus lectin protein kinase family protein | |
| 306 | | 93 | 37 | 58 | 52 | 36 | 45 | 38 | 38 | 44 | 42 | 41 | 47 | 42 | 38 | 0.2687 | FvH4_3g034 | | 183890 | 184224 | | |
| 93 | GENE30641 | 2 | 3 | 2 | 2 | 1 | 6 | 4 | 1 | 6 | 4 | 9 | 7 | 4 | 3 | 5975 | 00 | Fvb3 | 0 | 0 | lysine decarboxylase family protein | |
| 306 | | 35. | 52. | 49. | | | 33. | 31. | 36. | 51. | 50. | 46. | 44. | | 40. | 0.7533 | FvH4_3g033 | | 183245 | 183877 | | |
| 94 | GENE30642 | 7 | 1 | 3 | 71 | 27 | 8 | 6 | 2 | 4 | 5 | 3 | 6 | 45 | 6 | 0227 | 90 | Fvb3 | 0 | 6 | S-locus lectin protein kinase family protein | |
| 306 | | 20 | 47 | 39 | 54 | 39 | 54 | 51 | 27 | 25 | 36 | 35 | 28 | 47 | 35 | 0.0888 | FvH4_3g033 | | 182905 | 183285 | | |
| 95 | GENE30643 | 7 | 1 | 7 | 8 | 5 | 0 | 2 | 5 | 8 | 6 | 7 | 7 | 3 | 3 | 9744 | 70 | Fvb3 | 7 | 1 | S-locus lectin protein kinase family protein | |
| 306 | | 10 | 70 | 89 | 11 | 59 | 67 | 80 | 55 | 22 | 58 | 52 | 46 | 10 | 28 | 0.0244 | FvH4_3g033 | | 181808 | 182133 | | |
| 96 | GENE30644 | 28 | 2 | 9 | 33 | 7 | 6 | 9 | 9 | 0 | 2 | 8 | 9 | 01 | 9 | 174 | 50 | Fvb3 | 9 | 2 | S-locus lectin protein kinase family protein | |

| | | | | | | | | | | | | | | | | | | | | | | |
|---|---|---|---|---|---|---|---|---|---|---|---|---|---|---|---|---|---|---|---|---|---|---|
| 306 | | | 14. | 16. | 13. | 19. | 14. | 22. | 13. | 11. | | 10. | 12. | 11. | 13. | 0.0054 | FvH4_3g033 | | 181256 | 181546 | | |
| 97 | GENE30645 | 23 | 7 | 8 | 7 | 2 | 7 | 8 | 6 | 2 | 9.5 | 9 | 5 | 8 | 3 | 6086 | 25 | Fvb3 | 9 | 2 | | 0 |
| 306 | | 85. | 69. | 11 | 11 | 98. | 66. | 14 | 77. | 10 | 78. | 11 | 11 | 14 | 11 | 0.5152 | FvH4_3g033 | | 180595 | 180656 | | |
| 98 | GENE30646 | 8 | 3 | 4 | 2 | 8 | 9 | 6 | 6 | 8 | 6 | 5 | 7 | 3 | 9 | 1273 | 24 | Fvb3 | 9 | 2 | | 0 |
| 307 | | 15 | 24 | 14 | 22 | 21 | 12 | 16 | 23 | 85. | 24 | 13 | 16 | 24 | 94. | 0.7529 | FvH4_3g033 | | 178699 | 179034 | | |
| 01 | GENE30649 | 6 | 4 | 3 | 8 | 6 | 8 | 0 | 8 | 1 | 4 | 0 | 2 | 8 | 2 | 314 | 20 | Fvb3 | 2 | 9 | S-locus lectin protein kinase family protein | |
| 307 | | 11 | 20 | 16 | 26 | 12 | 12 | 18 | 11 | 88. | 13 | 14 | 12 | 11 | 11 | 0.0587 | FvH4_3g033 | | 177999 | 178352 | | |
| 02 | GENE30650 | 4 | 5 | 3 | 2 | 6 | 6 | 1 | 6 | 3 | 6 | 9 | 7 | 9 | 2 | 5793 | 10 | Fvb3 | 4 | 7 | S-locus lectin protein kinase family protein | |
| 307 | | 20. | 23. | 18. | 23. | 23. | 19. | 19. | 21. | 26. | 22. | 17. | 19. | 24. | 21. | 0.5972 | FvH4_3g033 | | 177560 | 177923 | | |
| 03 | GENE30651 | 1 | 2 | 6 | 4 | 2 | 4 | 4 | 6 | 3 | 9 | 3 | 2 | 1 | 3 | 1195 | 01 | Fvb3 | 3 | 7 | S-locus lectin protein kinase family protein | |
| 307 | | | 17. | 19. | | 20. | 19. | 21. | 17. | 25. | 26. | 23. | | 27. | | 0.0277 | FvH4_3g033 | | 177108 | 177442 | | |
| 04 | GENE30652 | 17 | 6 | 4 | 22 | 4 | 5 | 4 | 4 | 7 | 3 | 5 | 20 | 2 | 28 | 4372 | 00 | Fvb3 | 3 | 8 | S-locus lectin protein kinase family protein | |
| 307 | | 40. | | 52. | 62. | 50. | 49. | | 59. | 73. | 53. | 68. | 50. | 60. | 60. | 0.0335 | FvH4_3g032 | | 176845 | 176997 | | |
| 05 | GENE30653 | 4 | 44 | 8 | 6 | 8 | 1 | 56 | 2 | 4 | 1 | 5 | 1 | 9 | 4 | 9799 | 80 | Fvb3 | 0 | 9 | RNAse l inhibitor protein 2 | |
| 307 | | 65 | 59 | 40 | 30 | 66 | 27 | 10 | 49 | 47 | 72 | 40 | 70 | 15 | 42 | 0.5475 | FvH4_3g032 | | 176417 | 176634 | | |
| 06 | GENE30654 | 1 | 0 | 2 | 4 | 9 | 8 | 86 | 1 | 3 | 9 | 6 | 3 | 65 | 7 | 2617 | 60 | Fvb3 | 4 | 6 | cytokinin oxidase/dehydrogenase 6 | |
| 307 | | 24. | 24. | 24. | 25. | 26. | 28. | 26. | 21. | 27. | 23. | 27. | 26. | 25. | | 0.9584 | FvH4_3g032 | | 176250 | 176323 | | |
| 07 | GENE30655 | 5 | 8 | 3 | 6 | 3 | 2 | 2 | 7 | 3 | 5 | 8 | 8 | 1 | 28 | 3804 | 51 | Fvb3 | 2 | 2 | | 0 |
| 307 | | 43. | 37. | 36. | 46. | 37. | 27. | 56. | 27. | 39. | 29. | 41. | 31. | 37. | 48. | 0.3318 | FvH4_3g032 | | 175796 | 176045 | | |
| 08 | GENE30656 | 8 | 1 | 6 | 8 | 7 | 6 | 9 | 9 | 2 | 1 | 4 | 5 | 4 | 2 | 7116 | 50 | Fvb3 | 7 | 9 | crooked neck protein, putative / cell cycle protein, putative | |
| 307 | | 31. | 31. | 29. | 35. | 33. | 28. | 33. | 28. | 36. | 36. | 35. | 29. | 41. | 40. | 0.1194 | FvH4_3g032 | | 175416 | 175758 | | |
| 09 | GENE30657 | 1 | 8 | 7 | 3 | 6 | 8 | 3 | 4 | 8 | 9 | 2 | 7 | 9 | 1 | 7463 | 43 | Fvb3 | 4 | 6 | S-locus lectin protein kinase family protein | |
| 307 | | 58 | 66 | 42 | 98 | 55 | 60 | 30 | 36 | 40 | 46 | 38 | 21 | 26 | 10 | 0.0154 | FvH4_3g032 | | 174642 | 175009 | | |
| 10 | GENE30658 | 4 | 4 | 8 | 7 | 5 | 6 | 5 | 9 | 5 | 4 | 8 | 8 | 2 | 9 | 8426 | 42 | Fvb3 | 3 | 9 | S-locus lectin protein kinase family protein | |
| 307 | | 29. | 22. | 29. | | 31. | 25. | 29. | 18. | 21. | | 22. | 21. | 25. | 23. | 0.0017 | FvH4_3g032 | | 174013 | 174398 | | |
| 11 | GENE30659 | 9 | 1 | 4 | 32 | 1 | 6 | 4 | 7 | 5 | 18 | 9 | 7 | 9 | 7 | 7957 | 41 | Fvb3 | 2 | 2 | S-locus lectin protein kinase family protein | |
| 307 | | 36. | 33. | 61. | 89. | 71. | 64. | 60. | 42. | | | 53. | 43. | 52. | 38. | 0.5695 | FvH4_3g032 | | 173425 | 173767 | | |
| 12 | GENE30660 | 2 | 4 | 6 | 1 | 8 | 7 | 5 | 6 | 69 | 79 | 8 | 2 | 7 | 9 | 1732 | 40 | Fvb3 | 1 | 0 | S-locus lectin protein kinase family protein | |
| 307 | | 66 | 57 | 54 | 67 | 61 | 53 | 71 | 51 | 43 | 39 | 35 | 29 | 34 | 34 | 4.9149 | FvH4_3g032 | | 172920 | 173299 | | |
| 13 | GENE30661 | 08 | 32 | 17 | 63 | 04 | 98 | 60 | 38 | 56 | 23 | 46 | 26 | 73 | 13 | E-05 | 31 | Fvb3 | 1 | 0 | S-locus lectin protein kinase family protein | |
| 307 | | 49. | 66. | 74. | 90. | 90. | 12 | 69. | 11 | 76. | 58. | | | 69. | 47. | 0.4774 | FvH4_3g032 | | 171785 | 172195 | | |
| 14 | GENE30662 | 1 | 5 | 7 | 7 | 8 | 6 | 9 | 8 | 5 | 1 | 76 | 57 | 8 | 1 | 1716 | 30 | Fvb3 | 1 | 2 | S-locus lectin protein kinase family protein | |
| 307 | | 16 | 15 | 16 | 16 | 78 | 12 | 17 | 98 | 11 | 84 | 13 | 10 | 17 | 13 | 0.1658 | FvH4_3g032 | | 171212 | 171507 | | |
| 15 | GENE30663 | 19 | 97 | 19 | 24 | 6 | 44 | 79 | 6 | 84 | 5 | 82 | 14 | 50 | 20 | 323 | 20 | Fvb3 | 3 | 8 | plant U-box 8 | |
| 307 | | ## | ## | ## | ## | ## | ## | 93 | 65 | ## | 96 | ## | 91 | ## | 70 | 0.0967 | FvH4_3g032 | | 170702 | 171010 | | |
| 16 | GENE30664 | ## | ## | ## | ## | ## | ## | 36 | 12 | ## | 59 | ## | 83 | ## | 67 | 6101 | 10 | Fvb3 | 4 | 1 | flavodoxin-like quinone reductase 1 | |
| 307 | | 57. | 56. | 66. | 58. | 62. | 73. | 81. | 46. | 79. | 62. | 94. | | 60. | 87. | 0.2628 | FvH4_3g032 | | 170111 | 170415 | | |
| 17 | GENE30665 | 1 | 8 | 9 | 6 | 5 | 1 | 6 | 9 | 1 | 4 | 1 | 89 | 9 | 2 | 6617 | 00 | Fvb3 | 0 | 8 | basic helix-loop-helix (bHLH) DNA-binding superfamily protein | |
| 307 | | 33. | 22. | 27. | | 34. | 29. | 25. | | 26. | 31. | 29. | 33. | 28. | 35. | 0.8593 | FvH4_3g031 | | 168752 | 168881 | | |
| 19 | GENE30667 | 3 | 3 | 5 | 42 | 7 | 2 | 4 | 27 | 8 | 1 | 3 | 1 | 2 | 5 | 9642 | 70 | Fvb3 | 0 | 8 | basic helix-loop-helix (bHLH) DNA-binding superfamily protein | |
| 307 | | 22 | 24 | 28 | 32 | 28 | 33 | 33 | 23 | 47 | 25 | 38 | 31 | 21 | 36 | 0.4701 | FvH4_3g031 | | 168545 | 168659 | | |
| 20 | GENE30668 | 55 | 85 | 10 | 99 | 49 | 41 | 50 | 88 | 36 | 36 | 53 | 06 | 94 | 12 | 6552 | 60 | Fvb3 | 6 | 2 | Tetratricopeptide repeat (TPR)-like superfamily protein | |
| 307 | | ## | 74 | 98 | 86 | ## | 64 | ## | 75 | ## | 79 | 78 | 70 | ## | 95 | 0.9156 | FvH4_3g031 | | 168058 | 168522 | | |
| 21 | GENE30669 | ## | 03 | 02 | 25 | ## | 92 | ## | 97 | ## | 77 | 15 | 83 | ## | 90 | 6098 | 50 | Fvb3 | 3 | 5 | terpene synthase 14 | |
| 307 | | 12 | 63 | 13 | 95. | 98 | 13 | 64. | 36 | 13 | 21. | 86 | 13 | 13 | 55. | 0.7510 | FvH4_3g031 | | 167843 | 168023 | S-adenosyl-L-methionine-dependent methyltransferases | |
| 22 | GENE30670 | 46 | 6 | 8 | 7 | 9 | 72 | 3 | 9 | 20 | 4 | 2 | 07 | 09 | 3 | 5379 | 30 | Fvb3 | 0 | 5 | superfamily protein | |

| | | | | | | | | | | | | | | | | | | | | | | |
|---|---|---|---|---|---|---|---|---|---|---|---|---|---|---|---|---|---|---|---|---|---|---|
| 307 | | 38 | 41 | 64 | 46 | 45 | 34 | 49 | 26 | 49 | 44 | 79 | 43 | 69 | 84 | 0.2527 | FvH4_3g042 | | 240287 | 240368 | | |
| 24 | GENE30672 | 7 | 4 | 2 | 7 | 7 | 1 | 5 | 2 | 4 | 6 | 9 | 1 | 4 | 6 | 8373 | 60 | Fvb3 | 0 | 8 | Protein of unknown function (DUF1191) | |
| 307 | | 20 | 13 | 24 | 18 | 18 | 19 | 23 | 10 | 18 | 20 | 13 | 18 | 19 | 17 | 0.1348 | FvH4_3g042 | | 239402 | 239585 | | |
| 25 | GENE30673 | 33 | 23 | 91 | 93 | 89 | 36 | 57 | 43 | 93 | 03 | 01 | 33 | 29 | 02 | 5725 | 50 | Fvb3 | 4 | 0 | Putative lysine decarboxylase family protein | |
| 307 | | 18 | 19 | 53 | 50 | 44 | 33 | 52 | 15 | 43 | 37 | 57 | 55 | 64 | 71 | 0.2959 | FvH4_3g042 | | 239281 | 239332 | | |
| 26 | GENE30674 | 98 | 15 | 65 | 49 | 79 | 92 | 55 | 21 | 20 | 10 | 49 | 45 | 19 | 28 | 0918 | 42 | Fvb3 | 1 | 3 | | 0 |
| 307 | | 19 | 17 | 15 | 16 | 17 | 13 | 13 | 14 | 16 | 11 | 11 | 13 | 12 | 13 | 0.0210 | FvH4_3g042 | | 238814 | 238991 | | |
| 27 | GENE30675 | 70 | 98 | 57 | 53 | 55 | 87 | 27 | 24 | 61 | 91 | 16 | 95 | 89 | 15 | 3939 | 41 | Fvb3 | 3 | 6 | Protein of unknown function (DUF300) | |
| 307 | | 86. | 90. | 20 | 16 | 25 | 16 | 13 | 98. | 24 | 22 | | 24 | 36 | 58. | 0.5260 | FvH4_3g042 | | 238623 | 238754 | | |
| 28 | GENE30676 | 1 | 4 | 4 | 6 | 7 | 8 | 9 | 3 | 9 | 5 | 93 | 2 | 1 | 9 | 9097 | 40 | Fvb3 | 3 | 6 | | 0 |
| 307 | | 10 | 15 | 12 | 97. | 19 | 18 | 26 | 13 | 28 | 34 | 15 | 15 | 17 | 21 | 0.2230 | FvH4_3g042 | | 237268 | 237802 | | |
| 29 | GENE30677 | 9 | 2 | 9 | 3 | 8 | 0 | 3 | 4 | 0 | 0 | 9 | 0 | 3 | 9 | 7154 | 33 | Fvb3 | 7 | 4 | | 0 |
| 307 | | 48 | 47 | 42 | 46 | 47 | 36 | 38 | 36 | 45 | 33 | 35 | 43 | 33 | 40 | 0.0476 | FvH4_3g042 | | 236928 | 237021 | | |
| 30 | GENE30678 | 18 | 40 | 27 | 42 | 87 | 56 | 42 | 15 | 71 | 72 | 19 | 04 | 43 | 11 | 8259 | 32 | Fvb3 | 0 | 6 | Protein of unknown function (DUF300) | |
| 307 | | 68 | 72 | 52 | 74 | 65 | 71 | 69 | 54 | 66 | 59 | 59 | 59 | 63 | 47 | 0.0260 | FvH4_3g042 | | 236129 | 236549 | | |
| 31 | GENE30679 | 04 | 49 | 44 | 91 | 76 | 29 | 34 | 65 | 00 | 22 | 43 | 43 | 60 | 75 | 7224 | 20 | Fvb3 | 5 | 0 | ARF-GAP domain 5 | |
| 307 | | | 11 | 68. | 75. | 13 | 14 | 15 | 11 | 29 | 15 | 15 | 13 | 77. | 19 | 0.1087 | FvH4_3g042 | | 235213 | 235827 | | |
| 32 | GENE30680 | 66 | 2 | 9 | 4 | 2 | 7 | 2 | 7 | 1 | 0 | 8 | 1 | 2 | 6 | 1419 | 00 | Fvb3 | 9 | 6 | homolog of yeast sucrose nonfermenting 4 | |
| 307 | | 16. | 15. | | 16. | 15. | 16. | 16. | 15. | 15. | 18. | 18. | 18. | 16. | | 0.1411 | FvH4_3g041 | | 233723 | 233909 | | |
| 34 | GENE30682 | 8 | 3 | 17 | 3 | 3 | 2 | 5 | 4 | 1 | 2 | 5 | 1 | 6 | 19 | 0745 | 80 | Fvb3 | 7 | 9 | 1-amino-cyclopropane-1-carboxylate synthase 8 | |
| 307 | | 17. | 45. | 49. | | 20. | | 20. | 49. | 37. | 42. | 10 | 10 | 31. | 50. | 0.0639 | FvH4_3g041 | | 233165 | 233538 | | |
| 35 | GENE30683 | 5 | 3 | 8 | 52 | 5 | 20 | 6 | 5 | 1 | 3 | 2 | 1 | 4 | 2 | 7179 | 70 | Fvb3 | 1 | 1 | floral meristem identity control protein LEAFY (LFY) | |
| 307 | | 25. | 28. | 31. | 25. | 25. | 27. | 26. | | 32. | 29. | 31. | 28. | 34. | | 0.1695 | FvH4_3g041 | | 233005 | 233099 | | |
| 36 | GENE30684 | 2 | 6 | 8 | 3 | 4 | 2 | 7 | 21 | 4 | 7 | 2 | 8 | 9 | 32 | 2313 | 61 | Fvb3 | 1 | 6 | | 0 |
| 307 | | 90 | 12 | 25 | 31 | 12 | 23 | 24 | 97 | 10 | 13 | 14 | 17 | 15 | 11 | 0.0810 | FvH4_3g041 | | 232271 | 232466 | | |
| 37 | GENE30685 | 1 | 66 | 99 | 19 | 79 | 09 | 89 | 9 | 25 | 77 | 65 | 27 | 11 | 53 | 3084 | 60 | Fvb3 | 0 | 5 | Protein of unknown function (DUF761) | |
| 307 | | 48 | 68 | 49 | 47 | 68 | 70 | 71 | 73 | 74 | 65 | 57 | 60 | 62 | 68 | 0.2773 | FvH4_3g041 | | 231678 | 232161 | | |
| 39 | GENE30687 | 04 | 39 | 24 | 37 | 07 | 19 | 63 | 49 | 15 | 75 | 69 | 93 | 04 | 92 | 1852 | 50 | Fvb3 | 4 | 3 | Tetratricopeptide repeat (TPR)-like superfamily protein | |
| 307 | | ## | ## | ## | ## | ## | ## | ## | ## | ## | ## | ## | ## | ## | ## | 0.0104 | FvH4_3g041 | | 231398 | 231624 | Vacuolar protein sorting-associated protein VPS28 family | |
| 40 | GENE30688 | ## | ## | ## | ## | ## | ## | ## | ## | ## | ## | ## | ## | ## | ## | 4264 | 40 | Fvb3 | 6 | 2 | protein | |
| 307 | | | 25. | 27. | 32. | 26. | 34. | 28. | 24. | | 28. | 35. | 29. | 26. | 45. | 0.3807 | FvH4_3g041 | | 231192 | 231380 | | |
| 41 | GENE30689 | 24 | 8 | 9 | 6 | 5 | 5 | 4 | 4 | 30 | 2 | 4 | 1 | 7 | 3 | 4744 | 30 | Fvb3 | 5 | 8 | syntaxin of plants 124 | |
| 307 | | 46. | | 66. | 56. | 39. | 33. | | 35. | 27. | | 32. | 41. | | 35. | 0.0098 | FvH4_3g041 | | 228850 | 229223 | Haloacid dehalogenase-like hydrolase (HAD) superfamily | |
| 43 | GENE30691 | 1 | 53 | 6 | 7 | 4 | 2 | 47 | 2 | 5 | 28 | 4 | 9 | 34 | 9 | 348 | 00 | Fvb3 | 4 | 0 | protein | |
| 307 | | 90 | 92 | 95 | ## | 76 | 83 | 89 | 80 | 80 | 93 | 83 | 71 | 87 | 81 | 0.0870 | FvH4_3g040 | | 227955 | 228396 | | |
| 44 | GENE30692 | 58 | 41 | 70 | ## | 44 | 95 | 78 | 60 | 99 | 31 | 33 | 80 | 19 | 57 | 6282 | 90 | Fvb3 | 9 | 1 | sphingosine kinase 1 | |
| 307 | | 23. | 19. | 17. | | 18. | | 17. | | 16. | | 19. | 15. | 14. | 18. | 0.1159 | FvH4_3g040 | | 227359 | 227655 | | |
| 45 | GENE30693 | 1 | 8 | 8 | 21 | 1 | 14 | 3 | 14 | 6 | 15 | 6 | 9 | 8 | 9 | 5511 | 70 | Fvb3 | 2 | 6 | | 0 |
| 307 | | 89 | 14 | 13 | 25 | 94 | 14 | 17 | 89 | 66 | 10 | 10 | 72 | 11 | 12 | 0.0552 | FvH4_3g040 | | 226761 | 227297 | | |
| 46 | GENE30694 | 4 | 34 | 44 | 82 | 3 | 76 | 52 | 6 | 2 | 56 | 37 | 4 | 69 | 46 | 2903 | 60 | Fvb3 | 4 | 5 | Calmodulin-binding protein | |
| 307 | | 82 | 64 | 83 | 73 | 97 | 10 | 11 | 10 | 69 | 70 | 68 | 60 | 74 | 94 | 0.2782 | FvH4_3g040 | | 226048 | 226921 | | |
| 47 | GENE30695 | 3 | 2 | 4 | 8 | 7 | 26 | 15 | 64 | 4 | 3 | 4 | 7 | 9 | 9 | 4823 | 50 | Fvb3 | 8 | 0 | Transducin/WD40 repeat-like superfamily protein | |
| 307 | | 23 | 34 | 27 | 30 | 29 | 24 | 18 | 35 | 26 | 38 | 31 | 34 | 38 | 17 | 0.1794 | FvH4_3g040 | | 225490 | 226002 | | |
| 48 | GENE30696 | 18 | 95 | 94 | 11 | 01 | 16 | 68 | 55 | 80 | 51 | 21 | 12 | 74 | 84 | 7174 | 40 | Fvb3 | 8 | 0 | riboflavin kinase/FMN hydrolase | |
| 307 | | 38. | 43. | 38. | 43. | 39. | 31. | 42. | 27. | 40. | 38. | 43. | | 39. | | 0.6038 | FvH4_3g040 | | 225241 | 225272 | | |
| 49 | GENE30697 | 5 | 3 | 7 | 1 | 8 | 7 | 9 | 8 | 1 | 6 | 8 | 47 | 1 | 55 | 7658 | 31 | Fvb3 | 6 | 7 | | 0 |

| | | | | | | | | | | | | | | | | | | | | | | |
|---|---|---|---|---|---|---|---|---|---|---|---|---|---|---|---|---|---|---|---|---|---|---|
| 307 | | 36 | 64 | 48 | 48 | 50 | 60 | 43 | 64 | 41 | 53 | 35 | 49 | 36 | 23 | 0.2923 | FvH4_3g040 | | 224823 | 225199 | | |
| 50 | GENE30698 | 6 | 9 | 7 | 3 | 6 | 9 | 0 | 5 | 0 | 6 | 4 | 6 | 6 | 5 | 44 | 30 | Fvb3 | 8 | 1 | Transducin/WD40 repeat-like superfamily protein | |
| 307 | | | 52. | 11 | 48 | 96. | 19 | 75. | 11 | 78. | 80. | | 63. | 58. | 61. | 0.2086 | FvH4_3g040 | | 224525 | 224648 | | |
| 51 | GENE30699 | 81 | 3 | 5 | 4 | 2 | 3 | 8 | 4 | 6 | 6 | 75 | 7 | 1 | 8 | 7116 | 20 | Fvb3 | 0 | 4 | F-box family protein | |
| 307 | | 25. | 30. | 34. | 32. | 30. | 32. | 33. | 29. | 31. | 32. | 32. | 33. | 36. | 36. | 0.2172 | FvH4_3g040 | | 224144 | 224454 | | |
| 52 | GENE30700 | 3 | 5 | 4 | 7 | 5 | 4 | 2 | 5 | 6 | 3 | 4 | 8 | 7 | 5 | 3504 | 12 | Fvb3 | 3 | 2 | actin related protein 2 | |
| 307 | | 9.6 | | 8.9 | 8.4 | 7.8 | 10. | 8.6 | 8.6 | 11. | 8.3 | 7.4 | 7.2 | 9.0 | 8.4 | 0.6180 | FvH4_3g040 | | 223878 | 224038 | | |
| 53 | GENE30701 | 1 | 9.1 | 9 | 1 | 1 | 5 | 3 | 7 | 5 | 7 | 7 | 7 | 3 | 3 | 3243 | 11 | Fvb3 | 4 | 3 | | 0 |
| 307 | | 12 | 15 | 28 | 34 | 21 | 10 | 30 | 12 | 31 | 17 | 30 | 33 | 25 | 18 | 0.6137 | FvH4_3g040 | | 223325 | 223527 | | |
| 54 | GENE30702 | 8 | 3 | 5 | 2 | 6 | 7 | 2 | 8 | 8 | 1 | 9 | 7 | 7 | 3 | 3135 | 10 | Fvb3 | 4 | 3 | Auxin-responsive GH3 family protein | |
| 307 | | 26 | 20 | 26 | 21 | 15 | 17 | 16 | 16 | 13 | 15 | 18 | 20 | 18 | 19 | 0.1463 | FvH4_3g040 | | 222812 | 223049 | | |
| 55 | GENE30703 | 07 | 86 | 82 | 92 | 41 | 51 | 22 | 76 | 86 | 97 | 04 | 96 | 31 | 28 | 1639 | 00 | Fvb3 | 5 | 9 | SGNH hydrolase-type esterase superfamily protein | |
| 307 | | 13. | 16. | 21. | | 15. | 15. | 15. | 14. | 16. | 16. | 15. | 16. | 16. | | 0.6179 | FvH4_3g039 | | 222463 | 222586 | | |
| 56 | GENE30704 | 4 | 9 | 8 | 17 | 5 | 9 | 4 | 7 | 2 | 3 | 7 | 3 | 8 | 16 | 9757 | 90 | Fvb3 | 6 | 8 | F-box family protein | |
| 307 | | 59 | 33 | 39 | 35 | 39 | 62 | 44 | 48 | 16 | 22 | 22 | 15 | 44 | 20 | 0.0215 | FvH4_3g039 | | 222024 | 222364 | | |
| 57 | GENE30705 | 68 | 94 | 18 | 10 | 58 | 59 | 34 | 23 | 27 | 54 | 33 | 32 | 43 | 64 | 4559 | 80 | Fvb3 | 7 | 8 | | 0 |
| 307 | | ## | ## | ## | ## | ## | ## | ## | ## | 96 | ## | ## | ## | ## | ## | 0.0059 | FvH4_3g039 | | 221749 | 221832 | | |
| 58 | GENE30706 | ## | ## | ## | ## | ## | ## | ## | ## | 67 | ## | ## | ## | ## | ## | 8277 | 70 | Fvb3 | 8 | 8 | | 0 |
| 307 | | ## | ## | ## | ## | ## | ## | ## | ## | ## | ## | ## | ## | ## | ## | 0.9542 | FvH4_3g039 | | 220947 | 221408 | | |
| 59 | GENE30707 | ## | ## | ## | ## | ## | ## | ## | ## | ## | ## | ## | ## | ## | ## | 8156 | 60 | Fvb3 | 9 | 2 | NAD(P)H dehydrogenase B2 | |
| 307 | | | 17. | | 17. | 23. | 19. | 19. | | 22. | 20. | 25. | 21. | 21. | 21. | 0.0882 | FvH4_3g039 | | 220766 | 220849 | | |
| 60 | GENE30708 | 17 | 7 | 24 | 2 | 1 | 8 | 8 | 22 | 6 | 3 | 6 | 5 | 9 | 2 | 5401 | 50 | Fvb3 | 5 | 6 | Dynein light chain type 1 family protein | |
| 307 | | 72. | 51. | 59. | 88. | 50. | 11 | 44. | | 52. | 81. | 62. | 38. | 66. | 62. | 0.3514 | FvH4_3g039 | | 220289 | 220506 | | |
| 61 | GENE30709 | 7 | 7 | 1 | 6 | 3 | 9 | 5 | 44 | 2 | 6 | 1 | 2 | 6 | 3 | 9013 | 40 | Fvb3 | 3 | 7 | F-box family protein | |
| 307 | | | 35. | 29. | 33. | 41. | 28. | 31. | 31. | 30. | 28. | 30. | | 43. | 31. | 0.9400 | FvH4_3g039 | | 219783 | 220132 | | |
| 62 | GENE30710 | 37 | 1 | 5 | 9 | 2 | 5 | 1 | 3 | 9 | 7 | 9 | 38 | 1 | 9 | 2831 | 32 | Fvb3 | 6 | 4 | zinc knuckle (CCHC-type) family protein | |
| 307 | | 35. | 36. | 54. | 47. | 45. | 49. | 42. | | 48. | | 60. | 58. | 48. | 64. | 0.2793 | FvH4_3g039 | | 219285 | 219733 | | |
| 63 | GENE30711 | 6 | 7 | 2 | 6 | 4 | 4 | 4 | 30 | 1 | 43 | 1 | 6 | 3 | 7 | 3673 | 31 | Fvb3 | 9 | 8 | | 0 |
| 307 | | 42 | 40 | 46 | 42 | 52 | 49 | 49 | 56 | 51 | 63 | 48 | 53 | 64 | 43 | 0.0366 | FvH4_3g039 | | 218579 | 218751 | | |
| 64 | GENE30712 | 57 | 19 | 92 | 42 | 98 | 31 | 97 | 55 | 73 | 02 | 99 | 76 | 82 | 02 | 6631 | 30 | Fvb3 | 7 | 8 | ARM repeat superfamily protein | |
| 307 | | 55. | | 45. | 44. | 42. | 24. | 42. | 27. | 38. | 27. | 40. | 40. | 50. | 35. | 0.4003 | FvH4_3g039 | | 218385 | 218480 | | |
| 65 | GENE30713 | 2 | 36 | 4 | 3 | 6 | 4 | 3 | 8 | 6 | 3 | 8 | 6 | 7 | 6 | 2547 | 20 | Fvb3 | 5 | 5 | | 0 |
| 307 | | 32. | | 66. | 62. | 43. | 36. | 64. | 31. | 98. | 36. | 73. | 65. | 56. | 14 | 0.1783 | FvH4_3g039 | | 218055 | 218335 | | |
| 66 | GENE30714 | 9 | 38 | 3 | 1 | 8 | 8 | 3 | 6 | 8 | 3 | 3 | 8 | 9 | 3 | 3003 | 10 | Fvb3 | 0 | 4 | Tetratricopeptide repeat (TPR)-like superfamily protein | |
| 307 | | ## | ## | ## | ## | ## | ## | ## | ## | ## | ## | ## | ## | ## | ## | 0.4165 | FvH4_3g039 | | 217252 | 217553 | | |
| 67 | GENE30715 | ## | ## | ## | ## | ## | ## | ## | ## | ## | ## | ## | ## | ## | ## | 4692 | 00 | Fvb3 | 4 | 8 | sugar transporter 1 | |
| 307 | | 22. | 19. | 18. | 17. | 18. | 20. | | 18. | 20. | 18. | 18. | 19. | 14. | 24. | 0.8552 | FvH4_3g038 | | 216617 | 216685 | | |
| 68 | GENE30716 | 2 | 2 | 8 | 6 | 4 | 8 | 19 | 3 | 6 | 1 | 6 | 7 | 4 | 6 | 8336 | 82 | Fvb3 | 2 | 4 | | 0 |
| 307 | | 12. | 14. | | | 13. | 13. | 14. | 13. | 13. | 12. | | 13. | 13. | 17. | 0.3791 | FvH4_3g038 | | 216417 | 216494 | | |
| 69 | GENE30717 | 9 | 7 | 13 | 13 | 3 | 2 | 1 | 2 | 9 | 4 | 14 | 5 | 9 | 4 | 0942 | 81 | Fvb3 | 4 | 3 | | 0 |
| 307 | | 22 | 25 | 31 | 28 | 30 | 29 | 35 | 22 | 30 | 31 | 26 | 30 | 36 | 23 | 0.9686 | FvH4_3g038 | | 215807 | 216193 | | |
| 70 | GENE30718 | 14 | 39 | 62 | 83 | 56 | 13 | 51 | 98 | 98 | 27 | 60 | 31 | 92 | 43 | 5874 | 80 | Fvb3 | 4 | 8 | PapD-like superfamily protein | |
| 307 | | 26. | 37. | 31. | 21. | 43. | 52. | 46. | 38. | 70. | 42. | 40. | 47. | | 31. | 0.5281 | FvH4_3g038 | | 215513 | 215679 | | |
| 71 | GENE30719 | 8 | 8 | 7 | 6 | 8 | 3 | 9 | 3 | 7 | 7 | 5 | 7 | 22 | 6 | 8457 | 70 | Fvb3 | 4 | 8 | | 0 |
| 307 | | 20 | 26 | 34 | 22 | 26 | 23 | 39 | 24 | 32 | 24 | 34 | 30 | 25 | 40 | 0.4412 | FvH4_3g038 | | 214813 | 214851 | uridine 5'-monophosphate synthase / UMP synthase (PYRE-F) | |
| 72 | GENE30720 | 11 | 11 | 86 | 15 | 14 | 09 | 71 | 91 | 13 | 56 | 19 | 65 | 07 | 22 | 2735 | 40 | Fvb3 | 7 | 6 | (UMPS) | |

| | | | | | | | | | | | | | | | | | | | | | |
|---|---|---|---|---|---|---|---|---|---|---|---|---|---|---|---|---|---|---|---|---|---|
| 307 | | 27. | 26. | 32. | 34. | 31. | 31. | 32. | | 37. | 34. | 37. | | 29. | 38. | 0.1282 | FvH4_3g038 | | 213990 | 214183 | |
| 74 | GENE30722 | 9 | 3 | 4 | 5 | 5 | 2 | 6 | 26 | 8 | 2 | 2 | 37 | 9 | 4 | 57 | 21 | Fvb3 | 6 | 4 | |
| 307 | | 26 | 87 | 48 | 33 | 98 | 74 | 11 | 80 | 11 | 96 | 90 | 80 | 41 | 14 | 0.2159 | FvH4_3g038 | | 213634 | 213715 | |
| 75 | GENE30723 | 2 | 3 | 5 | 6 | 5 | 2 | 54 | 0 | 20 | 2 | 5 | 6 | 4 | 17 | 9967 | 10 | Fvb3 | 8 | 4 | Wound-responsive family protein |
| 307 | | 19 | 35 | 56 | 36 | 58 | 59 | 64 | 33 | 93 | 61 | 35 | 69 | 52 | 36 | 0.4976 | FvH4_3g037 | | 213040 | 213392 | |
| 76 | GENE30724 | 1 | 4 | 9 | 9 | 1 | 5 | 8 | 2 | 6 | 6 | 9 | 1 | 0 | 7 | 1691 | 90 | Fvb3 | 9 | 8 | Inositol monophosphatase family protein |
| 307 | | 30. | 77. | 50. | 55. | 54. | 42. | 44. | 90. | 51. | | | 79. | 76. | 71. | 0.0035 | FvH4_3g037 | | 212394 | 212566 | |
| 77 | GENE30725 | 2 | 1 | 9 | 3 | 7 | 2 | 7 | 8 | 8 | 97 | 96 | 2 | 3 | 3 | 622 | 80 | Fvb3 | 1 | 8 | MYB-like 102 |
| 307 | | 74 | 19 | 19 | 23 | 16 | 18 | 18 | 16 | 21 | 18 | 18 | 17 | 19 | 17 | 0.5795 | FvH4_3g037 | | 211436 | 211823 | |
| 78 | GENE30726 | 8 | 47 | 65 | 16 | 74 | 00 | 29 | 76 | 73 | 64 | 27 | 82 | 90 | 56 | 4188 | 70 | Fvb3 | 8 | 7 | Protein kinase superfamily protein |
| 307 | | 82 | 11 | 12 | 19 | 16 | 19 | 18 | 15 | 17 | 18 | 92 | 12 | 16 | 10 | 0.6093 | FvH4_3g037 | | 210810 | 211435 | Zinc finger, RING-type;Transcription factor jumonji/aspartyl |
| 79 | GENE30727 | 9 | 63 | 79 | 32 | 03 | 13 | 95 | 23 | 06 | 36 | 3 | 08 | 18 | 21 | 3769 | 60 | Fvb3 | 4 | 0 | beta-hydroxylase |
| 307 | | 51. | 42. | 45. | 50. | | 57. | 42. | 64. | 34. | 29. | 41. | 27. | 30. | 56. | 0.3699 | FvH4_3g037 | | 209782 | 210094 | |
| 80 | GENE30728 | 1 | 2 | 4 | 4 | 35 | 4 | 4 | 1 | 1 | 1 | 5 | 8 | 3 | 6 | 2601 | 50 | Fvb3 | 4 | 1 | Major facilitator superfamily protein |
| 307 | | 12 | 10 | 88. | 10 | 66. | 10 | 64. | 67. | 47. | 46. | 79. | 81. | 38. | 43. | 0.0057 | FvH4_3g037 | | 209347 | 209678 | |
| 81 | GENE30729 | 7 | 5 | 5 | 8 | 4 | 8 | 6 | 7 | 1 | 9 | 4 | 1 | 7 | 3 | 6001 | 40 | Fvb3 | 7 | 4 | Major facilitator superfamily protein |
| 307 | | 16 | 91 | 75 | 48 | 10 | 66 | 77 | 13 | 33 | 80 | 10 | 19 | 12 | 98 | 0.0778 | FvH4_3g037 | | 208704 | 208991 | |
| 82 | GENE30730 | 5 | 5 | 8 | 7 | 02 | 6 | 2 | 43 | 9 | 3 | 66 | 21 | 60 | 5 | 611 | 30 | Fvb3 | 1 | 2 | Octicosapeptide/Phox/Bem1p family protein |
| 307 | | 17. | 20. | 23. | 21. | | 17. | 16. | 20. | 17. | 23. | 28. | | 21. | 18. | 0.1051 | FvH4_3g037 | | 207958 | 208168 | |
| 83 | GENE30731 | 4 | 2 | 9 | 2 | 17 | 2 | 6 | 4 | 4 | 6 | 6 | 28 | 1 | 7 | 1701 | 20 | Fvb3 | 2 | 4 | Cyclin D6;1 |
| 307 | | 18. | 17. | | 15. | 19. | 18. | 17. | 15. | 17. | 20. | 16. | 19. | 19. | 19. | 0.5368 | FvH4_3g037 | | 207551 | 207753 | |
| 84 | GENE30732 | 3 | 6 | 18 | 3 | 7 | 6 | 3 | 1 | 2 | 6 | 7 | 9 | 7 | 8 | 8068 | 10 | Fvb3 | 6 | 3 | Serine protease inhibitor (SERPIN) family protein |
| 307 | | 28. | 28. | 36. | 34. | | 27. | 26. | 22. | 31. | 21. | 40. | 37. | 27. | | 0.8795 | FvH4_3g037 | | 206740 | 206934 | |
| 86 | GENE30734 | 8 | 8 | 3 | 9 | 34 | 3 | 9 | 4 | 2 | 3 | 3 | 1 | 4 | 41 | 0521 | 00 | Fvb3 | 8 | 1 | Serine protease inhibitor (SERPIN) family protein |
| 307 | | 35. | 37. | 47. | 53. | 42. | 51. | 44. | 60. | 43. | 48. | 65. | 83. | 38. | 36. | 0.2205 | FvH4_3g036 | | 206379 | 206709 | |
| 87 | GENE30735 | 8 | 9 | 8 | 3 | 3 | 6 | 2 | 8 | 6 | 3 | 3 | 4 | 6 | 9 | 1578 | 90 | Fvb3 | 1 | 7 | Protein of unknown function (DUF594) |
| 307 | | 61 | 20 | 18 | 18 | 20 | 21 | 28 | 56. | 11 | 11 | 99. | 15 | 28 | 57. | 0.0550 | FvH4_3g036 | | 205947 | 206107 | |
| 88 | GENE30736 | 4 | 1 | 8 | 8 | 8 | 6 | 0 | 5 | 6 | 8 | 1 | 9 | 0 | 7 | 7754 | 80 | Fvb3 | 5 | 5 | myb domain protein 58 |
| 307 | | 31 | 21 | 25 | 19 | 23 | 25 | 18 | 14 | 24 | 24 | 25 | 27 | 19 | 17 | 0.5303 | FvH4_3g036 | | 205553 | 205809 | |
| 89 | GENE30737 | 59 | 60 | 36 | 46 | 62 | 56 | 71 | 72 | 52 | 95 | 92 | 51 | 95 | 13 | 2208 | 70 | Fvb3 | 6 | 2 | PLAC8 family protein |
| 307 | | 14 | 21 | 30 | 20 | 27 | 25 | 43 | 40 | 34 | 33 | 24 | 45 | 43 | 33 | 0.0408 | FvH4_3g036 | | 204735 | 205130 | |
| 91 | GENE30739 | 8 | 9 | 1 | 1 | 6 | 5 | 5 | 7 | 1 | 6 | 2 | 2 | 2 | 7 | 3962 | 50 | Fvb3 | 6 | 6 | Adenine nucleotide alpha hydrolases-like superfamily protein |
| 307 | | 73. | 25. | 18. | 31. | 19. | 26. | 24. | 16. | 16. | 19. | 15. | 17. | 20. | 16. | 0.1026 | FvH4_3g036 | | 204515 | 204740 | |
| 92 | GENE30740 | 2 | 2 | 6 | 7 | 5 | 3 | 5 | 4 | 6 | 2 | 3 | 8 | 3 | 5 | 8793 | 40 | Fvb3 | 4 | 0 | B-box type zinc finger family protein |
| 307 | | 14 | 10 | 13 | 13 | 11 | 13 | 11 | 14 | 12 | 12 | 88. | 11 | 10 | 80. | 0.1573 | FvH4_3g036 | | 204191 | 204347 | |
| 93 | GENE30741 | 1 | 6 | 8 | 7 | 4 | 3 | 2 | 2 | 2 | 0 | 3 | 7 | 8 | 9 | 244 | 30 | Fvb3 | 6 | 9 | AGAMOUS-like 24 |
| 307 | | 65. | 21 | 11 | 16 | 72. | 75. | 70. | 51. | 57. | 61. | 61. | 59. | 59. | 76. | 0.0681 | FvH4_3g036 | | 201205 | 201259 | |
| 95 | GENE30743 | 6 | 9 | 0 | 8 | 6 | 3 | 1 | 9 | 9 | 5 | 7 | 8 | 9 | 4 | 8159 | 00 | Fvb3 | 2 | 6 | Calcium-binding EF-hand family protein |
| 307 | | 10 | 24 | 24 | 21 | 19 | 30 | 20 | 23 | 15 | 18 | 24 | 20 | 24 | 19 | 0.7559 | FvH4_3g035 | | 200520 | 201012 | |
| 96 | GENE30744 | 38 | 57 | 78 | 35 | 35 | 75 | 21 | 25 | 40 | 87 | 08 | 05 | 30 | 44 | 7448 | 90 | Fvb3 | 5 | 0 | S-locus lectin protein kinase family protein |
| 307 | | 10 | 93. | 12 | 15 | 10 | 92. | 16 | 90. | 10 | 97. | 11 | 10 | 13 | 14 | 0.5494 | FvH4_3g035 | | 200205 | 200412 | |
| 97 | GENE30745 | 3 | 1 | 5 | 2 | 9 | 2 | 2 | 2 | 0 | 5 | 6 | 3 | 3 | 2 | 8072 | 81 | Fvb3 | 7 | 8 | S-domain-1 29 |
| 307 | | 44. | 53. | 47. | 41. | 51. | 50. | 55. | 36. | 56. | 39. | 49. | 60. | 45. | 46. | 0.7349 | FvH4_3g035 | | 199330 | 199437 | |
| 98 | GENE30746 | 1 | 6 | 5 | 9 | 6 | 7 | 1 | 3 | 9 | 7 | 2 | 7 | 7 | 7 | 7824 | 80 | Fvb3 | 5 | 8 | NAC domain containing protein 100 |
| 308 | | 23. | 25. | 29. | 29. | 31. | 24. | 26. | 26. | 26. | 25. | 27. | | 28. | 33. | 0.7767 | FvH4_3g035 | | 197734 | 198161 | |
| 00 | GENE30748 | 9 | 2 | 7 | 4 | 7 | 7 | 1 | 3 | 3 | 5 | 7 | 26 | 8 | 3 | 4908 | 60 | Fvb3 | 2 | 9 | S-domain-1 29 |

| | | | | | | | | | | | | | | | | | | | | | | |
|---|---|---|---|---|---|---|---|---|---|---|---|---|---|---|---|---|---|---|---|---|---|---|
| 308 | | | | 30. | 41. | 42. | 42. | 67. | 34. | 59. | 27. | 50. | 51. | 50. | 37. | 0.7343 | FvH4_3g035 | | 195730 | 195966 | NAC (No Apical Meristem) domain transcriptional regulator | |
| 01 | GENE30749 | 37 | 66 | 8 | 3 | 7 | 7 | 6 | 3 | 3 | 6 | 4 | 5 | 5 | 6 | 8589 | 40 | Fvb3 | 3 | 3 | superfamily protein | |
| 308 | | 41. | 14 | 12 | 14 | 41. | 43. | 40. | 13 | 11 | 13 | 27 | 35 | 96. | 14 | 0.0449 | FvH4_3g035 | | 195339 | 195595 | | |
| 02 | GENE30750 | 8 | 0 | 4 | 9 | 4 | 9 | 3 | 9 | 3 | 1 | 0 | 0 | 2 | 3 | 6318 | 30 | Fvb3 | 7 | 9 | floral meristem identity control protein LEAFY (LFY) | |
| 308 | | 15 | 28 | 33 | 44 | 38 | 18 | 61 | 34 | 80. | 26 | 27 | 35 | 94 | | 0.9663 | FvH4_3g035 | | 194770 | 195177 | | |
| 03 | GENE30751 | 6 | 2 | 9 | 2 | 0 | 2 | 3 | 9 | 9 | 5 | 9 | 3 | 2 | 85 | 0516 | 21 | Fvb3 | 3 | 8 | S-domain-1 29 | |
| 308 | | 44 | 50 | 49 | 77 | 58 | 37 | 73 | 51 | 23 | 40 | 53 | 53 | 14 | 13 | 0.9507 | FvH4_3g035 | | 193772 | 194139 | | |
| 04 | GENE30752 | 3 | 7 | 5 | 3 | 1 | 0 | 7 | 4 | 5 | 5 | 0 | 7 | 72 | 6 | 1241 | 20 | Fvb3 | 6 | 2 | S-domain-1 29 | |
| 337 | | 12 | 94 | 10 | 76 | 88 | 68 | 66 | 19 | 29 | 46 | 37. | 41 | 33 | 35. | 5.6077 | FvH4_3g030 | | 161598 | 161734 | | |
| 29 | GENE34030 | 62 | 2 | 09 | 5 | 8 | 1 | 5 | 9 | 4 | 6 | 2 | 1 | 3 | 2 | E-05 | 43 | Fvb3 | 2 | 9 | Tetratricopeptide repeat (TPR)-like superfamily protein | |
| 340 | | 43 | 67. | 68 | 10 | 11 | 24 | 28 | 44 | 89. | | 57. | 60. | 15 | 41. | 0.5098 | FvH4_3g042 | | 245107 | 245159 | | |
| 89 | GENE34397 | 38 | 5 | 1 | 2 | 6 | 39 | 86 | 36 | 1 | 74 | 2 | 7 | 97 | 2 | 4738 | 76 | Fvb3 | 6 | 6 | | 0 |
| 340 | | 27. | 30. | 49. | 46. | 48. | 51. | 49. | 33. | 41. | 45. | | 45. | 62. | 63. | 0.3828 | FvH4_3g042 | | 244550 | 244754 | | |
| 90 | GENE34398 | 5 | 2 | 8 | 5 | 9 | 6 | 6 | 1 | 1 | 5 | 49 | 3 | 4 | 6 | 2092 | 74 | Fvb3 | 3 | 4 | | 0 |
| 340 | | 12 | 12 | 14 | 20 | 11 | 99. | 14 | 12 | 91. | 14 | 10 | | 16 | 82. | 0.2495 | FvH4_3g033 | | 181580 | 181687 | | |
| 96 | GENE34404 | 0 | 9 | 3 | 3 | 7 | 2 | 6 | 4 | 2 | 5 | 7 | 94 | 8 | 1 | 7179 | 40 | Fvb3 | 9 | 2 | S-locus lectin protein kinase family protein | |
